# The non-stationary dynamics of fitness distributions: asexual model with epistasis and standing variation

**DOI:** 10.1101/079368

**Authors:** Guillaume Martin, Lionel Roques

**Affiliations:** ISEM, UMR CNRS-UM II 5554, Université Montpellier II, Montpellier Cedex 5, France; BioSP, INRA, 84914, Avignon, France

**Keywords:** asexual evolution, clonal interference, transient dynamics, mutation-selection balance, Fisher’s geometrical model

## Abstract

Various models describe asexual evolution by mutation, selection and drift. Some focus directly on fitness, typically modelling drift but ignoring or simplifying both epistasis and the distribution of mutation effects (travelling wave models). Others follow the dynamics of quantitative traits determining fitness (Fisher’s geometrical model), imposing a complex but fixed form of mutation effects and epistasis, and often ignoring drift. In all cases, predictions are typically obtained in high or low mutation rate limits and for long-term stationary regimes, thus loosing information on transient behaviors and the effect of initial conditions. Here, we connect fitness-based and trait-based models into a single framework, and seek explicit solutions even away from stationarity. The expected fitness distribution is followed over time via its cumulant generating function, using a deterministic approximation that neglects drift. In several cases, explicit trajectories for the full fitness distribution are obtained, for arbitrary mutation rates and standing variance. For non-epistatic mutation, especially with beneficial mutations, this approximation fails over the long term but captures the early dynamics, thus complementing stationary stochastic predictions. The approximation also handles several diminishing return epistasis models (e.g. with an optimal genotype): it can then apply at and away from equilibrium. General results arise at equilibrium, where fitness distributions display a ‘phase transition’ with mutation rate. Beyond this phase transition, in Fisher’s geometrical model, the full trajectory of fitness and trait distributions takes simple form, robust to details of the mutant phenotype distribution. Analytical arguments are explored for why and when the deterministic approximation applies.

**Significance statement:** How fast do asexuals evolve in new environments? Asexual fitness dynamics are well documented empirically. Various corresponding theories exist, to which they may be compared, but most typically describe stationary regimes, thus losing information on the shorter timescale of experiments, and on the impact of the initial conditions set by the experimenter. Here, a general deterministic approximation is proposed that encompasses many previous models as subcases, and shows surprising accuracy when compared to stochastic simulations. It can yield predictions over both short and long timescales, hopefully fostering the quantitative test of alternative models, using data from experimental evolution in asexuals.

## Introduction

Empirical dynamics of fitness in simple environments are still not quantitatively predicted by evolutionary biology, in spite of a wealth of theoretical progress and an ever-growing corpus of data produced by experimental evolution. To our knowledge, no model exists that was parameterized from independent data, and then has proved to predict observed fitness trajectories; in either sexual or asexual organisms, from de novo mutations or preexisting standing variance. Patterns of fitness trajectories in microbes (*de novo* mutations in asexuals) have been confronted to and fitted with various theoretical predictions, showing qualitative agreement with models of clonal interference (Tsimring *et al.* 1996; Miralles *et al.* 2000; Gerrish 2001; Desai *et al.* 2007), and suggesting pervasive diminishing return epistasis among beneficial mutations (Chou *et al.* 2011; Khan *et al.* 2011). However, *fitting* is not *predicting*: several alternative models can be qualitatively consistent with the same dataset (Frank 2014). Regarding fitness dynamics during adaptation from standing variance, both theory and data are relatively scarce, at least in asexuals; this limits our knowledge of the transient effects of standing variance, while these can be critical for short-term adaptive responses to environmental challenges.

Important progress has been made, over several decades, with a rich variety of models predicting fitness dynamics. These models critically depend on (i) a mutation rate and (ii) a distribution of fitness effects of mutations (DFE), which is either independent of the background genotype (no epistasis for fitness), or depends on it, minimally on its fitness. They differ in the genotype-fitness landscape considered and the regimes assumed to derive the evolutionary dynamics. Models of mutation and selection in asexuals roughly fall into two (seemingly disconnected) classes: DFE-based models that directly track the distribution of fitness and trait-based models that follow the distribution of underlying quantitative traits, which determine fitness. The aim of this work is to handle this variety of models into a single analytical framework (in terms of partial differential equations, or PDE), and to use it to derive new results for these models, regarding non-stationary dynamics or equilibria. We start by briefly summarizing these existing approaches, in a necessarily far from exhaustive manner.

**Fitness-based models** directly follow the dynamics of fitness distributions, typically with a constant mutation rate and DFE over time (no epistasis). Initially based on deterministic equations and diffusive mutation effects (Tsimring *et al.* 1996), they were then refined to include stochasticity and more general DFEs of purely beneficial mutations (Gerrish and Lenski 1998; Rouzine *et al.* 2003; Dwyer 2012; Good *et al.* 2012). More recently, the interplay of a distribution of deleterious and beneficial mutations has been studied in this context, in either low (e.g. Good and Desai 2014) or high (e.g. Neher and Hallatschek 2013) mutation rate limits. As beneficial mutation influx becomes large in asexuals, co-segregating lineages compete for fixation and slow down adaptation, a process further affected by the deleterious mutations that accumulate on each lineage. These ‘clonal interference’ dynamics, in the presence of stochastic fluctuations, are difficult to analyze and often yield complex or non-explicit formulae, but several models have provided important insight into this process. They have been handled through alternative modelling approaches, accurate in different regimes: low to intermediate mutation rate for the original clonal interference models (Gerrish and Lenski 1998; Gerrish 2001), or higher mutation rate for the more recent ‘travelling wave’ models (Rouzine *et al.* 2003; Good *et al.* 2012; Neher and Hallatschek 2013). Note that in the limit of very large populations, high mutation rates and weak mutation effects, a simple and explicit Gaussian travelling wave is retrieved for the expected fitness distribution (Neher and Hallatschek 2013).

This rich literature, reviewed elsewhere (e.g. Rouzine *et al.* 2003; Desai and Fisher 2007; Sniegowski and Gerrish 2010; Desai 2013), has a common feature: it describes the stationary regime of a stochastic process. This implies that a full trajectory from given initial conditions (possibly with standing variance) is not available, only the ultimate average rate of steady fitness change. Furthermore, as time goes on, the envelope around this mean fitness prediction typically explodes so that individual populations may lie far from the predicted mean at any time. This limits the comparison to empirical trajectories, which typically start away from stationary regime, and contain a few replicates. Note however, that this assumption of steady increase in fitness is often envisioned as reflecting a constant struggle between a steadily changing environment and an adapting population (Neher and Hallatschek 2013). It is possible that in such regime the envelope may remain narrow and steady-state may be reached faster.

Another aspect of the approach is that epistasis must be ignored here; otherwise mutation rates and effects may change over time (as the dominant backgrounds change), impeding the setting of a stationary regime. Recent extensions do include some form of epistasis or deleterious mutations (Kryazhimskiy *et al.* 2009; Dwyer 2012; Good and Desai 2015). However, analytical progress is then difficult beyond the master equation: relatively simple exemplary cases were analyzed in depth but always in regimes where clonal interference is negligible. Note also that other DFE-based models were devoted to describe mutation-selection balance (another stationarity assumption), ignoring drift and epistasis. General insight into equilibrium fitness distributions has been gained from quasi-species theory (Eigen 1971) or asexual mutation-selection-balance models (Johnson 1999). This literature will not be reviewed here either (see Wilke 2005), but in general analytical progress has often proved difficult unless simplified forms of DFE are assumed (discussed in Martin and Gandon 2010).

**Trait-based models** form an equally central body of literature that deals with adaptation affecting a trait or set of traits under selection for an optimum (via some concave phenotype-fitness function). These single peak trait-based models date back to Fisher’s (Fisher 1930) geometrical model (FGM), and also produced a rich literature connected to evolutionary quantitative genetics (Lande 1979). This approach is constrained into a particular form of DFE, but one that does include (i) pervasive epistasis and dominance, and (ii) both beneficial and deleterious mutations. Several patterns of mutant fitness expected in the FGM have been tested on fitness data from mutant lines (Martin *et al.* 2007; Trindade *et al.* 2010; Manna *et al.* 2011; Sousa *et al.* 2011; Trindade *et al.* 2012; Hietpas *et al.* 2013), showing promising overall agreement. The FGM also emerges as the limit of a broader class of genotype-phenotype-fitness landscapes involving highly integrated “small–world” phenotypic networks (Martin 2014). Overall, the FGM seems a reasonable null model for evolutionary predictions (reviewed in Tenaillon 2014). The population genetics of adaptation by mutation and selection, in such trait-based models, has also seen many developments, reviewed extensively elsewhere (e.g. Burger 2000; Orr 2005). It provides a well-studied theory for equilibrium states in various situations (detailed in Roze and Blanckaert 2014); several qualitative properties of equilibria have even been obtained for more general trait-fitness relationships, at least with a single trait (detailed in Burger 1998; Burger 2000). The effect of standing genetic variance has also been treated extensively (from its quantitative genetics heritage), making the FGM an interesting complement to DFE-based models. Furthermore, predictions on trait distributions can be transformed into predictions on measurable fitness distributions under the model (e.g. Martin and Gandon 2010). Yet, in spite of interest in its potential (Barton 1998; Gordo and Campos 2012), analytic progress in situations relevant for experimental evolution (notably asexuals), has proven equally difficult to obtain. Even equilibrium states are not fully resolved in the FGM. Alternative analytic approximations only exist at each extreme of the mutation rate spectrum: House of Cards for a single trait (Turelli 1984) vs. Gaussian for arbitrarily many traits (Kimura 1965; Lande 1980), respectively, in the low vs. large mutation rate limits. When dealing with the dynamics of adaptation, the classic approach (Lande 1979) focuses on large highly polymorphic sexual populations, where the genetic variance of the traits is transiently approximately constant: another stationarity assumption, valid this time over finite timescales. However, this option breaks down with asexuals, in general. Alternatively, stochastic models of mutation-selection-drift dynamics have been implemented under the FGM for adaptation trajectories (Orr 2000), or mutation-selection-drift balance (Tenaillon *et al.* 2007). However, they apply in a weak mutation strong selection limit (or with unlinked non-epistatic loci in sexuals) where clonal interference is negligible. Finally, it is noteworthy that treatments of trait-based models with high mutational input (Gaussian theories) put less emphasis on drift (often neglected), than their fitness-based counterparts. They do involve multiple co-segregating mutants (clonal interference), but the deterministic predictions prove fairly accurate in this case, suggesting that some difference in the assumptions makes the interplay of drift and other forces less critical.

### Aim of this work

Overall, we enjoy a wealth of alternative, complementary approaches of adaptive (or maladaptive) fitness dynamics in the presence of mutation, selection and possibly drift. Yet, they are not easily connected together. They do not provide a readily testable prediction, in terms of trajectories of fitness distributions over time, from known initial conditions, in the large asexual populations typical of evolution experiments. To derive such predictions, we extend an approach initially proposed by R. Bürger (1991), who studied trait-based models via the dynamics of the cumulants of the trait distribution, under selection and non-epistatic mutation. We apply this framework to fitness itself. Deterministic dynamics of fitness cumulants/moments have been used previously in non-epistatic fitness-based models: either neglecting drift (Johnson 1999; Desai and Fisher 2011; Gerrish and Sniegowski 2012) or including a stochastic diffusion component and considering the expected cumulants over replicates (Rattray and Shapiro 2001; Good and Desai 2013). Following Bürger’s (1991) strategy, these studies solved a finite set of cumulant equations numerically, but the system could not be closed, as cumulants/moments influence each other in cascade. Here, we focus on the moment and cumulant generating function (MGF and CGF, respectively) of the fitness distribution, which handles all moments (resp. cumulants) in a single function. In a variety of models, this allows to ‘close the system’ into a single partial differential equation (PDE) describing the dynamics of the expectation of the fitness distribution, among stochastic replicates, by ignoring the effect of drift. We further include mutational epistasis by considering DFEs that broadly depend on background fitness. Overall, several processes are jointly handled by the PDE (**Fig. 1**): starting from an arbitrary initial fitness distribution, new mutations accumulate on each lineage (with lineage-dependent DFE), which co-segregate under selection (clonal interference). In several classes of models, explicit solutions can be found for the PDE, providing a fully analytic theory in terms of mutational parameters and standing variance. We check the predictions against stochastic individual based simulations of various subcases.

**Fig. 1.**
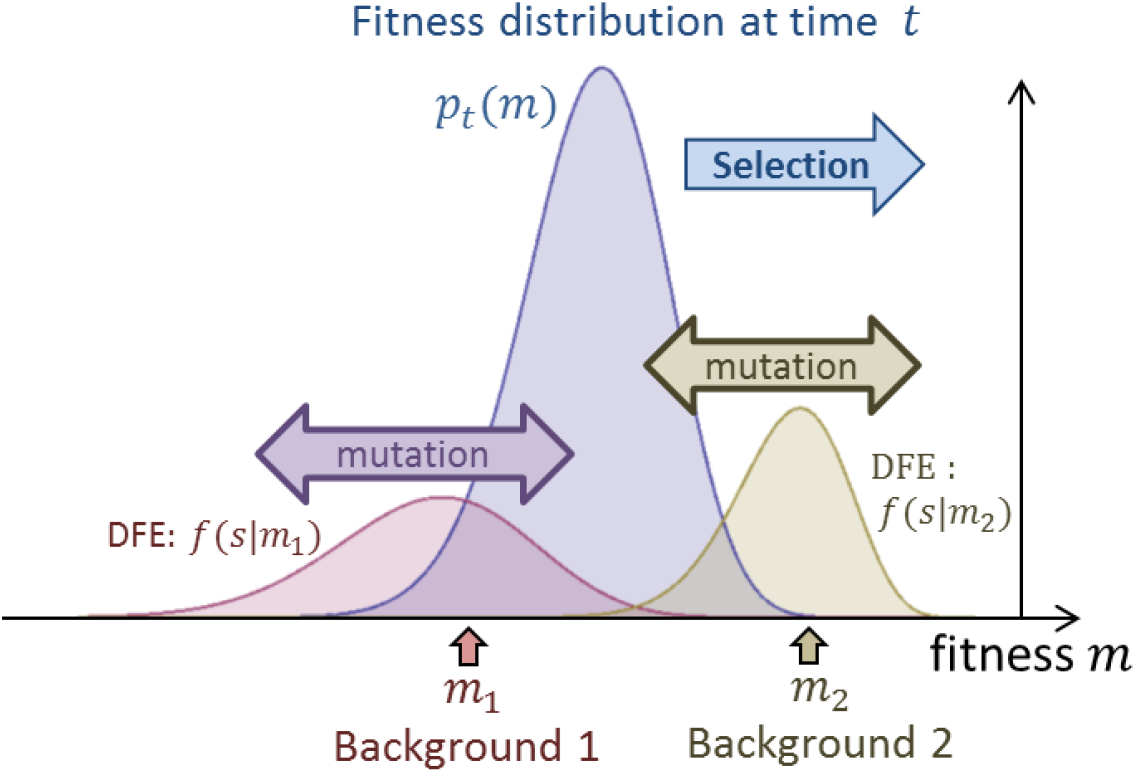
The standing fitness distribution (*p*_*t*_ (*m*), blue curve) travels to the right by selection. Each genetic background under this distribution (e.g. *m*_1_ and *m*_2_ here) mutates to new genotypes with fitness *m*_*i*_ *+ s* where *s* has the density *f*(*s*|*m*_*i*_) depending on background fitness (red and brown curves for *m*_1_ and *m*_*2*_, respectively).

### Heuristic statements

Before describing the model in more mathematical detail, we first tackle some qualitative aspects of fitness dynamics in the different models above. Let us start by a somewhat technical remark that justifies the use of generating functions here. With any model where the DFE only depends on parental fitness and in an asexual (no recombination/segregation), fitness is the only ‘trait’ which distribution fully determines its own evolution. We can thus follow this distribution alone, ignoring the genetic or phenotypic details underlying its variation, namely the number and effects of the mutations carried by different genotypes, over their entire genome. This does not preclude the complications described above: multiple mutations accumulate on each lineage, multiple lineages cosegregate and compete for ultimate fixation and each lineage may have its own background-dependent DFE (epistasis), as long as this dependence is entirely mediated by the background fitness. Generating functions handle sums of independent variables in a convenient manner, which helps study the cumulative effect of multiple mutations accumulating in lineages. It is also known that the effect of selection on fitness distributions takes simple form in terms of generating functions (Hansen 1992; Manna *et al.* 2012).

Second, let us consider why and when drift may be ignored in a given finite population, or among replicate finite populations, to describe the average fitness trajectory. The primary impact of drift identified in stochastic fitness-based models lies in its impact on the very fittest edge of the fitness distribution. When this edge represents a small absolute number of individuals, stochastic fluctuations in this subpopulation indirectly bias the future mean fitness dynamics of the whole population, over longer times. This effect does not average out if we consider the average mean fitness of replicate populations. However, over a substantial initial period, this fitter edge has little influence on the mean fitness dynamics (discussed in Gerrish and Sniegowski 2012), for two reasons. First, in a large polymorphic population, the short term mean fitness dynamics are driven by selection and mutation in the bulk of the population, which behaves roughly deterministically. Second, even in a smaller population, drift, of itself, only slightly alters the average frequency dynamics of genotypes: roughly by an order 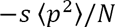 where *N* is population size and *p* and *s* are the allele’s frequency and fitness effect, respectively (see, e.g. Otto and Barton 2001). Therefore, any quantity that is linear in genotype frequencies, such as mean fitness or the moment generating function of the fitness distribution, is only slightly affected by drift over this timescale. It is only once new mutants establish (or not) that the future of the fitness dynamics is inaccurately predicted by a deterministic model: ignoring the stochastic loss of these fitter genotypes leads to overestimate mean fitness over longer timescales. Finally, even over longer timescales, the bias induced by drift is only visible if it accumulates over time, as the fittest edge stochastically moves towards fitter classes (at a speed overestimated by the deterministic model). If the set of all possible fitnesses is bounded by some maximal value, stochastic fluctuations should become less important, as the edge cannot spread forward forever: the delay between the edge and the bulk is bounded, and tends to decay over time (as the bulk adapts). Most trait-based models consider adaptation towards a phenotypic optimum, implying a form of diminishing returns epistasis, where fitness is bounded on the right by the fitness of this optimum. This may explain why the mean fitness dynamics in these models has been accurately captured by deterministic theories. The same applies for purely deleterious models, where fitness cannot travel beyond the unloaded fitness class. In this case, however, loss of the fitter class also happens and affects the long-term dynamics (Muller’s ratchet 1932). Yet, this happens over much longer timescales, as the edge is a large subpopulation and as each ‘click’ of the ratchet has a small impact (especially with continuous DFEs, where the new fittest class typically lies close to the previous one). This argument suggests that, in the presence of a fitness upper bound, it may be possible to accurately capture fitness dynamics by a mere deterministic model, even if clonal interference is involved and even over long timescales. It also suggests that non-epistatic models with beneficial mutations (where deterministic models fail in the long run) could still show transient fitness dynamics which average (over replicates) is captured by a deterministic model. Deriving such predictions (and justifying the above heuristic), as well as testing their accuracy with stochastic simulations is the central aim of this article.

## Model

### General setting

We assume finite haploid asexual populations and follow the expected fitness distribution among replicates, started from the same initial fitness distribution. We consider a continuous time model (overlapping generations), measured in arbitrary units (hours, days etc.). This setting can also approximate a discrete time model (non-overlapping generations) when effects are small per generations, the time *t* is then measured in generations: this will actually be our simulation scheme. We follow the dynamics of the distribution of the Malthusian fitness *m* (hereafter ‘fitness’). In continuous time, this is the expected exponential growth rate of a given genotype. In a discrete time approximation, *m* is the log of the Darwinian fitness (*m* = log*W*), namely the log of the expected geometric growth rate of a genotype. We define fitness relative to a reference, set at *m =* 0, without loss of generality. This reference is arbitrary as we consider evolutionary dynamics (relative fitness) without coupling to demography. In those models that include some fitness upper bound (e.g. single peak landscape models or models with only deleterious mutations), we set the optimal genotype (with fitness equal to this maximum) to be the reference *m* = 0 for convenience (so that all *m* ≤ 0). In other models (e.g. models with context-independent beneficial mutations), the reference is just an arbitrary point in fitness space. At any time *t*, an arbitrary set of *K*_*t*_ genotypes, with constant fitnesses 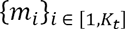, coexist in relative frequencies 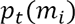, satisfying 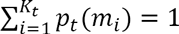. The approach can describe discrete classes (*K*_*t*_ finite) or infinite countable classes in the limit 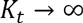 (with convergence to a continuous distribution of fitness). Genotypes compete by frequency-independent selection, and mutate according to a Poisson process with fixed rate *U* per capita per unit time. The fitness of a mutant which parent has fitness *m* is *m + s*, where *s* is the selection coefficient of the mutation relative to the parent, and is drawn from an arbitrary distribution with probability distribution function *f*(*s*|*m*) (pdf; a probability density function if the distribution is continuous) depending on the parent fitness *m*.

### Notations

We must define various expectations and means. We use an overbar 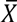 to describe any variable *X*(*m*), averaged over the current distribution of genotypes within a focal population: 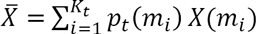. We define the expectation *E*(*Y*|*m*) of any variable Y(*s*) over the DFE in background 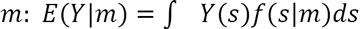, and we denote by 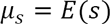 the mean DFE whenever it does not depend on *m.*

### Generating functions

The distribution of *m* at time *t* can be characterized by its moment generating function (MGF): 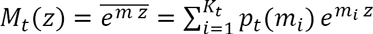. For any finite population (*K*_*t*_ finite) this MGF is always defined over the full line 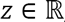, but we may study it on a compact subset spanning 0 (here, 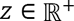), without loss of generality: this helps handle several continuous class limits (when 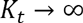). This generating function provides essential information on the distribution at time *t:* its derivatives at *z* = 0 are the raw moments of the fitness distribution, notably the mean fitness 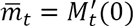 (the prime refers to differentiation with respect to *z*). For mathematical convenience, we mostly focus on the natural logarithm of the moment generating function, which is the cumulant generating function (CGF): *C*_*t*_(*z*) = log *M*_*t*_(*z*). Its derivatives at *z* = 0 are the cumulants of the distribution: in particular, the first three derivatives are the mean 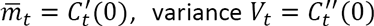 and third central moment (related to skewness) 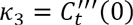. Additionally, the maximum of the distribution is given by 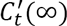 and the weight of the class *m=* 0 is given by 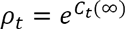; we say that the distribution has a *spike* at *m=* 0 when this quantity *ρ*_*t*_ is strictly positive. It should also be noted that the full distribution of *m* at time *t* can be retrieved by applying an inverse Laplace transform to 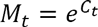.

Because each replicate population has its own trajectory of genotypic frequencies, the generating functions 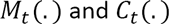 are stochastic functions of *z* over time. We seek to predict the behavior of the *expectation* of such variables over stochastic replicates, so we use 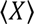 to denote any such expectation of *X.* In particular 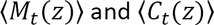 are the expected MGF and CGF, which are deterministic functions of *z* and *t*, while 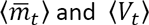 are the expected mean fitness and variance in fitness within populations. These are deterministic functions of time.

### Organization of the article

In Appendix A, we derive *exact* dynamics for the expected generating functions, which do not close. Then we describe *approximate* closed dynamics for these quantities under a deterministic approximation ignoring drift. In Appendix B, we derive general properties of the approximate dynamics, and Appendices C,D,E provide detailed applications to particular classes of mutational models. In the ‘Model’ section below, we summarize our results on the expected CGF 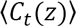, and its approximate deterministic counterpart, denoted 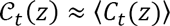 (the ≈ sign is a reminder that the result is approximate). The ‘Application’ section then illustrates applications to several classes of mutation models, evaluating the accuracy of the approximation on stochastic simulations. A last section summarizes some analytic results on the error involved by the approximation, and hints on why and when it applies. All notations are summarized in **Table 1**.

**Table 1.**
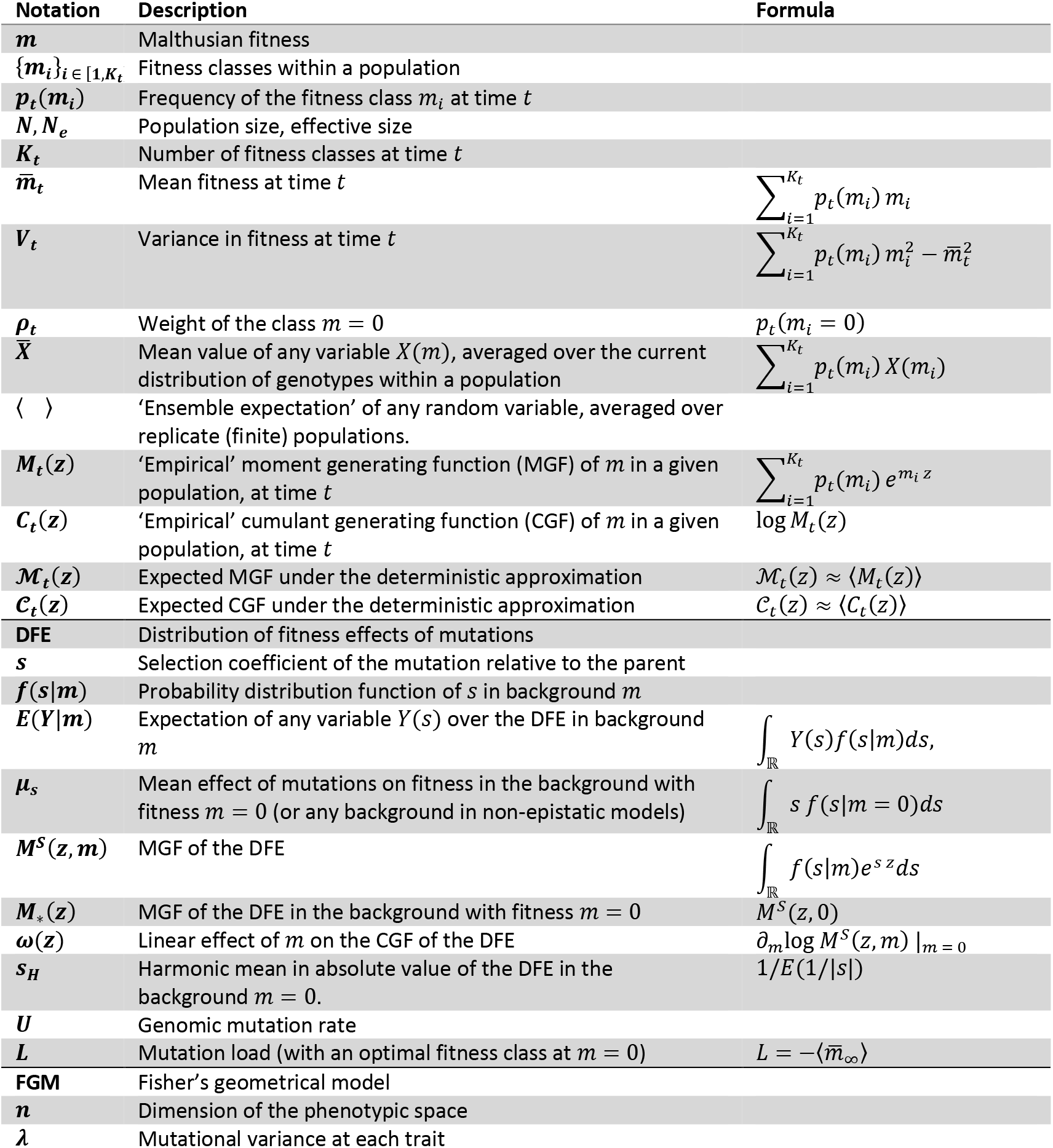
Main notations used throughout the article.

### Dynamics of the expected CGF under selection, drift and mutation

Using a multi-type Wright-Fisher diffusion approximation to genotype frequency dynamics (Section II in Appendix A), it can be shown that the change by selection and drift (‘SD’), over ∆*t*, in the expected CGF 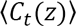 satisfies

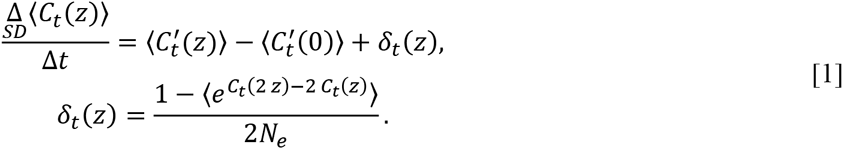

Here, *δ*_*t*_(*z*) is the contribution generated by drift (it vanishes if *N_e_ →* ∞), essentially the same as given in Good & Desai’s (2013) eq. (D.4). This dynamic term does not allow to close the system as *δ*_*t*_(*z*) does not depend directly on 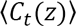. We thus rely on a deterministic approximation (that we will use all along), which simply ignores *δ*_*t*_(*z*) in the dynamics, yielding an approximate expected CGF 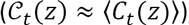, with closed dynamics 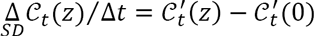.

Mutation (see the General setting section above) generates a *distribution of fitness effects* (DFE) which MGF is denoted 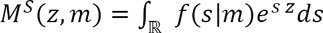. It is assumed to have known analytical form, over some positive domain 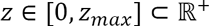, determined by the model considered. This may include continuous or discrete distributions, but it does require that the DFE have finite higher moments (so that an MGF can be analytically defined). The change in 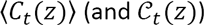 by mutation (‘mut’), over ∆*t*, takes the general form (Section III.1 in Appendix A):

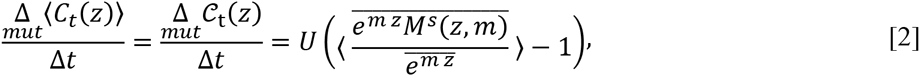

where we recall that 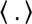 is the expectation over replicate populations, while the overbar refers to the averaging with respect to *m*, within a given population, at current time *t*. The limit, as ∆*t* → 0, of 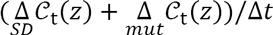 from Eqs. **[1]** and **[2]**, yields the continuous time dynamics of the expected CGF, under the deterministic approximation:

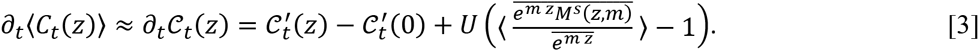

This is our central result, from which all following dynamics are derived. In general, the mutation kernel in Eq. **[3]** does not generate a closed system, even under the deterministic approximation, as the mutational term cannot be expressed in terms of 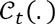. Fortunately, this term simplifies in several general classes of models, which we detail below, summarize in **Table 2** and implement in **Supplementary material 2** (see below).

**Table 2.**
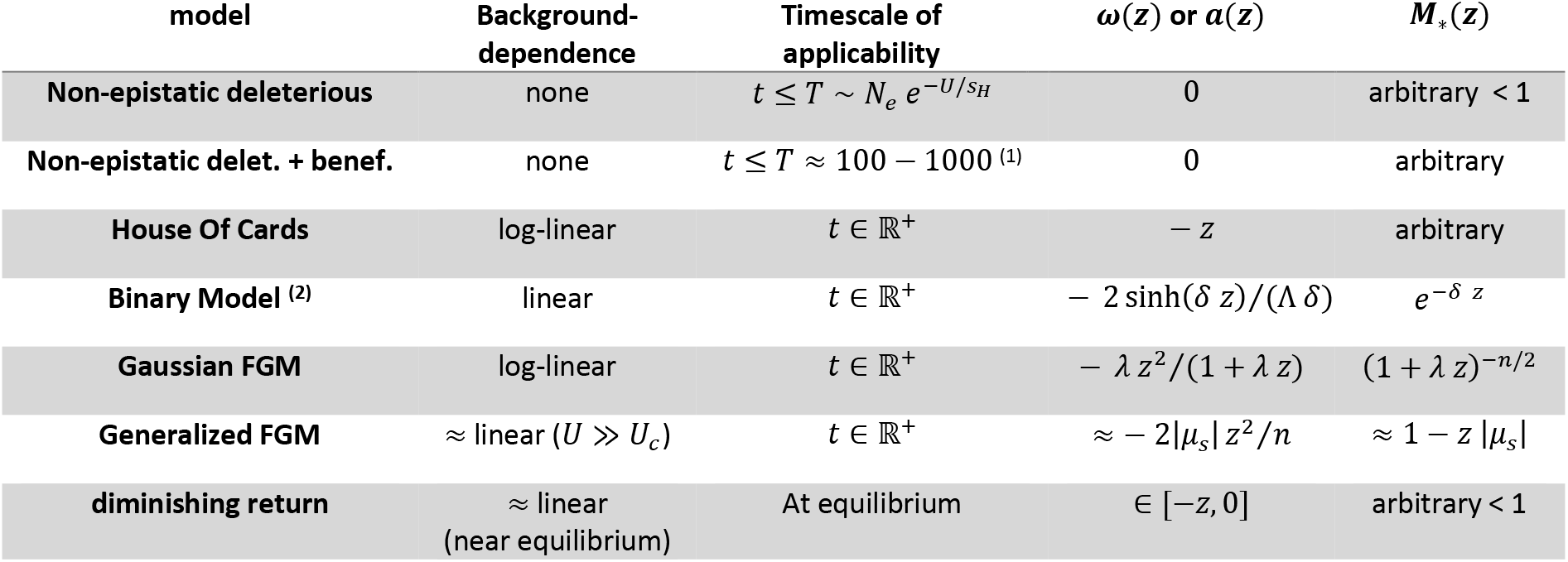
Various mutational models handled by the proposed framework. These models only apply when 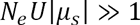. For each model, each column gives (i) the model type, (ii) the type of background dependence, (iii) the timescale (sometimes approximate) over which the prediction applies (in that it is expected to be reasonably to very accurate), (iv) the background dependence function (*ω*(*z*) for log-linear background-dependence or *α*(*z*) for linear background-dependence), and (v) the MGF *M*_*_(*z*) of the DFE in the background with fitness *m=* 0 (fittest background in models with a maximum fitness). In some models the ′ ≈ ′ notifies that this is an approximate result or a conjecture; 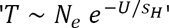 means that the two quantities have the same order of magnitude. ^(1)^: conjecture and timescale based on observations in our simulations. ^(2)^: simplified version of Rouzine et al.’s (2003) model, detailed in Appendix A III.2. Here, Λ is the number of sites (′*L* in the original paper) and *δ* is the constant deleterious effect of ‘mutant’ alleles (′*s′* in the original paper).

### Linear background-dependence

A first important situation is when the MGF of the DFE can be (exactly or approximately) written as a linear function of 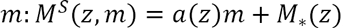, with some function *a* and with 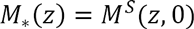 being the MGF of the DFE in the background with fitness *m=* 0. As an MGF, *M*_*_ is continuous on a domain including 0 and must satisfy 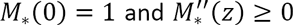. The function *a* must satisfy *a*(0) = 0 and either 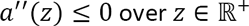, if fitnesses are bounded on the right so that all 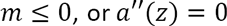 if fitnesses are unbounded on the right. This is required for *M*^*S*^ to satisfy the basic MGF properties *M*^*S*^(0,*m*) = 1 (conservation of probability) and 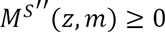 (convexity for all *m* and *z*). Linear background-dependence (see Section III.2 in Appendix A), implies a mutation kernel (Eq. **[2]**) of the form 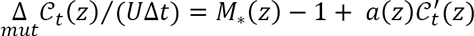. The (approximate) expected CGF 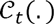 then satisfies a 1^st^ order linear nonlocal PDE:

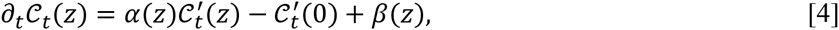

where the functional coefficient are *α(z*) = 1 + *U a(z*) and *β*(*z*)= *U*(*M*_*\**_(*z*) – 1), with *α*(0) = 1 and *β*(0) = 0. This PDE has the boundary condition 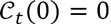, and initial condition 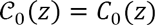 (initial fitness distribution); it can be solved analytically (Section II.1 in Appendix B). Define the function *y*, solution of the ODE *y*′(*z*) = *α*(*y*(*z*)) with initial condition *y*(0) = 0 and its functional inverse 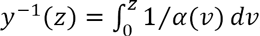, such that 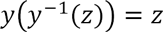, defined on [0, *z*_1_), where *z*_1_ is the first positive root of *α.* The unique solution of Eq. **[4]** from initial condition *C*_0_(*z*) is

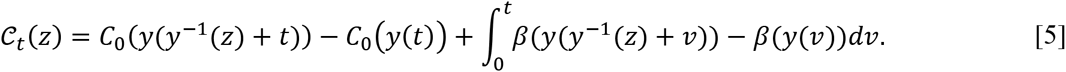

The corresponding trajectory of the expected mean fitness is (under the deterministic approximation)

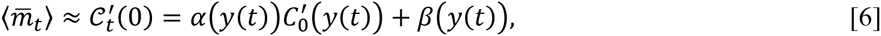

for all *t* ≥ 0. A similar explicit expression is given in Appendix B (Eq. B31) for the trajectory of the expected variance 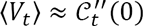. More generally, Eq. **[5]** gives the trajectory of the whole fitness distribution, for several classes of models described in the Application section.

### Examples of linear background-dependence models

#### Non-epistatic models

An obvious case of linear background-dependence is for any non-epistatic model (which DFE has finite moments, so that its MGF exists). In these, we have 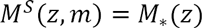 for all backgrounds, so that *a*(*z*) *=* 0 and *α*(*z*) = 1.

#### Simplified version of Rouzine et al.’s (2003) ‘Binary model’

In this model (detailed in Appendix A III.2), genotypes consist of Λ bins representing sites (we use notations different from the original article to avoid confusions with other quantities in this article). Each bin codes for a wild-type (‘0’) or mutant (‘1’) allele (with constant deleterious effect –*δ <* 0). Mutation, at rate *u* per site (genomic rate *U = u* Λ), randomly creates shifts between allele states and allelic effects add-up across the genome. This model shows mutational epistasis (the DFE depends on the background *m*), although fitness is still a sum of allelic effects over the genome. It also implies an upper bound *m=* 0 to all possible fitnesses (i.e. the unloaded wild-type with only ‘0’ bins) and has linear background-dependence (see Eq. **(A10)**, Appendix A and **Table 2**). It can be checked that 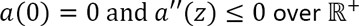. We do not explore this model further here, except in **Supplementary material 2** (see below).

### Log-linear background-dependence

Alternatively, the MGF of the DFE may be log-linear in 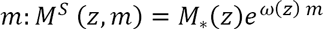. Here again, 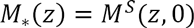 is convex and satisfies *M*_*_(0) = 1, while *ω* must be concave (with bounded fitness set *m* ≤ 0) and *ω*(0) = 0. Plugging this form into the mutational kernel in Eq. **[2]** yields another nonlocal 1^st^ order PDE for the (approximate) expected CGF, but this time it is nonlinear (Section III.4 in Appendix A):

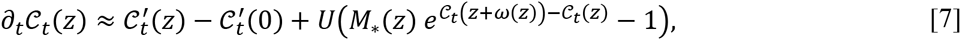

for *t* ≥ 0 and *z* ≥ 0, with the boundary condition 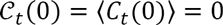. The second term 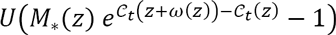 in Eq. **[7]** describes the effect of mutations accumulating on each background, with a dependence on the standing distribution of background fitnesses (on 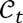) mediated by *ω*(*z*). Note that this time, this term is only approximate, under similar conditions as the deterministic approximation used all along (detailed in III.3 of Appendix A).

The well-posedness of Eq. **[7]** requires that 0 ≤ *z* + *ω*(*z*) so that the nonlocal term remains within the domain under study. It is the case for any epistatic model 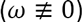 showing log-linear background-dependence, with a fitness optimum at *m=* 0 (see Section I.1 in Appendix B). Although we were not able to get an explicit solution of Eq. **[7]**, which is a nonstandard PDE problem due to the two nonlocal terms 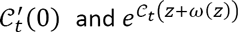, we were able to get some insight into the behavior of the solution. First, 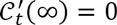 for all positive times (Section I.2 in Appendix B), with epistatic model 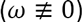. This means that the support of the fitness distribution instantaneously reaches the optimum *m=* 0, whatever the initial fitness distribution. It implies a memoryless property in the sense that the long-time behavior of the solution is not impacted by the initial fitness distribution, which is not obtained in non-epistatic models (*ω*= 0). Second, analytical expressions are derived (Section I.3 of Appendix B) for the *k*^*th*^ cumulants of the equilibrium distribution (*k* ≥ 0) and a dichotomy for the value of the equilibrium mean fitness; namely, either 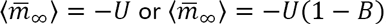, for some positive constant *B.* Third, the existence of a spike implies that 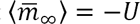 (Section I.4 of Appendix B). These results were obtained under any of the two general properties (Section I.2 in Appendix B): **(H)** any background can mutate to the optimal background; or **(H')** any background can at best mutate to some fitter but suboptimal class. Biologically, this simply means that some form of compensation of deleterious mutations exist.

### Examples of log-linear background dependent models

As an example, we describe two classic models of context-dependent DFEs where log-linear background dependence applies (see also **Table 2**).

#### Fisher’s (1930) geometrical model (FGM)

This model assumes that each genotype is characterized by a (breeding value for) phenotype at *n* traits 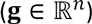 (possibly with some environmental variance effects). An optimal phenotype corresponds to maximal fitness and sets the origin of phenotype space (**g = 0**). Fitness decreases away from this optimum, and mutation creates random iid variation **dg** around the parent, for each trait. In all our examples, we will consider a quadratic fitness function: in continuous time models, Malthusian fitness is a quadratic function of the breeding value 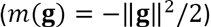, and in discrete time versions, Darwinian fitness is a Gaussian function of 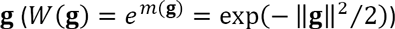. A classic version of this model is the ‘Gaussian FGM’, where mutation phenotypic effects are multivariate normal: 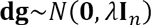, where *λ* > 0 is the mutational variance at each trait, and **I**_*n*_ is the identity matrix in *n* dimensions. This ‘Gaussian FGM’ is also the standard model of evolutionary quantitative genetics, dating back to Kimura’s (1965) and Lande’s (1980) work on mutation and selection on traits with a complex genetic basis (infinitely many possible alleles). The Gaussian FGM shows exact log-linear context-dependence (Martin 2014): 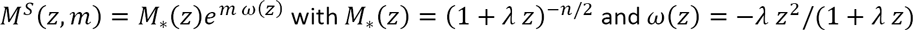. We study this model in depth in the Application section.

#### Kingman’s (1978) House of Cards (HOC) model

this model assumes that mutants have absolute fitness that follows a unique distribution, independently of the background in which they arise. This model is epistatic in that the DFE depends on the background: 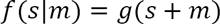 so that mutant absolute fitnesses *X* have a given fixed fitness distribution with pdf *g*(*x*). Versions of the HOC were used e.g. in (Kryazhimskiy *et al.* 2009) and (Mccandlish *et al.* 2014), respectively with an exponential or Gaussian distribution *g*, and focusing on a regime of low *NU* where substitutions occur sequentially (no clonal interference). In this model, the MGF of the DFE is 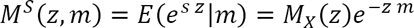 where 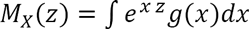 is the MGF of the chosen distribution of *X* with pdf *g.* Thus this model, in its general version, implies log-linear background dependence with 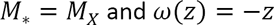. We do not explore this model further here, except in **Supplementary material 2** (see below).

### Individual based simulations

Individual based, discrete time simulations were used to check the validity of the approximations in finite populations, for various mutational models. Individuals were sampled every generation according to their fitness *W* = *e*^*m*^ (Wright Fisher model of genetic drift and selection). Mutation was simulated every generation in each individual by randomly drawing a Poisson number of mutations, each with effects drawn into a given DFE, and summing their effects to produce the mutant offspring. When considering trait-based models, genotypes where characterized by their breeding value in *n* dimensions 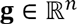. Mutation effects on traits were drawn into a given multivariate distribution and the fitness was computed as 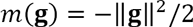 (quadratic landscape models, or ‘generalized FGM’, see Application section).

### Numerical solver

A numerical solver of Eq. **[7]**, applied to the FGM, is provided as a Matlab^©^ source code in **Supplementary material 1**, together with a Matlab^©^ graphical user interface and code for individual based simulation. The solver is based on a finite difference method with variable step sizes in *z* (smaller steps near *z* = 0, to get accurate values of the derivatives 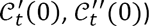 and an implicit scheme in time. Because of the transport term 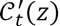, which tends to translate the solution towards the left with speed 1, the solution was computed on a finite interval 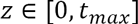 where *t*_*max*_ is the duration of the simulation. See Section V in Appendix D for more details. A Mathematica^®^ notebook is also available as **Supplementary material 2**: it provides a versatile (but less robust) solver (method of lines) of Eq. **[7]** and a code for individual based simulations, for four classes of models: non-epistatic models, Gaussian Fisher’s geometrical model (FGM), House of Cards and a simplified version of Rouzine et al.’s (2003) binary model.

## Application

Here we study various models for which the PDEs in Eqs. **[4]** and/or **[7]** apply. We distinguish three main applications: A) non-epistatic models of general form, B) epistatic models of general form, nearing an equilibrium, and C) epistatic models generated by quadratic fitness functions of phenotypes (FGM). All along we use the deterministic approximation, so we write ≈ to recall the approximate nature of our results.

### A. Non-epistatic models

Before tackling epistatic models, we first focus on context-independent mutation models, mostly to check that we retrieve previously known properties and to provide some new results. Because several results on non-epistatic models are already known, we put most of the results on this section in a dedicated Appendix C, and focused on new insights. As we have seen, any non-epistatic model is a trivial subcase of Eq. **[4]** with 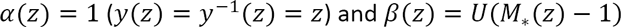: Eq. **[5]** yields

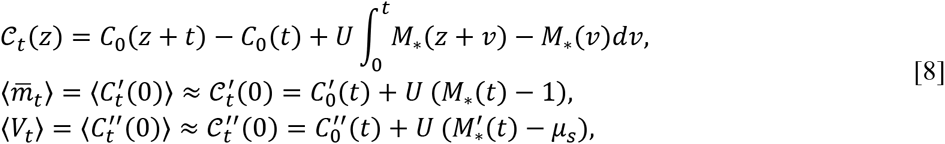

where we recall that *μ*_*s*_= *E*(*s*). This result essentially retrieves an alternative formulation of eq. (10) of (Desai and Fisher 2011), itself a continuous time version of Johnson’s (1999) eq. (13). These previous results both assumed purely deleterious mutations, which proves unnecessary in the derivation of Eq. **[8]**. Eq. **[8]** further allows for arbitrary standing variance in fitness via the additional term 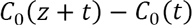, previously obtained for an infinite asexual population without mutation (Hansen 1992; Manna *et al.* 2012). As such, results in terms of CGFs or MGFs provide valuable information on the trajectory of moments, but are not so easy to fit on observed empirical distributions, which requires an explicit distribution function. In Appendix C II, we derive the stochastic representation of fitness from Eq. **[8]**, to help derive such functions. Supplementary **Movies 1A and 1B** illustrate the dynamics of the full fitness distribution for a negative gamma DFE and a constant DFE, respectively. In the parameter range chosen, the prediction from Eq. **[8]** accurately fits the observed distribution from the simulation of a single finite population of size *N=* 10^5^. Other illustrative examples are given in Appendix C.

#### Retrieving previous results

Several key known results on non-epistatic deleterious mutation models are readily obtained from Eq. [**8**] (detailed in Appendix C), such as properties of non-epistatic mutation-selection balance with arbitrary DFEs. In particular, Johnson’s (1999) result for discrete fitness classes straightforwardly extends to continuous DFEs: the equilibrium fitness distribution is a negative compound Poisson, with Poisson parameter 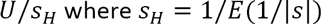 is the harmonic mean of the DFE in absolute value. Note that allowing for continuous distributions implies that the harmonic mean may be zero 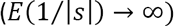, in which case the spike of fittest genotypes (with weight 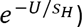) is *de facto* absent and the fitness distribution converges to a Gaussian (Eq. (C10) in Appendix C). Eq. **[8]** also implies that with arbitrary deleterious DFE and mutation rate *U*, the mutation load is 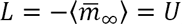, and the equilibrium variance in fitness is 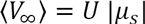. This extends a result previously derived as a low mutation rate limit (Burger and Hofbauer 1994) to the full mutation rate spectrum.

#### Timescales of load build-up *vs.* loss of accuracy with purely deleterious mutations

Eq. **[8]** allows to derive the ‘characteristic time’ *t*_*q*_ that it takes to reach some proportion *q* of the ultimate equilibrium 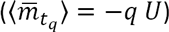. Neglecting standing variance, this time, *t*_*q*_ is the solution of 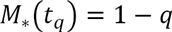: notably, it is independent of the mutation rate. This time can be computed for any given DFE, and admits simple bounds in the general case (see Appendix C III). For example, 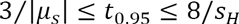, it takes between 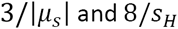 generations to reach 95% of the load. We recall that |*μ*_*s*_| and *S*_*H*_ are the arithmetic and harmonic means of the DFE in absolute value, respectively.

#### Non-epistatic models with beneficial mutations

When the kernel includes a portion of beneficial mutations (*M*_*_(∞) = ∞), mean fitness increases indefinitely 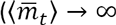 in Eq. [**8**]) and our approach overestimates this increase, after some time (see 3^rd^ section). For any non-epistatic model, the long-term fitness dynamics are best described by stochastic origin-fixation models (with or without clonal interference), once a stationary regime of fitness change has set. However, we propose that Eq. **[8]** can provide some connection between the transient and stationary regime and predict the fitness trajectory before stationarity (Section III in Appendix C). Assume a given rate *ν* of fitness change is predicted at stochastic stationary regime. If we assume a sharp transition from deterministic to stochastic stationary regime, this transition must then occur when the deterministic and stochastic models have equal rates of mean fitness change, namely at some time *t = τ* such that 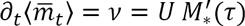 (Eq. **[8]** ignoring the contribution from standing variance). Up to this time, mean fitness is assumed to be given by the deterministic theory 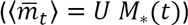 while it increases steadily at rate *ν* afterwards. This conjecture proves reasonable, as illustrated in **Fig. 2**. In **Fig. 2A** the DFE consists of purely beneficial, exponentially distributed, mutation effects 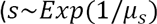, with *μ*_*s*_ > 0) and *ν* is given by clonal interference theory (eq. (16) in (Good *et al.* 2012)). In **Fig. 2B**, a shifted gamma DFE is considered: *s* ~ *s*_0_ + *x*, with *s*_0_ *>* 0 and x ~ −Γ(*a*, *b*) and the stationary rate *ν* is computed empirically, based on the adaptation rate that is observed at large times in the individual based simulations. Using only this rate *ν* as input, the transition time *τ* is computed and the full trajectory of expected mean fitness is predicted (see also **Figs. C2-C5** in Appendix C for other parameter values and another DFEs). By construction, theory (lines) and average from simulations (circles) should have the same slope *ν* in the late linear increase phase. However, they need not be superposed, especially over the full timescale studied. Coarse grain observation indeed suggests that the whole trajectory is surprisingly well captured by this simple heuristic. However, a transiently oscillating behavior (of the *average* trajectories) arises around the inferred transition time *τ* in all our simulations. This shows that the actual behavior is more complex than a simple transition from nonstationary/deterministic to stationary/stochastic (discussed in Desai and Fisher 2007).

**Fig. 2.**
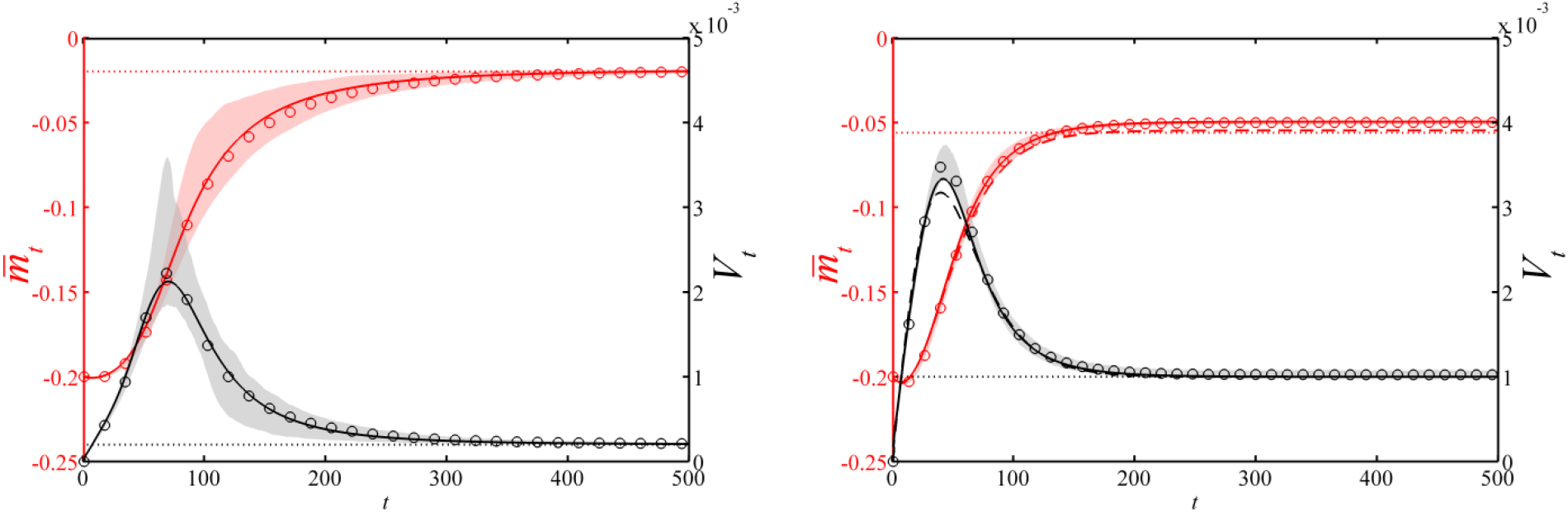
Mean fitness 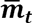 and variance *V*_*t*_ trajectories in non-epistatic models including beneficial mutations. (A) Exponential DFE: 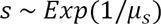 with mean effect *μ*_*s*_ = 0.001. (B) Shifted gamma DFE: 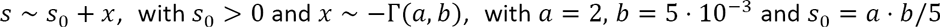. In both cases, *U=* 10^−3^. Plain lines: for *t < τ*, the expected trajectories 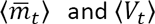 are given by our analytical theory (Eq. **[8]**); for *t ≥ τ*, the slope 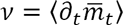 and the variance 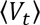 are kept constant. In panel A, the transition time 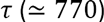 is such that *ν* equals the theoretical asymptotic slope given by eq. (16) in (Good *et al.* 2012); in panel B, the transition time 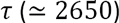 is such that *ν* equals the empirical slope observed in the individual based simulations during the interval *t* ∈ (4000,6000). Circles: empirical mean fitness and variance given by individual based simulations, averaged over 10^3^ populations (panel A) or 10^2^ populations (panel B), with *N = N*_*e*_= 10^6^; shaded regions: 99% confidence intervals for the mean fitness (in red) and the variance (in gray). We assumed initially clonal populations with *m*_0_ = 0.

In any case, the simulations in **Fig. 2** and **Figs. C2–C5** (Appendix C) show that the simple deterministic approximation does capture the dynamics over possibly several hundred (**Fig. 2A**) or thousand (**Fig. 2B**) generations (all the more as the proportion of beneficial mutations is small, apparently).

Furthermore, recall that this treatment only applies to thin-tailed DFEs (that fall off faster or as an exponential), otherwise the MGF is not analytic and *τ* → 0. The limiting case is an exponential tail, for which *τ* becomes smaller as the tail falls slower (larger mean). Yet, the simple heuristic did show good accuracy when simulating exponential DFEs with *μ*_*s*_= 0.01 *or* 0.001 (**Fig. 2** and **C2–C3**).

Finally, the **Figs. 2 and C2–C5** in Appendix C also illustrate that variation around the expected mean fitness explodes over time (red envelopes), especially after the transition to stationarity (late linear phase). Therefore, the empirical insight gained from the sole prediction of the expected mean fitness dynamics (without its envelope) can be *de facto* limited in this regime.

### B. Equilibrium in the presence of diminishing returns epistasis

Now consider an epistatic model 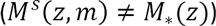, where beneficial mutations become less frequent and of smaller effect, as the population adapts, corresponding to a form of ‘diminishing returns’ epistasis. More precisely, we assume that (i) fitnesses are bounded on the right (the maximum fitness is then set at *m* = 0) and (ii) there is compensation (suboptimal backgrounds produce a portion of beneficial mutations). In this case, near equilibrium, the fitness distribution shrinks towards the maximum, and a 1^st^ order Taylor series of 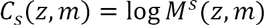 in small *m* yields 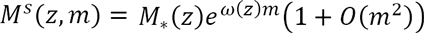. Here 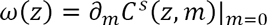 is the slope of the change with *m* of the CGF of the DFE, in the vicinity of *m=* 0, while 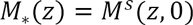 is the MGF of the DFE in the optimal background. Arbitrary models with diminishing returns epistasis (and a fitness upper bound), converge to log-linear background dependence near equilibrium. Then, by the memoryless property of log-linear background dependent models (see Eq. (B3) in Appendix B I), the CGF converges as *t* → ∞ to a unique equilibrium, independently of the initial CGF (the equilibrium cumulants are detailed in Appendix B I). Overall, mutation-selection balance is therefore a local attractor for this class of models and a global attractor for models with exact background dependence (such as the FGM).

In order to get further insight into the equilibrium fitness distribution, we now use a linear approximation to the MGF with small *m*, yielding 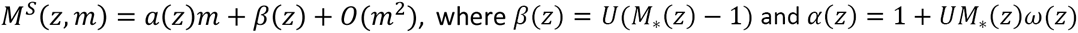. The asymptotic properties of Eq. **[5]** as ***t*** → ∞ (Section II.2 and II.3 in Appendix B) then yield a general theory for mutation-selection balance in the presence of diminishing return epistasis.

#### Mutation load

In particular, mean fitness stabilizes to 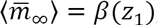, where *z*_1_ is the smallest positive root of *α*. Therefore, the mutation load is 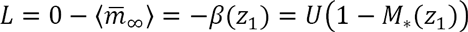. Two situations can occur: either *α* has no such root (*z*_1_ = ∞) in which case *L* = *U*, or it has a root 0 < *z*_1_ < ∞ in which case 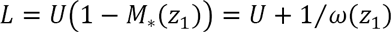. As 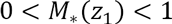, the load is then smaller than the mutation rate 0 < *L < U.* The first situation (*L* = *U*) always arises as *U →* 0. We thus have some form of ‘phase transition’ in the dependence of the load on mutation rate, as *U* increases.

#### Equilibrium fitness variance

We have 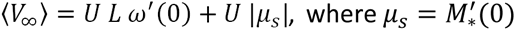, as above. Note that the term 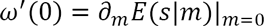 is the slope of the change in the mean of the DFE with *m* in the vicinity of *m* = 0. It seems likely that in most models, this slope is of same order as the mean itself: 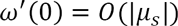. We thus have, *a priori*, 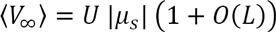 where we have seen that *L* ≤ *U;* therefore, the fitness variance is close to *U* |*μ*_*s*_| at equilibrium, in a vast variety of models (epistatic or not), as long as *U* ≪ 1. It is easily checked that the equilibrium for a non-epistatic model with deleterious mutations (see above) is retrieved as a subcase: 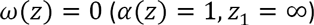 so that *L* = *U* and 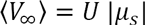.

#### Spike of optimal genotypes

a spike may exist (Section II.4 in Appendix B), but only provided the load *is L = U* and if 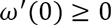, namely when maladaptation at most aggravates the mean deleterious effect of mutations (they become more or equally deleterious as the background gets suboptimal). The spike converges as *U →* 0, to that of the corresponding non-epistatic model with the same 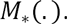 We have 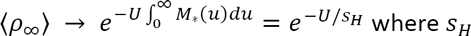, as previously, is the harmonic mean (in absolute value) of the DFE in the optimal background at *m* = 0. Furthermore, whenever *s_H_=* 0, the spike is vanishing at equilibrium, for any *U.* Finally, when 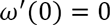 (as in the FGM), the weak mutation limit is also the upper bound 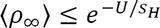 for any *U.*

Some of the qualitative results above are reminiscent of Burger’s (2000) propositions 2.1 p.127 and 5.1 p.145, proven for a single continuous trait, by a very different approach. It states that, independently of the trait mutational kernel or the trait-to-fitness function, the load (i) converges to *L = U* as *U* → 0, (ii) is exactly equal to *U* whenever a spike exists at the optimum, and (iii) is always less or equal to this limit 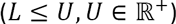. This section thus extends this result by providing a general approach to analytically compute these mutation loads, spike heights and higher moments, for *all U.*

### C. Fisher’s geometrical model

Let us now consider a classic model with diminishing returns epistasis: Fisher’s (1930) geometrical model (FGM), described in the Model section, as an example of log-linear background dependence.

#### Gaussian FGM

Recall that we denote Gaussian FGM the classic version with a multivariate normal distribution for mutation phenotypic effects, which shows exact log-linear context-dependence (**Table 2**) so that Eq. **[7]** applies.

#### Trajectories

The fitness mean and variance trajectories over time (predicted by numerically solving Eq. **[7]**) are illustrated for a small mutation rate in **Fig. 3A** (*U* < *U*_*c*_, see below for the definition of the critical value *U*_*c*_) and a high mutation rate (*U* > *U*_*c*_) in **Fig. 3B**. They are compared with the average fitness mean and variance in simulations (population size *N=* 10^5^). Smaller and larger population sizes and other mutation rates are illustrated in **Figs D2** and **D3** (Appendix D). The deterministic approximation is here accurate across the mutation rate spectrum (roughly as long *as* 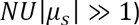. Note that, while the two first derivatives at *z* = 0 (expected mean 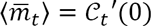 and variance 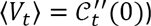 are accurately retrieved from the numerical solution of Eq. **[7]**, the third order derivative is more problematic to obtain (due to limited machine epsilon) and would require to solve the PDE satisfied by 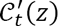, together with Eq. **[7]**.

**Fig. 3.**
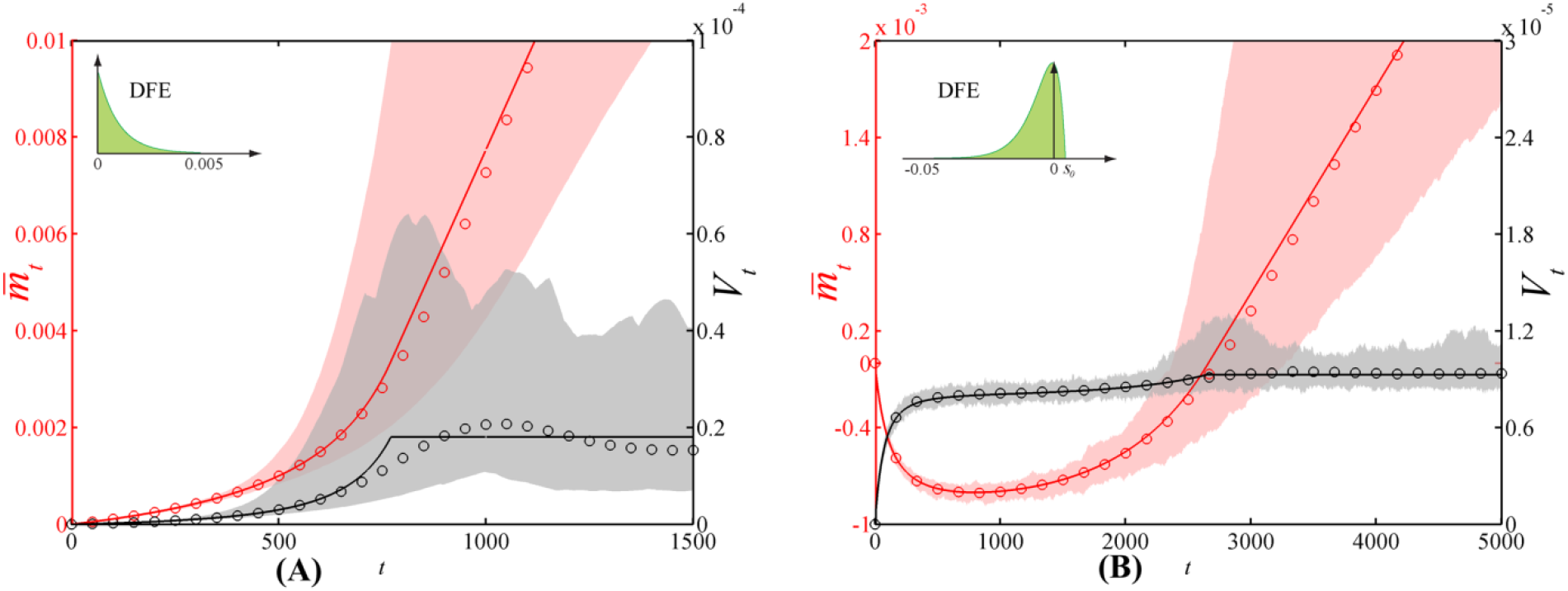
Mean fitness 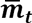 and variance *V*_*t*_ trajectories in a Gaussian Fisher’s geometrical model. (A) *U=* 0.02 < *U_c_;* (B) *U=* 0.1 > *U_C_.* Plain lines: expected trajectories 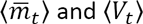 given by the numerical solution of Eq. **[7]**, with 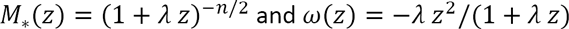. Dotted lines: equilibria 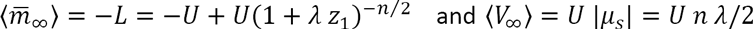 given by the analytical theory. Dashed lines (panel B): expected trajectories from the weak selection strong mutation (WSSM) approximation (Eq. **[12]** for 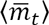 and (**B31**) for 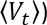. Circles: empirical mean fitness and variance given by individual based simulations, averaged over 10^3^ populations (*N* = *N*_*e*_= 10^5^); shaded regions: 99% confidence intervals for the mean fitness (in red) and the variance (in gray). The parameter values are *n=* 6 traits and *λ=* 1/300 (|*μ*_*s*_| = 0.01), leading to a critical mutation rate 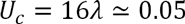. We assumed initially clonal populations with 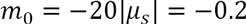.

#### Equilibrium

The equilibrium for the Gaussian FGM is a global attractor (by the memoryless property of log-linear background dependence models, Appendix B). Its properties are readily derived from the framework in Section **B**. (detailed in Appendix D), and summarized in **Table 3** (approximate results for *n* ≥ 3 are derived in Appendix E). Three qualitatively distinct situations arise according to the dimensionality *n* and mutation rate *U*, which determine the existence of a finite positive root to *α*. Consistent with the general results in **B.**, a ‘phase transition’ can occur (if *n* ≥ 3) at a critical mutation rate *U*_*c*_, which depends on dimension and scale (explicit formulae in Appendix D, section III and **Table 3**). The results are consistent with Waxman and Peck’s (1998) conclusions: a spike of optimal genotypes only exists at low enough mutation rate (*U* < *U*_*c*_) and in *n* ≥ 3 dimensions. Here an exact expression is obtained for the critical mutation rate where the spike vanishes, for the spike height below this threshold and for the mutation load over the full range of *U.* Note that explicit expressions for the spike height in *n=* 3 dimensions were also obtained (for a non-Gaussian FGM) in (Waxman and Peck 2006), by a different approach.

A simple approximation emerges for the equilibrium fitness distribution when *U* < *U*_*c*_ in terms of a mixture of a probability mass of optimal genotypes and a negative gamma distribution of suboptimal genotypes, corresponding to a Gaussian FGM in *n* − 2 dimensions (with *n* ≥ 3):

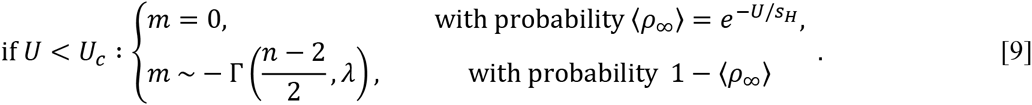

Strikingly, the weight of the spike is *exactly* the same as that in the corresponding non-epistatic model here (gamma DFE), whereas our heuristic analysis (Application **B.**) only suggests such convergence in the low mutation rates, in general. The full fitness distribution in Eq. **[9]** is exactly that expected in the absence of epistasis, in the small *U*/*s* approximation described in Appendix C II (Eq. C9). A simple pattern thus emerges: for any *U* < *U*_*c*_, the equilibrium fitness distribution in the FGM is approximately ‘blind’ to the presence of epistasis, and behaves as the equivalent non-epistatic model with DFE given by that of the optimal genotype. We thus retrieve essentially a House of Cards approximation (Turelli 1984) on fitness itself.

On the other hand, when *U* ≫ *U*_*c*_, a weak selection strong mutation (WSSM) limit (detailed below and in Appendix E) yields a complementary approximation for the fitness distribution at high mutation rates.

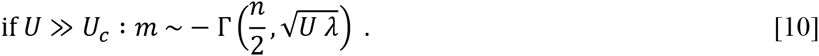

Finally, note that the equilibrium higher moments of Eq. **[7]** (exact for the Gaussian FGM) can be studied analytically (Appendix B) and are very close to the general expressions derived from the linearization in Application section **B.**.

**Table 3.**
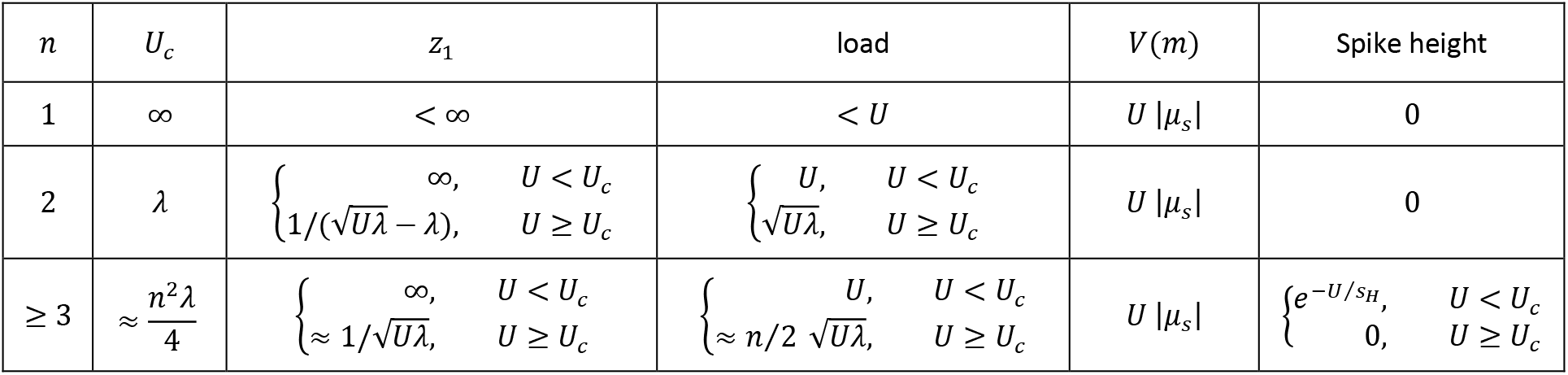
Mutation-selection balance properties in the Gaussian FGM. Here 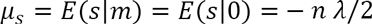 arithmetic mean of the DFE and 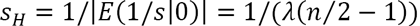, harmonic mean of the DFE. ′ ≈ ′ notifies that this is an approximate result (Appendix E).

#### Generalized FGM

Fisher’s original formulation did not specify the shape of the fitness function (linear, quadratic etc.) or the distribution of mutation effects on **g** (normal, uniform etc.), except that it must be spherically symmetric (effects are *iid* across traits), and centered on the parental phenotype. Keeping the quadratic fitness function, we study a ‘generalized FGM’ (see Appendix E) with arbitrary spherically symmetric distributions of mutation phenotypic effects 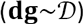. A given distribution 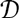 determines a given DFE in the optimal background (a given 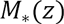). The function *ω* is then 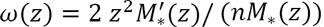, thus allowing the study of equilibria for any choice of 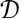 (Appendix E I). As an example (Appendix E II), we derive the equilibrium properties of a model with arbitrary dimension *n* and negative exponential DFE at the optimum: 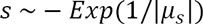. In particular, the load is 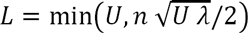, where 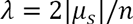 is the mutational variance on each trait (for consistency with the Gaussian FGM).

#### Weak selection strong mutation (WSSM) approximation

More general and simple results (Appendix E III) are obtained from a WSSM approximation, more precisely whenever 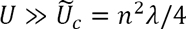 where 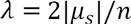 is the mutational variance on each trait. Note that 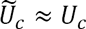 (for substantial *n*): it is roughly at the same mutation rate threshold that equilibria (*U*_*c*_) and transient dynamics 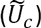 show a qualitative transition. In the WSSM regime, the mutational kernel is approximately linear in *m*, so that Eq. **[4]** captures the CGF dynamics, even away from equilibrium. The coefficients are 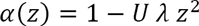 and 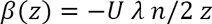.

#### Equilibrium

As was already stated above (Eq. **[10]**) the corresponding equilibrium fitness distribution is a negative gamma: 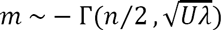. Connecting this approximation with the known value of the load at lower mutation rates *L = U* provides a simple expression covering all the range of *U:*

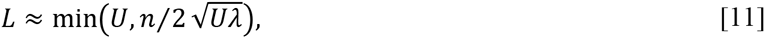

with a ‘phase transition’ at 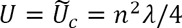. The accuracy of this simple result is illustrated in **Fig. 4A**, where the load is shown for single replicate simulations over a range of *U.* We simulated two alternative models (Gaussian FGM with a gamma DFE and an Inverse Gaussian DFE), scaled to the same value of |*μ_s_|* and with the same dimensionality *n.* Both models yield the same results, accurately captured by Eq. **[11]**. The spike of optimal genotypes is shown in **Fig. 4B** for the same simulations: here, all genotypes pertaining to an effectively neutral fitness class relative to the optimum 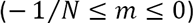 were counted as ‘under the spike’. As expected by theory, the spike weight is approximately 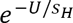, where *s*_*H*_ differs between the two models (Gaussian or Inverse Gaussian).

**Fig. 4.**
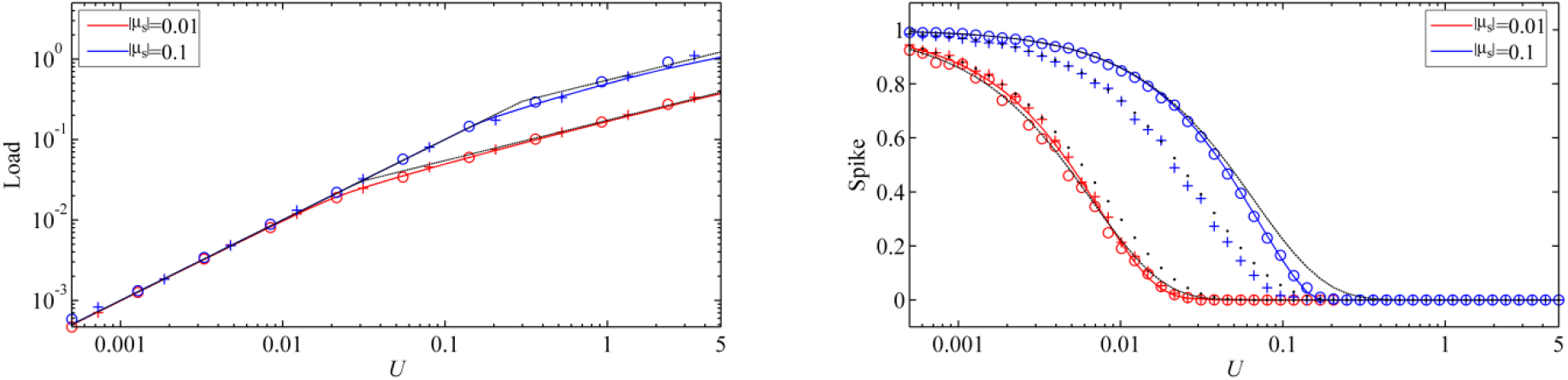
Mutation load *L* **(A) and spike** *ρ*_∞_ (B) as a function of mutation rate *U*: with two values of |*μ*_*s*_| (see legend) and with the standard Gaussian Fisher’s geometrical model (Gaussian FGM) or an FGM with Inverse Gaussian DFE at the optimum (IG FGM). Plain red and blue lines: numerical values obtained with the Eq. **[7]** for the Gaussian FGM (estimated at a large time *T=* 10^3^); the load (panel A) was computed as 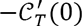 and the expected spike (panel B) as 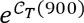. Panel A, black dashed lines: analytic approximations 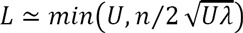 (Eq.[**11]**, panel A). Panel B, black dashed or dotted lines: 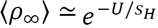, where *S*_*H*_ is the harmonic mean of the DFE (in absolute value) at the optimum (Gaussian or IG FGM respectively, Eq. **(D8)**). Circles (Gaussian FGM) and crosses (IG FGM): simulated values of the mutation load and of spike at time *T=* 10^3^ given by individual based simulations of a single population (*N* = *N*_*e*_= 10^5^). The parameter values are *n* = 6 traits and |*μ_s_|=* 0.01 or 0.1. The inverse Gaussian distribution has mean |*μ_s_|* and shape parameter 0.05.

#### Trajectories

The analytic solution (Eq. **[5]**) applied to the WSSM approximation can be equated, at all times, to a known explicit distribution, depending on the initial condition. The corresponding distribution of the underlying phenotype is also explicit, and happens, in all cases, to be multivariate Gaussian (with time-varying variances and means). Therefore, the WSSM approximation exactly matches Kimura’s (1965) and Lande’s (1980) Gaussian approximation for traits at equilibrium, and extends it away from equilibrium. Indeed, although obtained in very different manners, these two approaches rely on qualitatively the same WSSM assumption. Lande already conjectured that this approximation was mostly independent of the underlying distribution of mutation effects on phenotype, and should be valid away from equilibrium, as the dynamics of phenotypic variance are then independent of the mean (eq. (19) in Lande 1980). Here, the result arises explicitly as a WSSM limit of a generalized FGM. The present approach extends these former results to fitness (and trait) dynamics where the phenotypic variance is not constant, and provides an explicit threshold 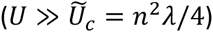, beyond which it is accurate. All results are given in Appendix E, we here only detail the mean fitness trajectories.

#### Adaptation from a clone

For a population started with a clone at given fitness *m*_0_ ≤ 0, the mean fitness trajectory, given by Eq. **[6]**, is

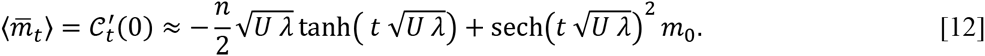

This WSSM approximation was illustrated in **Fig. 3B** (dashed lines) and proves fairly accurate even when *U* is only mildly superior *to* 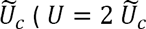 in this example). The corresponding trajectory of the full fitness and phenotype distributions are illustrated in supplementary movie files **(Movie 2A** and **2B** respectively), showing the agreement between simulations and theory, for a single replicate. The characteristic time of this fitness trajectory is the time *t*_0.99_ taken to fulfill 99% of the trajectory. Strikingly, it is independent of the details of the model: 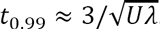. In particular, it is independent of the distance to the optimum (*m*_0_): it takes roughly the same time to reach equilibrium from an optimal or a highly suboptimal clone, in the WSSM regime.

#### Adaptation from an equilibrium population

In a similar manner, we may consider a population starting at equilibrium, undergoing a sudden shift in the optimum and responding to this new environment, this time with standing variance available. Here too, the whole fitness and phenotype distributions are explicit over time, including with a change in *U* or *λ* between the former and new environments. If the shift only affects the optimum (not *U or λ*) and is such that the mean phenotype shows a fitness lag *m*_0_ (mean fitness is then 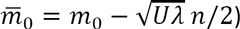, then

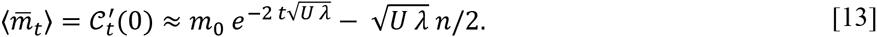

The trajectory of the fitness and phenotype distributions are illustrated in supplementary movie files **(Movie 3A** and **3B** respectively), with an additional doubling of the mutation rate in the new environment. Here too, the characteristic time is independent of the distance to the optimum 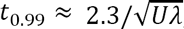, and it is only mildly shorter than the characteristic time in the absence of standing variance. In all cases the characteristic times scale with 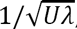, showing that the ‘cost of complexity’ well known in the FGM (Orr 2000) is only mediated by 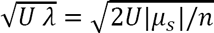 in this regime. When comparing different dimensionalities *n*, if we scale *λ* to the same 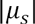, complexity slows down adaptation as 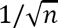. Otherwise, simply adding traits with the same variance *λ* does not affect the characteristic time, it simply increases the mutation load as 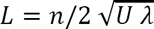.

## Convergence to the deterministic approximation

Our simulations, which included full stochasticity (individual based model) showed good agreement with the theory in Eq. **[3]**, that ignores drift. This seems to hold over either effectively infinite timescales (e.g. FGM, **Figs. 3–4**, and other models illustrated in **Supplementary material 2**), over very long timescales (non-epistatic models with purely deleterious mutations, **Fig. C1**), or over only a few hundred/thousand generations (non-epistatic models with beneficial mutations, **Figs. C2–C5**). Accuracy also seems to increase as *NU* gets larger for the models and parameters we explored. It has indeed long been observed that deleterious mutation models or models with an optimum could be handled reasonably well by deterministic population genetics. This then raises the question of why the deterministic approximation ultimately breaks down with non-epistatic models, whereas it does not seem to do so with diminishing returns epistasis.

This can obviously be tackled by individual based simulations for any given model. In the case of non-epistatic models, analytical studies have also pointed to a complex interplay of drift and other forces in the mid to long term behavior of asexual models (e.g. Desai and Fisher 2007): the importance of the “stochastic edge” of the fitness distribution (Brunet *et al.* 2008) depends on whether this edge is stochastic or not (highly populated or not). The present treatment provides some hint on the issue, by looking at the term neglected in the deterministic dynamics: *δ*_*t*_(*z*) in Eq. **[1]**. This ‘stochastic source term’ is negative, vanishes at *z* = 0 but increases with *z* (see Appendix B III). That *δ*_*t*_(*z) <* 0 means that the deterministic prediction overestimates the cumulants (for ex., the expected mean fitness is actually bellow the deterministic prediction). That δ_*t*_(*z*) is small about *z* = 0, means that the current error on the bulk of the distribution (the first derivatives of δ_*t*_(.) at *z* = 0) is limited. On the other hand, because of the transport term 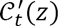 in Eq. **[3]**, the larger error |δ_*t*_(*z*)| for large *z* progressively affects the accuracy of the deterministic approximation around *z* = 0 (hence the bulk itself) at later times. Intuitively, this reflects the fact that sampling (drift) induces relatively more stochastic variation in the extrema than in the mean and variance of a distribution: the maximum can be very important for the long term rate of adaptation (“stochastic edge” Brunet *et al.* 2008), while the mean and variance influence the short term “bulk” dynamics.

Whether and when a substantial deviation will accumulate depends on the details of the model, and can be difficult to quantify. However, in the case of linear background-dependence (Eq. **[4]**) some general insight can be obtained, focusing on mean fitness trajectories. The relative deviation between the ‘exact’ expected mean fitness 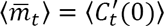 and that predicted by the deterministic approximation 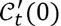, has an explicit upper bound at all times:

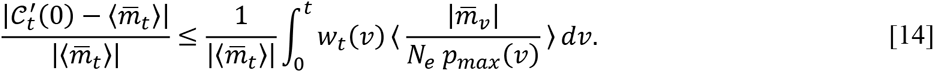

Here 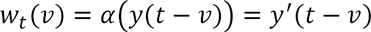 is a weight which depends on the form of epistasis (via *y*), see Eq. **[4]** and the paragraph below. Roughly, if 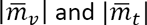 are of comparable order, the relative error is proportional to (i) *t/N*_*e*_ and (ii) to a weighted mean over the period (0, *t*) of the expected inverse frequency (across replicates) of the fittest class. Eq. **[14]** provides some intuition on how and why different mutation models deviate from the deterministic prediction. We treat each in turn.

### Non-epistatic models

The weights are then 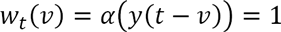, so the error must accumulate over time. With purely deleterious mutation models, *p*_*max*_(*t*) remains large for a long time 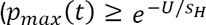 in the deterministic approximation), and it can be shown (Appendix C III.) that the relative error in Eq. **[14]** remains ≤ 1 for some ‘time to loss of accuracy’ of order 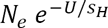 (see **Table 2**). The ‘characteristic time’ to reach 95% of equilibrium (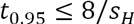, see section **A.**) is therefore often much less than the timescale over which the deterministic approximation breaks down and Muller’s ratchet starts to ‘click’ (of order 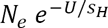, **Table 2**).

With beneficial mutations however, the fittest class consists of a small number of fit mutants so the error accumulates much faster. Furthermore, as the error depends on inverse frequencies of the fittest class, the fluctuations of this ‘stochastic edge’ (across replicates and times), especially through smaller values, are important, a fact already pointed out for these models (Hallatschek 2011).

### Diminishing returns epistatic models

With diminishing returns, two effects alleviate the deviation. First, mere intuition suggests that, as there is a reachable fitness upper bound, this fitness edge should ultimately become highly populated 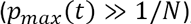, after sufficient time. This remains a verbal argument. Second, beyond the critical mutation rate threshold (whenever *α*(.) has a finite root), the weights *w*_*t*_(*ν*) in Eq.**[14]** vanish as *t* → ∞. This implies that the error ultimately becomes independent of the earlier dynamics of *p*_*max*_(*ν*) and remains bounded by a constant independent of *t* (see Appendix B, part III.2). This explains why these models are always accurately captured by the deterministic approximation at large times (see **Fig. 4** on equilibrium states), even when a substantial deviation from the deterministic trajectory builds up transiently (as observed e.g. in **Fig. D2** with *U=* 0.0002, *NU=* 2). Intuitively (without formal proof) we expect the transient error to be larger with smaller *NU* and when starting from a strongly maladapted population, as the fittest class may be small for a long time.

Qualitatively, this absence of accumulation of deviation over large times is a key difference introduced by epistasis. The result is reminiscent of Poon and Otto’s (2000), who showed that even a minimal amount of compensating mutations can stop Muller’s ratchet. A substantial transient deviation may arise at intermediate times, , but it ultimately shrinks again.

## Discussion

The proposed approach models the dynamics of fitness distributions in the presence of selection and mutation (neglecting drift), in large asexual populations, with a variety of distributions of mutation fitness effects (DFE). A deterministic PDE arises as an approximation to the dynamics of the expectation (over stochastic replicates) of the cumulant generating function (CGF) of the fitness distribution. This allows to easily handle clonal interference between co-segregating mutants (drawn from various classes of mutation models), and the contribution from standing variance, at or away from stationary regimes.

### Main results and possible empirical tests

When considering only the contribution from standing variance (negligible contribution from *de novo* mutation), Eq. **[8]** with *U* = 0 predicts the full fitness distribution over time from arbitrary initial condition. This provides a versatile model for the response to selection of large polymorphic asexual populations, over short timescales, i.e. before new mutations contribute to adaptation. The predicted trajectories are highly testable in experimental evolution: it only requires a measurement of the initial fitness distribution. We hope it may foster empirical tests of adaptive dynamics from standing variance in model asexual organisms, with a potential for faster observable responses than when a single clone adapts by new mutations.

The use of CGFs also simplifies the treatment of non-epistatic models with fairly general DFEs (**Figs. 2, C1–C5**). For non-epistatic deleterious mutation, most previous results are retrieved as a subcase (see Application **A.**). We further find that the fitness distribution admits explicit (testable) form over time (Appendix C, **Fig. C1, Movies 1A & 1B**), that the timescale to reach equilibrium from an optimal clone is independent of the mutation rate *U* (and of order 1/|*μ*_*s*_|), which is easily smaller than that over which the deterministic approximation breaks down.

When non-epistatic beneficial mutations are added, the approach breaks down over shorter timescales (detailed in Model section and Application section **A.**, **Table 2**). In general, the deterministic approximation breaks (after some time) when the fittest class is only represented by a few copies (see Appendix B III), forming a “stochastic edge” (Brunet *et al.* 2008). However in this case, we observe by simulation that Eq. **[8]** still provides a rough connection (**Fig. 2**) between the early regime of adaptation (deterministic), and the ultimate stationary regime (stochastic). Because Eq. **[8]** easily handles a wide variety of DFEs (e.g. including beneficial and deleterious mutations) which are not easily treated in the stationary stochastic regime, it may also be used as a more general null model over shorter empirical timescales (albeit still ignoring epistasis).

The same framework can be applied to mutation kernels showing diminishing returns epistasis (Application **B**. and **C**.). In that case, the discrepancy with the deterministic approximation remains bounded (sometimes very small) at all times (**Figs. 3 and 4 and Supplementary material 2**), because the fittest class is rapidly filled with a substantial number of selected mutants. The fitness distribution and the proportion of optimal genotypes at equilibrium then take testable explicit forms (Application **B**. and **Fig. 4**) in a variety of diminishing returns epistasis models where beneficial mutations compensate suboptimal genotypes. Overall, our most robust prediction (both with and without epistasis), at equilibrium, is that fitness variance should be close to 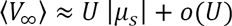, whenever *U* ≪ 1. This is also testable (given a large population at equilibrium), as the product *U* |*μ*_*s*_| can be directly estimated from mutation-accumulation experiments (reviewed e.g. in Keightley and Eyre-Walker 1999). It is also easier to estimate the fitness variance (and possibly skewness etc.) than the mutation load, as the latter requires an estimate of the maximal fitness. Such estimate would only be possible if optimal genotypes were frequent (not always the case), or given a particular model for the equilibrium fitness distribution (e.g. Eqs. **[9]**–**[10]**), which depends on the assumed DFE at the optimum.

The approach via CGFs is also particularly well suited for the Gaussian FGM with normally distributed mutant phenotypes. This model has recently served as a landmark null model of context-dependent DFE (background and/or environment dependence, Tenaillon 2014). It has also long been a landmark tool in evolutionary ecology and quantitative genetics: most treatments of the adaptive and demographic responses to environmental challenges, or of the distribution of phenotypes under stabilizing selection are based on its assumptions (Tenaillon 2014). Under this Gaussian FGM, the fitness dynamics (averaged over replicates) are fully captured by a single PDE (Eq. **[7]**, **Fig. 3**) covering the full mutation rate spectrum. Known analytical treatments of this model mostly described equilibria under two extreme regimes: in the limit *U* ≪ |*μ_s_|* with *n=* 1 dimension (Turelli 1984) or in the limit *U* ≫ |*μ_s_|* with arbitrary *n* (Lande 1980). Here, the full fitness distribution, at or before equilibrium, is predicted (analytically or numerically by solving Eq.[**7**]), for all *U* and arbitrary *n* (Appendix D, **Fig. 3 and 4**, **Movies 2 and 3**). This yields a fully testable pattern to fit to observed fitness distribution or mean fitness dynamics.

Finally, the results extend to arbitrary (spherically symmetric) distributions of mutant phenotypes, in a weak selection strong mutation (WSSM) limit 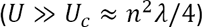. In this limit, both traits and fitness, at all times, converge to simple analytic distributions, independently of the details of mutational effects. This limit (**Fig. 3B** and Appendix E) arises here as a diffusion approximation in fitness space, and corresponds to normally distributed phenotypes (with time-varying mean and variance), consistent with M. Kimura’s derivation at equilibrium in one dimension (1965), and R. Lande’s conjecture for multiple dimensions (Lande 1980). The approach extends these theories away from equilibrium and clarifies the threshold mutation rate 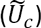 where they apply. These trajectories are also highly testable. Indeed, (i) the full distribution is analytic at all times from known initial condition (it may be applied on short experimental timescales) and (ii) the FGM can be parameterized (Martin and Lenormand 2006) from data on deleterious mutation effects (|*μ*_*s*_| and *n*) and rates (*U*), which are more readily available to the experimenter than beneficial mutation kernels and rates.

The evolutionary process inherent in the FGM is complex in large asexual populations and at high mutation rates: it includes clonal interference, both deleterious and beneficial mutations and pervasive epistasis. Yet, the resulting fitness trajectories in the WSSM limit (Eqs. **[12]**-**[13]**) display surprisingly simple and robust patterns, independently of the details of the underlying mutational process. In particular, the mean fitness (at any time away from equilibrium), scales simply with the initial maladaptation: 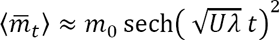, Eq. **[12]**. This latter pattern is, at least qualitatively, in agreement with the observation that the cumulative mean fitness increase (over stochastic replicates and between distant generations) scales almost linearly with initial maladaptation 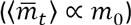. Couce and Tenaillon {, 2015 #3227} recently showed this empirical pattern to be hold across several species and datasets, and suggested that the FGM may be one among several models yielding such linearity. The present analytic treatment might allow to go beyond qualitative analyses and perform quantitative tests based on known parameters. A test of the FGM and other models would (ideally) require confronting full observed trajectories with (independently parameterized) predictions. We hope that the proposed approach may help such quantitative testing. Deriving (approximate or exact) analytic solutions to Eq. **[7]** away from the WSSM limit would also be useful in this regard, but requires further effort.

Finally, and although not detailed here, other epistatic models can be predicted analytically (Eq. **[4]**) or numerically (Eq. **[7]**) through the proposed framework. Two such examples are summarized in **Table 2**: Kingman’s (1978) House of Cards model (Eq. **[7]**) and a simplified version of Rouzine et al.’s (2003) binary model (Eq. **[4]**). Evaluating how accurate the predictions are, depending on the models and parameters, requires extensive simulation work beyond the scope of this article, but the necessary tools (and illustrations of the accuracy) are provided in **Supplementary material 2**.

### Limits

The model has several limits obviously; first of all, not all equations proposed here can be solved analytically (Eq. **[7]**) and we must then rely on numerical solutions. But more fundamental issues can be raised about the approach itself. We detail them below and discuss how to improve these aspects.

#### Genetic drift and clonal interference

Drift is explicitly modelled in Appendix A, but only to determine the error implied by neglecting it (Eq.**[14]**). Our results suggest that if the fittest class of genotypes quickly reaches (and remains at) substantial frequency, the deterministic approximation is accurate, even over the long term (see the section on ‘Convergence’.). This is typically what occurs with diminishing returns epistatic models (where fitness is bounded from above), which also prove to have a memoryless property that makes them less prone to accumulate stochastic deviations.

During adaptation over a single peak landscape, clonal interference is pervasive (multiple asexual lineages compete for fixation); yet, modelling the stochastic fate of each mutant does not prove critical in this model. Conversely, in similar conditions, it proves critical to do so with non-epistatic models of beneficial mutation, at least over long timescales. Overall, clonal interference need not always be described in the presence of drift: non-epistatic models with beneficial mutations, most studied in this context (Sniegowski and Gerrish 2010), happen to be a case where it is particularly important to do so. From an empirical perspective, it is simpler to avoid a theory that requires details of the genetic drift process, as the relevant parameters are notoriously difficult to measure (*N*_*e*_, the stochastic reproductive variance which may vary between genotypes etc.). However, a proper treatment of effect of stochastic forces (drift and mutations) would still be useful even in models where the expected trajectory is robust to their effect: it would provide envelopes around the deterministic expectation. Models of stochastic fronts and cutoffs may be used once translated into CGF dynamics, or stochastic PDEs using the results of Appendix A.

#### Segregation and recombination

Asexuals are our focus here, because they form the vast majority of model organisms in experimental evolution, for which this work is intended. However, sex is the norm *in natura* and will also likely become increasingly more studied empirically. The approach by CGFs was originally designed to handle recombinant genomes (Burger 1991), as the CGF from independent loci add up, providing simple extensions. Indeed, some of our results naturally extend to sexuals in simplified situations (not detailed here). However, fitness is typically non-additive across loci, so that simple additive theory may prove inaccurate in more realistic models.

#### Substitution data

The present model directly follows fitness dynamics, without explicitly modeling substitutions at the molecular level. They do occur (an allele becomes dominant, then another takes over etc.), but their dynamics may be complex (co-segregating alleles). By not requiring an explicit description of these dynamics, fitness trajectories in non-stationary regimes, with complex epistatic models can be handled. Yet, this is at the cost of providing no information on the underlying genetic basis of adaptation (which are now partly available empirically). For some important models, possibly epistatic but with low polymorphism, these underlying dynamics may be inferred from backward modeling. However, regimes with high polymorphism might show more complex molecular signatures, especially away from stationary regimes. The proposed framework may generate alternative coalescent models suited for epistatic, non-stationary models, just as travelling wave models have been successfully used (Good *et al.* 2014), for non-epistatic models at stationary regime.

#### More complex environments and landscapes

The models considered here mostly assumed a fixed environment in which adaptation occurs, as is typical in most theories of adaptation (Orr 2005), and as is relevant to many experimental evolution settings. However, more complex situations are of interest: multiple environments connected by migration, a continuously changing environment with a moving optimum, trade-off in life history traits. In some cases, these can be expressed as an adaptive process on multiple fitness components, and may then be handled by considering the dynamics of a multivariate CGF, describing the joint distribution of these components. Also, trait-based landscapes where traits are not equivalent for selection and/or mutation (e.g. anisotropic FGM) are not handled by the model as such. Indeed, the DFE is then not only dependent on the background fitness alone (distance to the optimum), but also on additional details (direction to the optimum). These can also be handled by introducing multivariate CGFs, describing the joint fitness contributions from each phenotypic dimension. We believe PDEs for such multivariate CGF dynamics can be written for many important classes of models where multiple fitness components interact. The open question will more likely be whether they can yield analytical insight.

#### Conclusion

We believe theoretical tools are now available that provide “null” adaptation models, which may be quantitatively confronted to experimental evolution data (including with standing variance, rarely studied in these experiments). Such tests of basic process predictions are necessary if we are ever to apply our theories quantitatively, into the wild or into the human body.

## Acknowledgments

This work was funded by INRA grant ‘MEDIA network’ and by the French National Research Agency (projects NONLOCAL, ANR-14-CE25-0013 and MECC, ANR-13-ADAP-0006) and the European Research Council under the European Union’s Seventh Framework Programme (FP/2007-2013) / ERC Grant Agreement n.321186 - ReaDi. This work was fostered by discussions with François Hamel, Marie-Eve Gil, Phillip Gerrish, Thomas Lenormand, David Waxman, Sylvain Gandon, Ophélie Ronce, Jimmy Garnier, and Luis Miguel Chevin.

## Appendix A: Derivation of the general PDEs for the evolution of the fitness distribution

### I. Notations and general setting

We call *t* the ‘time’ variable and *z* the real-valued argument of the moment generating functions (MGFs) and cumulant generating functions (CGFs) that we define below. We express the reference to time *t* by an index for compactness: *X*_*t*_(*z*) is the value of the function *X*(*t*, *z*). We denote 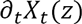 the 1^st^ derivative of function *X*(*t, z*) with respect to time *t* and 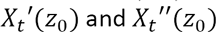 the 1^st^ and 2^nd^ derivatives, respectively, with respect to *z*, taken at *z* = *z*_0_. All integrals involving the probability density function (PDF) of a distribution are implicitly taken over the domain of this distribution. Expectations indicated as 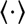 are taken over replicate populations (‘ensemble expectation’), while the overbar sign refers to the weighted average within a population at time *t.*

We consider a population of *N* asexual haploids, in continuous time (overlapping generations), measured in arbitrary units (hours, days etc.). This setting can also approximate a discrete time model (non-overlapping generations) when effects are small per generations, the time *t* is then measured in generations. We follow the dynamics of the distribution of the Malthusian fitness *m* (growth rate of a given genotypic class, hereafter ‘fitness’) under selection and mutation. At any time *t*, an arbitrary set of *K*_*t*_ genotypes, indexed by *i* ∈ [1, *K*_*t*_], with constant fitnesses *m*_*i*_, coexist in relative frequencies 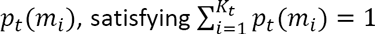. The approach can describe discrete classes (*K*_*t*_ finite) or infinite countable classes in the limit *K*_*t*_ → ∞. All co-segregating genotypes compete by frequency-independent selection, and mutate according to a Poisson process with fixed rate *U* per capita per unit time. The fitness of a mutant which parent has fitness *m* is *m + s*, where *s* is a random variable corresponding to the selection coefficient of the mutation relative to the parent. We measure fitness relative to a reference, set at *m=* 0 without loss of generality. Indeed, this reference is arbitrary because we are not considering demographic dynamics but only adaptation trajectories, namely relative fitness not absolute fitness. In those models that include some fitness optimum (e.g. single peak landscape models or models with only deleterious mutations), we set this optimum genotype to be the reference *m=* 0 for convenience (so that all *m* ≤ 0). In other models (e.g. models with background-independent beneficial mutations), the reference is just an arbitrary point in the fitness domain.

The distribution of *m* at time *t* can also be characterized by generating functions. We will consider the moment generating function (MGF) for a given finite population: 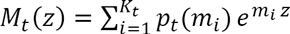, which for a finite number of genotypic classes, is always defined over the full line 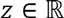. We will also consider the cumulant generating function (CGF): 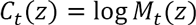. Note that, by definition of a probability distribution *M*_*t*_(0) = 1 and *C*_*t*_(0) = 0. Furthermore, the two functions are convex and *M*_*t*_(*z*) is positive on 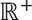.

For compactness, we use simplified notations for some key quantities: 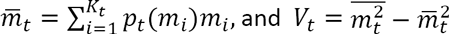 are, respectively, the mean and variance of the Malthusian fitness at time *t* for a given population (with 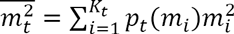). At any time, replicate populations may differ in the number (*K*_*t*_), fitness (*m*_*i*_) and frequencies 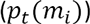 of co-segregating alleles. Averaging over these possible trajectories among replicates yields ‘ensemble expectations’. For the mean and variance in fitness within populations, the corresponding ‘ensemble expectations’ are the expected mean fitness 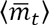 and the expected fitness variance within populations 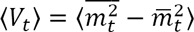. We also use simplified notations for the ensemble expectation of generating functions, under the deterministic approximation (see main text): 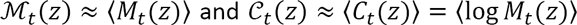 are the approximate expected MGF and CGF, respectively. We first describe exact dynamics for 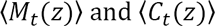, then introduce the dynamics of 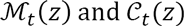 under the deterministic approximation.

### II. Effect of drift and selection

Accounting for genetic drift, the CGF *C*_*t*_(*z*) and MGF *M*_*t*_(*z*) themselves are random variables, generated by the random process governing the vector of frequencies of the *K*_*t*_ different genotypes 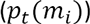 present at time *t.* For compactness, we note this vector 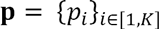 and *K = K_t_.* When population is large enough, this random *K* dimensional vector is approximately described by a *K*-type Wright-Fisher diffusion with selection and drift, characterized by a given variance effective size *N_e_.* The infinitesimal generator 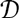 of this diffusion (eq. 4.83 p. 154 of Ewens 2004) can be expressed as follows, for any twice differentiable function 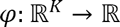 of the vector **p**

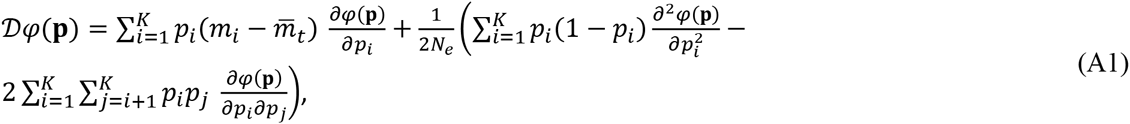

where 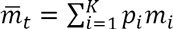 is the population mean fitness at time *t*. This infinitesimal generator formally describes the expected change of the arbitrary function *φ* of the random process **p**_*t*_ over some infinitesimal time interval 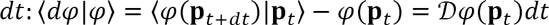. Recall that the expectation 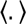 is taken over replicate populations. We wish to follow the dynamics of the unconditional expectation 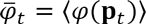 of the function 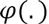, over time, over replicate populations with similar initial conditions 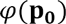.

Rattray and Shapiro (Rattray and Shapiro 2001) used a somewhat similar Wright-Fisher generator-based approach in the study of fitness cumulants, in a model of sexuals with constant effect mutation. The fitness distribution under study could thus be simplified to that of the number of mutations carried by each individual, assuming linkage equilibrium. Rattray and Shapiro’s model was not framed in terms of PDEs as here but rather as an infinite set of ODEs, solved numerically for some threshold level, (Burger 1991; Gerrish and Sniegowski 2012).

Good & Desai (2013) also obtained a similar result (see their Appendix D), in terms of the dynamics of the expected CGF, using Itô calculus. They worked on absolute numbers of lineages (while we consider diffusion on frequencies), assuming a constant effect of mutations (while we consider arbitrary DFE), but they obtained essentially the same results on CGF dynamics. Here we use an alternative method, via the Feynman-Kac theorem (theorem 8.1.1 in Øksendal 2003) and derive the dynamics of both the expected MGF and CGF.

**II.1 Dynamics of the expected MGF**. The MGF is a particular function of genotypic frequencies 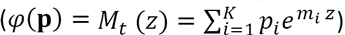, for which 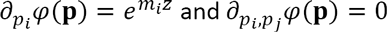. Eq. **(A1)** applied to this function can be written in terms of derivatives of 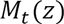 with respect to 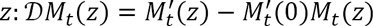 where we used the fact that 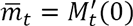. This only reflects the effect of selection on multiple gentoypes: drift induces no bias on the MGF as it is a linear function of allele frequencies, themselves unchanged, on average, by drift. Taking expectation over replicate populations starting from the same initial distribution **p**_0_, we can derive the dynamics of the expected MGF 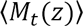 Using the Feynman-Kac formula, the expected MGF satisfies 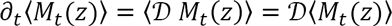, with initial condition 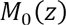, leading to the PDE:

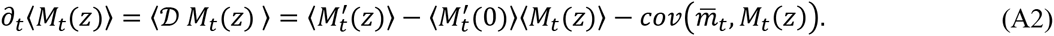

This equation is not closed as such, as it is affected by the covariance, across populations, of the mean fitness with the MGF: 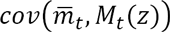, which itself will depend on higher order covariances.

**11.2 Dynamics of the expected CGF**. Consider now the CGF 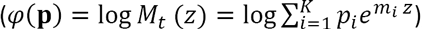, so that 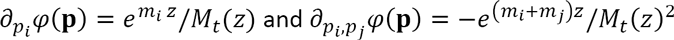. Eq. **(A1)** applied to this function can be written (recalling that 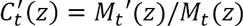 and 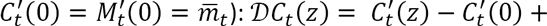 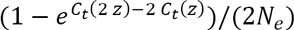. Starting from the distribution **p**_0_, and considering the expected CGF 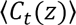, the Feynman-Kac formula leads to

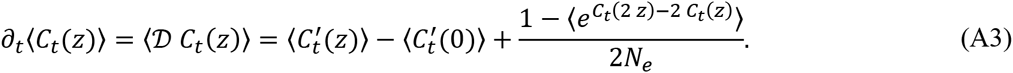

Note that the term introduced by drift, 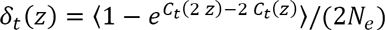, is the same as in Good & Desai’s (2013) eq. (D.4). Here again the equation cannot be solved unless we ignore this term, which is the deterministic approximation described in the next section.

**11.3 Neglecting the bias induced by drift**. For the rest of the article, we neglect the impact of drift on expected cumulants and moments, which boils down to neglecting 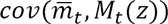 in (A2) and 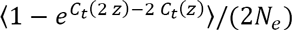 in (A3). We call this the “deterministic approximation”, and we define 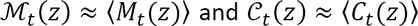 the expected MGF and CGF (respectively) under this approximation. Noting that ensemble expectation and derivation with respect to both *z* and *t* are exchangeable (linear operators), this yields a closed system for the dynamics of the approximate expected MGF and CGF:

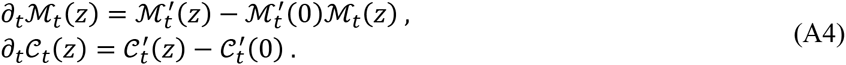

We then observe that 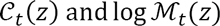 satisfy the same equation. The uniqueness of the solution of this equation (see Appendix B, II.1) thus implies that 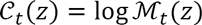; in other words, the deterministic approximation equates 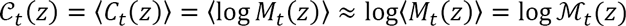.

This amounts to ignoring variation in 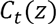 among replicates, relative to its expectation. Indeed, let the random deviation of 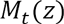 from its expectation be 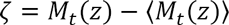 in any population. Then 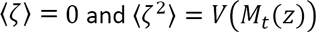, the variance in 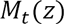 among populations. To leading order in 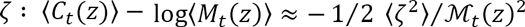, while 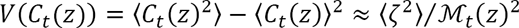. Overall, to leading order we have: 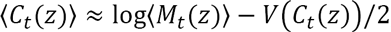, so the deterministic approximation (which equates 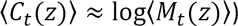 is consistent when 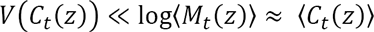. We discuss the timescale over which this approximation may be accurate, depending on the models considered, in Appendix B III (i.e. including in the presence of mutation).

### III. Background-dependent mutation

#### III.1 General expression

We allow the distribution of mutation fitness effects (DFE) to depend on the fitness *m* of the background on which they appear, and denote *f*(*s*|*m*) the probability distribution function (PDF, probability *density* function if the random variable *s* is continuous) of this conditional distribution. This *conditional* DFE remains constant over time, for each given background fitness class *m*, but the overall distribution of mutation effects spawned by the population may change over time, through the change in background distribution. We define the MGF of the conditional DFE as 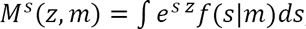, and the corresponding CGF 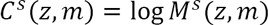. We assume that these quantities are well-defined and finite for any *m* attainable in the model, and over some interval 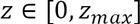: this means the DFE has finite moments in all backgrounds.

A single mutation occurs within a small enough time interval Δ*t* with probability *N U* Δ*t*(Poisson process). Given the effect *s* of the mutation and the background *m* where the mutation occurs, the conditional change in 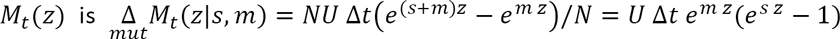.

Taking the expectation over the DFE *s* in background *m* yields:

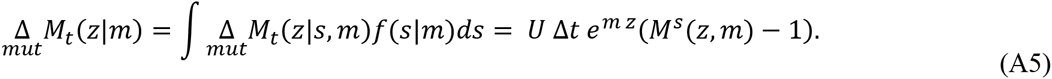

Then taking expectations over the background distribution *m* yields

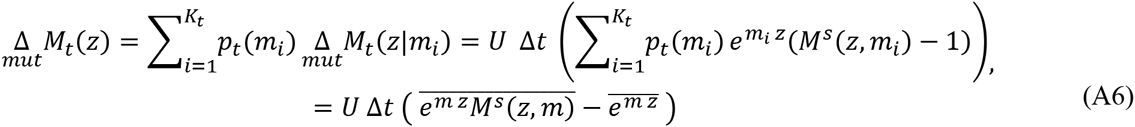

where the overbar refers to the weighted average within a population at time *t.* The corresponding change in 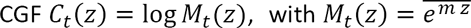 is obtained by noting that, with infinitesimal change in the continuous time limit (as 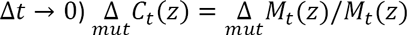

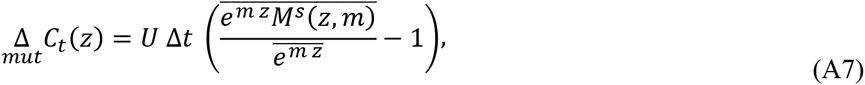

Taking ensemble expectation, over the stochastic trajectories of 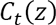 among replicate populations, yields an exact expression for the mutational contribution 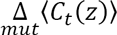 to the expected CGF 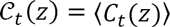:

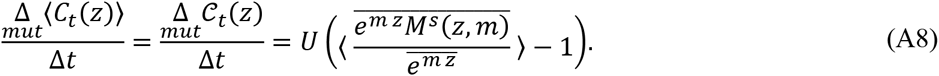

As such, the effect of mutation on 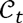 cannot be expressed in terms of 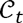 for general DFEs: further assumptions on the DFE are thus required to close the system.

#### III.2 Linear background-dependence

Situations may arise where the MGF of the DFE is linear in *m* : 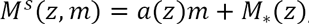, for some functions 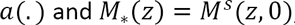, which is again the MGF in the background with fitness *m* = 0. This linearity may be exact or approximate, depending on the models and regimes. By definition of an MGF, 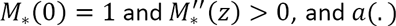 must satisfy *a*(0) = 0. Also we have 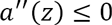 for all *z*, whenever the fitness set is bounded (so that *m* ≤ 0 with our convention). The MGF 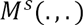 then satisfies the required properties 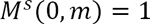 (conservation of total probability for all *m*) and 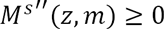 (convexity for all *m* with *m* ≤ 0). Note that when *m* is unbounded on the right (so that 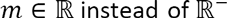), then convexity 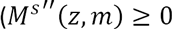 for all *m* and *z*) implies that 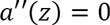 for all *z*. In this case, linear background-dependence can only be consistent with a function of the form 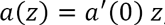, namely a DFE that is background independent, except for its mean: 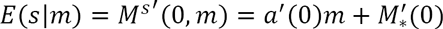.

Replacing 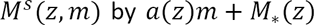 in the mutational term in Eq. **(A8)** yields 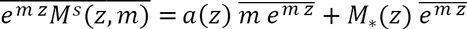. We then note that 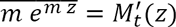, so that 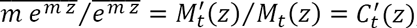, so that the mutational contribution in Eq. **(A8)** can be written: 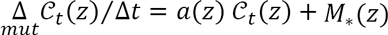. Taking a continuous time limit (Δ*t* → 0), the full dynamics of the expected CGF under the deterministic approximation is a nonlocal linear 1^st^ order PDE:

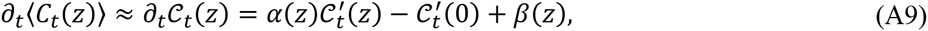

with 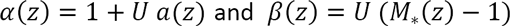. This PDE is studied and solved analytically in Appendix B.

##### Application: binary model (BM)

One example of mutation model that has linear background dependence is simplified version of Rouzine et al.’s model of asexual sequence evolution. In this model, genotypes are composed of Λ bins representing sites (*L* in the original paper, but we use other notation to avoid confusion with mutation load here). We thus denote this model ‘binary model’. Each bin is 0 for wild-type allele or 1 for mutant allele at the site and a given genotype *i* is a vector 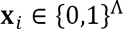. Each mutant allele incurs a deleterious effect – *δ <* 0 on Malthusian fitness (‘*s’* in the original paper: again notation is changed to avoid confusion with the random variable *s* describing selection coefficients in the present paper). Fitness is additive across sites so that *m*(*X*_*i*_) = *−δ k* for a genotype carrying 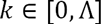 mutant alleles. Mutation is symmetric at each site with rate *u* per capita per generation per site: mutant alleles mutate back to wild type (effect + *δ*) and wild type alleles mutate forward to a deleterious mutant (effect − *δ*). The net genomic mutation rate is 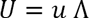 per generation per capita. Conditional on a mutation occurring, it hits a site at random and has an effect *+δ* if it hit a mutant allele (probability *k*/ Λ) or *−δ* if it hit a wild type allele (probability 1 − *k*/Λ). This DFE has stochastic representation 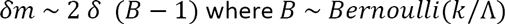. Recalling that *k = −m*/*δ*, we can write the MGF of this DFE as a function of the background fitness *m*:

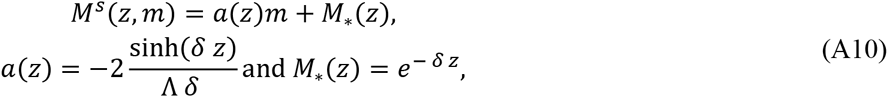

Which is a linear background-dependent model. This is a simplification of the original model in (Rouzine *et al.* 2003) in that they introduced an extra parameter *q* which is a proportion of sites that may compensate for a deleterious mutation at a given site. In the present ‘binary model’, *q = k*/Λ is by construction the current proportion of sites that are in ‘mutant’ state.

##### Application: Background - independent models

<u>Bürger’s (1991) model</u>: another instance where Eq. (A9) applies is for arbitrary background-independent models where 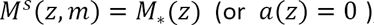, a given *M*_*\**_(*z*) characterizing the background-independent DFE. Logically, Eq. **(A9)** is then consistent with the CGF dynamics derived by R. Bürger (eq. 4.2 in (Burger 1991), for discrete time, when applied to a trait (fitness itself) with “exponential selection scheme”. Indeed here, Darwinian fitness (*W*, fitness for the discrete time model) is an exponential function of the trait *z = m* (here Malthusian fitness): 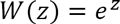.

<u>Diffusion in fitness space</u>: A simple form of background-independent model is one where mutation effects are modeled as a diffusion term in fitness space (Tsimring *et al.* 1996), yielding the so-called ‘replicator-mutator equation’ (Alfaro and Carles 2014). In this model, the mutational contribution on the dynamics of genotypic frequencies (on 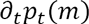) are described by a Laplace diffusion operator: 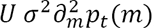, where *σ*^*2*^ is the mutational variance in fitness per generation. Multiplying this quantity by 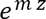 and integrating by parts with respect to *m*, it can be shown that the corresponding mutational input Eq. **(A8)** is 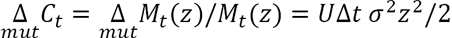, a quadratic function of *z*. In this model, drift is ignored so the deterministic approximation applies directly 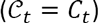.

Alternatively, this polynomial form of the mutational contribution arises as a weak selection limit of the background-independent kernel. Consider an arbitrary DFE with some MGF 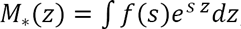, with mean 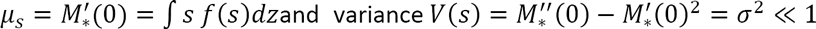. Consider the scaled variable 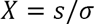, with MGF 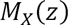: by definition, we have 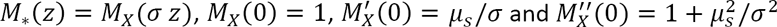. A Taylor series of the mutational kernel 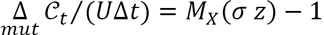, to leading order in *σ*, yields 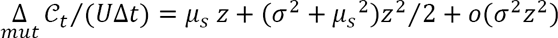, again a 2^nd^ order polynomial in *z*. Setting a symmetric DFE (*μ*_*s*_= 0) yields the exact same contribution as above 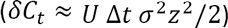: the diffusion kernel can be equated to a small variance limit of an arbitrary mutational kernel with zero mean, consistent with the rationale behind the diffusion approximation. This diffusion limit extends to include non-zero mean, but it can only be defined if the initial distribution has an analytical MGF, which amounts to the same condition as found in (Alfaro and Carles 2014): the initial distribution must fall off exponentially or faster, or put differently, it must have finite moments.

Note that such diffusion in fitness space can also be used with a context-dependent mutation kernel, from the background dependent mean and variance in fitness (see e.g. Appendix E for the FGM).

#### III.3 Log-linear background-dependence

Another possibility is that the dependence of the CGF of the DFE on the background fitness *m* is linear (so the MGF is log-linear):

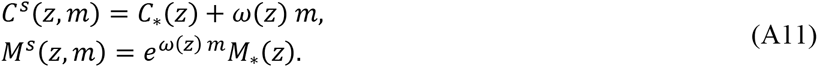

The function *ω*(*z*) describes how the DFE is affected by background fitness. By definition, 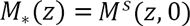 is the MGF of the DFE in the ‘reference’ background with fitness *m* = 0. Thus *ω ≡* 0 corresponds to background-independent mutations. In the case of background-dependent mutations 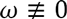 and we further assume that there is a fitness optimum at *m=* 0, so that *m* ≤ 0. The properties of *M*_*_ and *ω* are further detailed in Appendix B where this model is analyzed in detail.

Plugging Eq. **(A11)** into Eq. **(A8)** yields 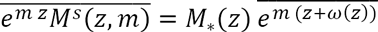 and thus the mutational contribution on the expected CGF has exact form 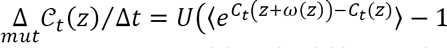. We have seen that the deterministic approximation amounts to equate 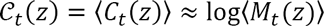 which implies 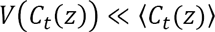. At roughly the same level of approximation, 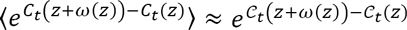, and the system can be closed Taking the continuous time limit (Δ*t* → 0), the full dynamics of the expected CGF under the deterministic approximation is then a 1^st^ order nonlinear nonlocal PDE:

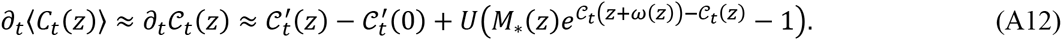

##### Application: Fisher’s (1930) geometrical model (FGM)

Our landmark example of background-dependent mutation is a particular version of Fisher’s geometric model (FGM), which we call ‘Gaussian FGM’. With *n* dimensions, the FGM generally assumes that fitness is a quadratic function of some *n-*dimensional vector of breeding values for phenotype 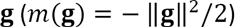, while mutations create a perturbation **dg** on this vector, that is unbiased and follows an isotropic multivariate distribution. In a particular version, the ‘Gaussian FGM’, this distribution is multivariate Gaussian: 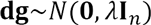, where 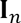 is the identity matrix in *n* dimensions. The reference is the optimal phenotype (**g = 0**) with fitness *m=* 0, and the conditional DFE, for the background with breeding value **g** and fitness *m = m*(g) has stochastic representation (Martin 2014): 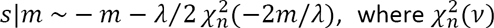 is a non-central chi-square deviate with *n* degrees of freedom and non-centrality parameter *v* (Martin 2014). The CGF of this DFE is exactly log-linear in background fitness:

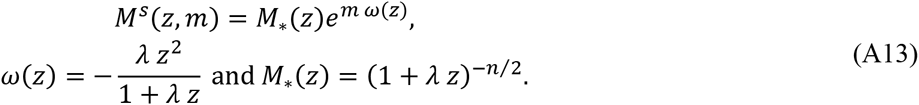

Eq. (A12) thus applies directly to this model. In appendix E, we show that a generalized version of this model with arbitrary, isotropic distribution of mutation effects on phenotypes, can be equated to a log-linear background dependent model when mutation effects are small relative to mutation rate.

##### Application: Kingman’s (1978) House of Cards model (HOC)

Under this model, mutants have a given fixed absolute fitness distribution *X*, which means that the relative fitness effects of mutations are background dependent (*s = X − m*, so that *m + s = X*). If 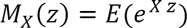 is the MGF of the fixed absolute fitness distribution, then the MGF of the DFE shows log-linear background dependence

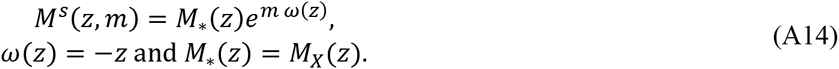

## Appendix B: formal properties and solutions of the general PDEs used in the article

This appendix is divided into two sections. In Section I, we study the mathematical properties of the nonlocal nonlinear equation **[7]** (main text), for 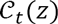 corresponding to log-linear background-dependence. In Section II, we solve the linear nonlocal equation **[4]** (main text) corresponding to linear background-dependence, and which can also be seen as an approximation of the nonlocal nonlinear equation **[7]** obtained by linearizing the MGF of the DFE, 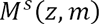 around *m=* 0. We use results from Section I to justify that the equilibrium of Eq. **[4]** is memoryless (independent of the initial fitness distribution) in the presence of epistasis. We also show that the two equations lead to consistent results at equilibrium (when such equilibrium exists). In the following, we consider only non-neutral mutation (the mutation rate *U* considered is a rate of mutation to non-neutral effects (*s ≠* 0)), so that the DFE has no Dirac mass at 0.

### I. Nonlinear nonlocal PDE for log-linear background-dependence

We investigate some a priori properties of the solutions of the nonlocal equation:

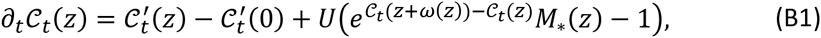

for *t* ≥ 0, *z* ≥ 0 and with the boundary condition 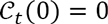.

Below, we first detail (Section I.1) the properties of *M*_*_ and *ω* that may be compatible with log-linear background dependence. Then (Section I.2) we show that the support of the fitness distribution instantaneously reaches the optimum *m=* 0 (or equivalently that 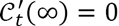, for all *t* > 0). Then we use this property to study the properties of the CGF as *t* → ∞: in particular the first cumulants (Section I.3) and the existence of a spike in the distribution (discrete probability mass) at the optimal genotype (Section I.4).

#### I.1 Properties of *ω* and *M*_*_

The function *M*_*_ is the MGF of the DFE for the genotype with fitness *m=* 0; it satisfies MGF properties *M*_*_(0) = 1 and 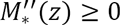. Furthermore, we consider only epistatic models that have an upper bound for fitness; whenever *ω ≠* 0, we thus set this maximum to max(*m*) = 0 without loss of generality. This implies that 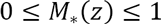 whenever *ω* ≠ 0; we do not require 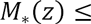 1 in the non-epistatic models (*ω=* 0).

For the well-posedness of Eq. **(B1)** we need that *z* + *ω*(*z*) ≥ 0 for all positive *z*. In order to establish this inequality, we recall that the optimal fitness was set at *m=* 0. Thus, the DFE is such that *f*(*s*|*m*) = 0 for all *s* ≥ *-m*, otherwise a mutant could overshoot *m=* 0. This implies that 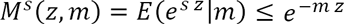. Using the log-linearity assumption 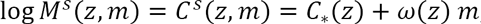, we get 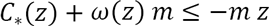, for all *m* ≤ 0 and *z* ≥ 0. Dividing this last inequality by *m <* 0 and passing to the limit *m* → −∞, we get: *z* + *ω*(*z*) ≥ 0 for all *z* ≥ 0.

Furthermore, the convexity of CGFs implies that 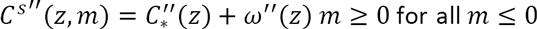. This implies that 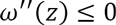 for all *z* ≥ 0, in other words *ω* is concave over 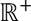. This in turn implies that 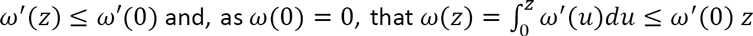. The inequality *z* + *ω*(*z*) ≥ 0 for all *z* ≥ 0 and the concavity of *ω* also implies that 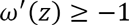 for all *z* ≥ 0. Then two cases may arise depending on 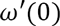. Note that 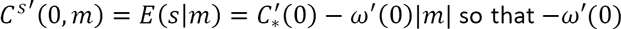 describes how the mean effect of mutations on fitness changes with increased maladaptation |*m*|.

<u>Case 1, ω′(0) ≤ 0</u>: in this case, the mean effect of mutations on fitness is unchanged or less deleterious as |*m*| increases: maladaptation alleviates mutation effects. Then *ω* is negative over 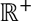 and 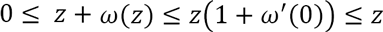. The nonlocal term 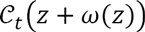 in Eq. **(B1)** applies to a value that always remains within the domain [0, *z*].

<u>Case 2, *ω*′(0) > 0</u>: in this case, the mean effect of mutations on fitness is more deleterious as |*m|* increases: maladaptation aggravates mutation effects. Then *ω*(*z*) may change sign over 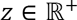 (and starts at positive values for small *z*).

Finally, we note that since the optimal fitness was set at 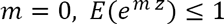 for all positive *z*, which implies that

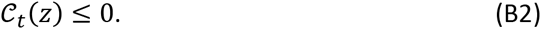

#### I.2 Flatness at infinity

As a key preliminary result, we show that, even if 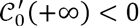, instantaneously any solution 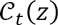 of **(B1)** becomes flat at infinity. More precisely:

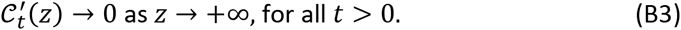

Intuitively, property **(B3)** arises because (i) we allow for beneficial mutations, (ii) the maximum fitness is set at *m* = 0, (iii) we ignore the stochastic loss of rare mutants and (iv) we consider continuous time. Therefore, even at very low mutation rates and after a very small period of time, a proportion of the fitness distribution (albeit initially infinitesimal) reaches the optimum. More rigorously, this result is achieved under any of the following biological/mathematical assumption:

*Assumption **H**: any background can mutate to the optimal background.* This means that max{*s*, such that 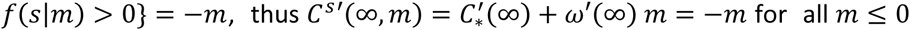 and for all *m* ≤ 0. This implies that 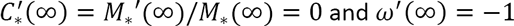.

or

*Assumption **H’**: any background can mutate to some fitter but suboptimal class.* Here, sup{*s*, such that 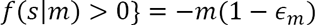, for some 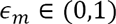 and for all *m* ≤ 0. In this case, 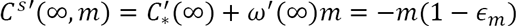 for all *m* ≤ 0, which implies that 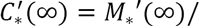 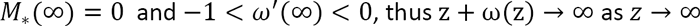.

**H** or **H’** imply that there is compensation, i.e., all suboptimal backgrounds (*m <* 0) produce at least some beneficial mutations. Thus, it also implies that, for all *m <* 0,

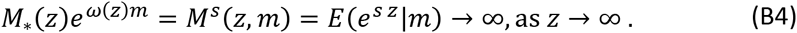

##### Proof of the property (B3)

Let us set 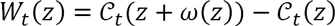. Using the convexity of the CGFs, we have 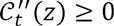; the function 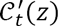 is therefore increasing (nondecreasing) in *z*. Thus, since 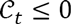 (Eq. **(B2)**), 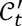 cannot reach positive values, otherwise it would remain positive at all larger *z* and 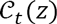 would converge to ∞ as *z* → ∞, contradicting **(B2)**. Thus, for each time 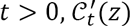 admits a limit 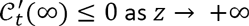. The function 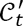 also satisfies the following equation:

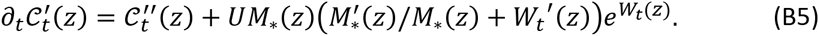

*Assume by contradiction that:*

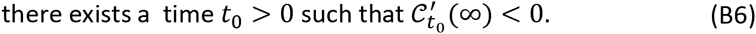

First, we compute 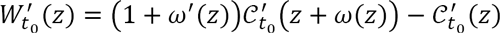. Using Assumption 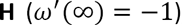, we get that 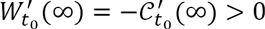. If we replace Assumption 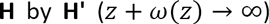, we obtain: 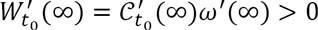. Thus, in all cases, for some constant *δ* > 0 and large *z*, we have:

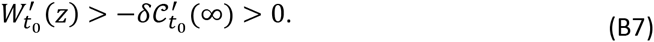

We know that 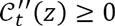 and, from Assumption 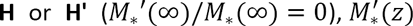 is negligible compared to *M*_*_(*z*) for large *z*. Equation **(B5)** together with **(B7)** and with *U* > 0 imply that 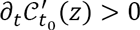 for large *z*.

Assume that there exists 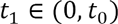 such that 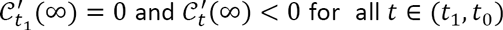. Again, 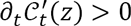 for all 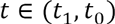 and large *z*. As a consequence, the limit 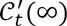 is a nondecreasing function of *t* ∈ (*t*_1_, *t*_0_) which implies that 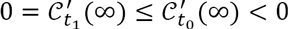 and leads to a contradiction. As a consequence, Property **(B6)** implies that 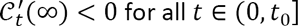.

The same arguments as above imply that for each 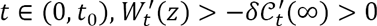 for *z* large enough and that 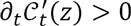 for all 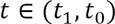. Therefore, 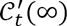 is nondecreasing for *t* ∈ (0, *t*_0_), and

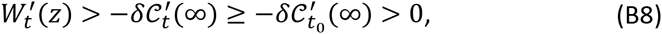

for all *t* ∈(0, *t*_0_) and large *z*.

Second, note that W_*t*_(*z*) can be bounded from below, for all *t* ∈ (0, *t*_0_). From Assumption **H** or **H'**, we know that *ω*(*z*) < 0 for large *z*. Then, we can write, for large *z*:

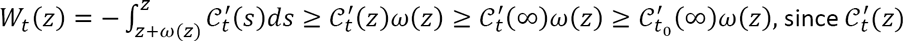 is nondecreasing in *z* and 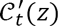 is nondecreasing in *t*. Using this lower bound, together with the formulas **(B5)** and **(B8)**, we get:

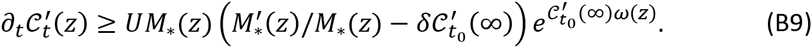

Let *t*_2_ ∈ (0, *t*_0_); integrating the inequality **(B9)** between *t*_2_ and *t*_0_, we get:

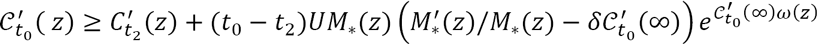, for all *z* large enough. Using Assumption 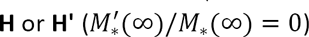 and Property **(B4)** which is a consequence of both **H** and **H'**, we conclude that 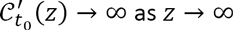. This contradicts the inequality 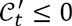. Finally, since Property

**(B6)** leads to a contradiction, we conclude that 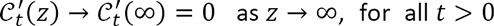. This concludes the proof of Property **(B3)**.

#### I.3 Stationary solution

Using the preliminary result of Section I.1, we are able to derive some properties of the long-time behavior of the solutions of Eq. **(B1)**.

##### First cumulant (load)

By definition, a stationary solution of the nonlocal equation **(B1)** does not depend of time; it is a solution 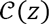 of:

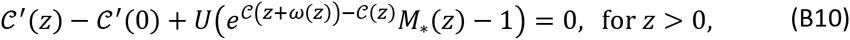

with the boundary condition 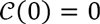. This equilibrium solution describes the fitness distribution at mutation-selection balance.

It is easily seen that the solution of **(B10)** is not unique. However, if the stationary solution is obtained as the limit of the solution 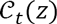 of Eq. **(B1)**, using the result **(B3)** of Section I.1, and passing to the limit *t* → ∞, we observe that 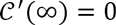. As a consequence, for any model where context-dependence affects the system (ω(*z*) ≠ 0) in a way that satisfies the Assumptions **H** or **H'**, we must have:

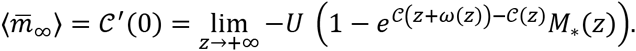

Then, two situations are possible, depending on *U, ω*(*z*) and *M*_*_(*z).*

*First case: 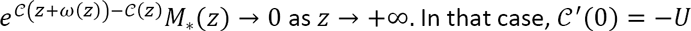.*

*Second case:* 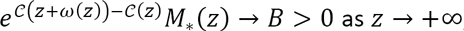, for some positive constant *B.* In that case, 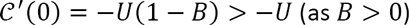.

This provides a general result on the mutation load, namely the difference between the maximal fitness possible (*m=* 0) and the mean fitness at mutation-selection balance: 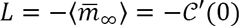. As the optimal DFE corresponds to a probability distribution function 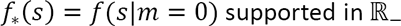 and with no "Dirac mass" at 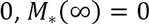: indeed, for any 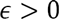 small enough and 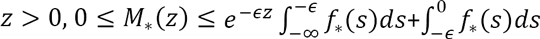, which shows that 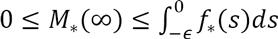 for all *ϵ* small enough and therefore

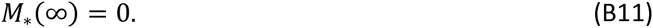

##### Higher cumulants

Differentiating the solution of Eq. **(B1)** with respect to *z*, and looking for stationary solutions, we can easily compute the cumulants 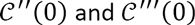 in terms of the functions *ω*(*z*) and *M*_*_(*z)* and of the load 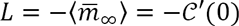 (recall that the maximal fitness is at *m=* 0):

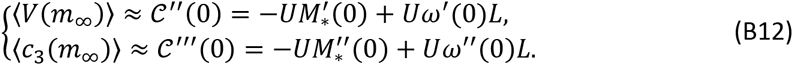

This provides an exact theory for the variance and skewness in fitness at equilibrium for models satisfying log-linear context dependence (such as Fisher’s geometrical model for example), but only given some explicit result regarding the load *L.*

#### I.4 Weight of the optimal genotype

Without loss of generality, we can write 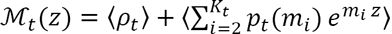, where *ρ*_*t*_ is the weight of the optimal genotype (with fitness *m*_1_ *=* 0) and *m*_*i*_ < 0 for all *i ≥ 2* and *t* ≥ 0. Passing to the limit *z* → ∞, we get:

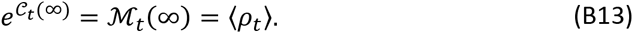

Thus, the limit of the MGF 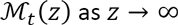 describes the expected weight of the optimal genotype *m=* 0 in the distribution PDF 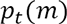. This weight is positive if and only if 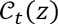 converges to some finite limit as *z* → ∞.

Consider now an equilibrium 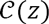 obtained as the large time limit of 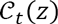. If the expected weight of the optimal genotype at equilibrium satisfies 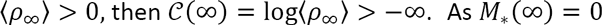 (Eq. **(B11)**), it follows that 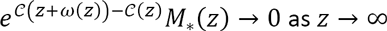. It then follows from the analysis in Section I.2 (first case) that 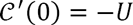. We thus have the following implication

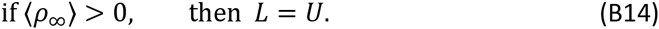

By contraposition, we obtain that if 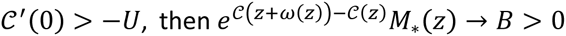 (second case in Section I.2), which, as 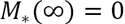, yields the following alternative implication

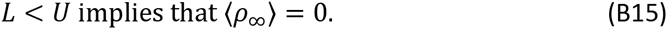

The above analysis (Section I.2) shows that necessarily the mutation load *L* at equilibrium satisfies *L ≤ U.* Furthermore, either *L = U* and there can be a spike at 0, or *L < U* and no spike can exist.

### II. Exactly soluble PDE for linear background-dependence

We consider the general linear transport equation with nonlocal term 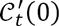:

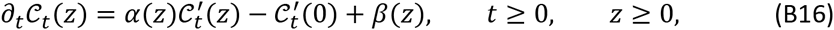

with the boundary condition 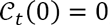, and where *α* is bounded from above and globally Lipschitz-continuous in [0, +∞), *β* is continuous and 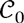 is continuously differentiable. We assume that *α*(0) = 1 and *β*(0) = 0.

To the best our knowledge, there is no general theory for solving this type of nonlocal PDEs. Here, we construct an explicit solution of this equation in terms of the solution of a simple ordinary differential equation (ODE).

#### II.1 General time-dependent solution: an explicit formula

First, we consider the solution *y*(*z*) of the ODE:

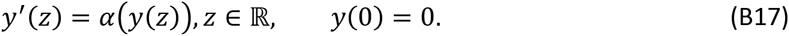

With our assumptions on *α*, the Cauchy-Lipschitz theorem implies that the solution *y* of the ODE **(B17)** exists and is unique. We then make the following change of variable:

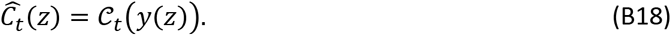

Since *α*(0) = 1, we observe that *Ĉ* satisfies the following equation:

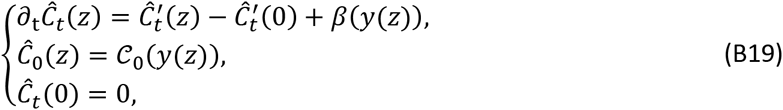

for *t* ≥ 0 and *z* ≥ 0. In order to get rid of the transport term and of the nonlocal term in this equation, we set, for some arbitrary constant *R* > 0:

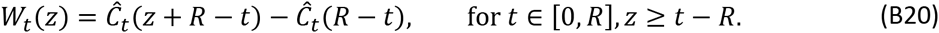

The definition of *W*_*t*_ implies that:

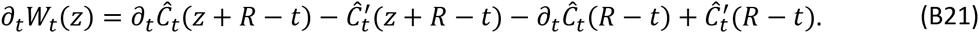

Coming back to (B19), we get that the function *W*_*t*_ satisfies:

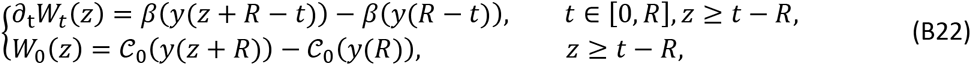

with the boundary condition *W*_*t*_(0) = 0. Integrating the above expression between 0 and *t*, we get:

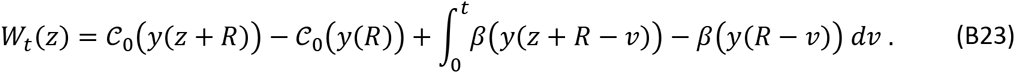

Using the definition **(B20)** of *W*_*t*_, we get *Ĉ*_*t*_(*z*) = *W*_*t*_(*z* − *R + t*) − *W*_*t*_(*t* − *R*), which leads to:

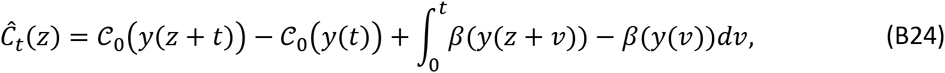

for all *t* ∈ [0, *R*] and *z* ≥ 0. Since *R* was chosen arbitrarily, the function *Ĉ*_*t*_ satisfies **(B24)** for all *t* ≥ 0.

Now, define *z*_1_ as the smallest positive root of *α*:

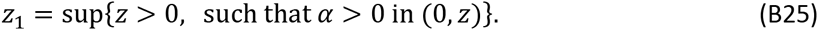

Either *α* > 0 in (0, +∞); in that case *z*_1_ = ∞, or otherwise *z*_1_ > 0 is finite. In all cases, the function *y*, defined by **(B17)** is a one to one and onto function from [0,+∞) to [0, *z*_1_). Then, using **(B18)**, we can write:

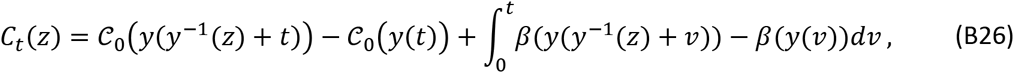

for *t* ≥ 0 and all *z* ∈ [0, *z*_1_). It is immediate to check that this is a solution of the problem **(B16)**, and by construction, it is the only solution of this equation. Note that 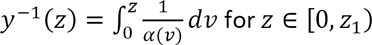.

Note that, when *α* and *β* are defined only in a finite interval [0, *z*_0_) Eq. **(B18)** can still be solved explicitly, but only for a finite range of values of *z* and *t:* the formula **(B31)** remains true whenever *y*(*y*^−1^(*z*) + *t*) < *z*_0_, or equivalently when 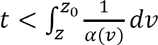.

**Cumulants.** Our objective here is to compute 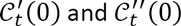. Differentiating the expression **(B18)** with respect to *z*, we obtain:

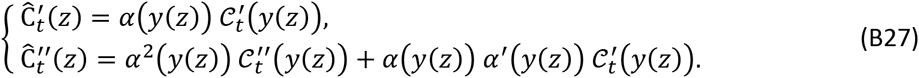

Computing these expressions at *z* = 0, we get:

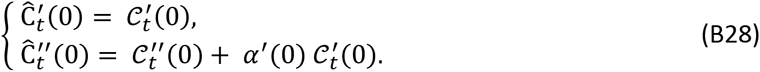

Differentiating the expression **(B24)** with respect to *z*, and computing it at *z* = 0, we get:

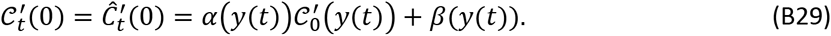

Differentiating two times the expression **(B24)** leads to the following expression for 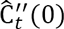:

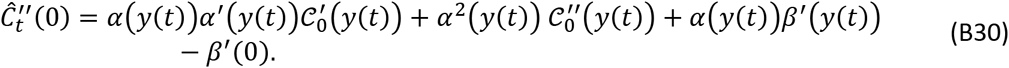

Using the expressions **(B28)** and **(B29)**, we finally obtain:

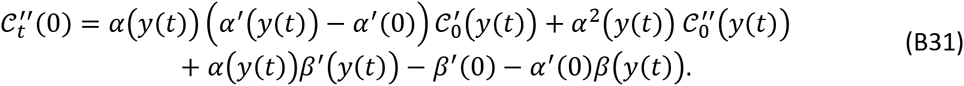

#### II.2 Stationary states and large time behavior

The stationary states of **(B16)** are the solutions of the following nonlocal ODE:

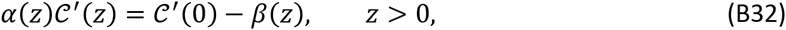

with the boundary condition *C*(0) = 0.

Let *z*_1_ be defined by **(B25)**. By definition, for all *z* ∈ (0, *z*_1_), *α*(*z*) > 0. For all *z* ∈ (0, *z*_1_) we can divide **(B32)** by *α*(*z*) and integrate between 0 and *z*:

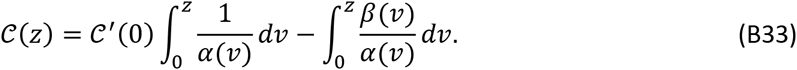

This gives an explicit expression for the stationary states of **(B16)**. Unfortunately, the stationary states are not unique since *C*′(0) is an arbitrary constant in this expression. Thus, *C*′(0) cannot be directly determined from **(B33)**. However, if the stationary state is obtained as the large time limit of the solution of **(B16)**, the expression **(B29)** implies that:

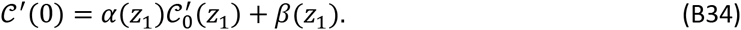

Then, three situations may occur.

*Case 1: z*_1_ *is finite*. In this case *α*(*z*_1_) = 0 and Eq. **(B34)** implies that *C*′(0) = *β*(*z*_1_).

*Case* 2: *z*_1_ = ∞ and *C*_0_(*z*) *coincides with the solution of the nonlinear model **(B1)** under the Assumption **H** or **H'***. If the linear equation **(B16)** was intended to be an approximation of the nonlinear model **(B1)**, the assumption 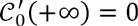 arises naturally from property **(B3)** of Section I.1: 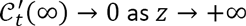, for any arbitrarily small time *t* > 0. Since *α* is bounded, we again obtain 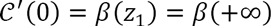.

Finally, in both cases, we obtain:

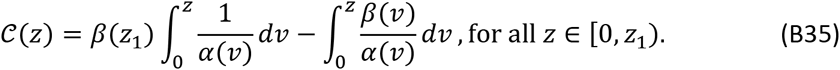

Thus, in spite of the dependence of 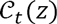 with respect to the initial condition 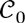 (see Eq. **(B26)**) we note that the reached stationary state 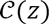 does not depend on 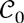, at least when context-dependence is present and of a form satisfying *Assumption **H** or **H'***.

Differentiating two times the expression **(B35)** with respect to *z* and computing the resulting expression at *z* = 0, we get:

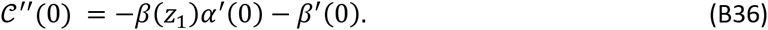

*Case 3: general case z*_1_ = ∞. In that situation, we cannot draw general *a priori* conclusions but we can solve the problem for an important particular case: any non-epistatic model. Indeed, these models are characterized by *ω*(*z*) = 0 in the fully nonlocal equation **(B1)**, which then reduces exactly to the linear PDE **(B16)**, with *α*(*z*) = 1. In such cases, *z*_1_ = ∞ but the Assumptions **H** or **H’** are not satisfied (since *ω*′(∞) = 0) so they do not pertain to *Case* 2 above. This case is fully treated in Appendix C.

#### II.3 Computation of the approached equilibrium cumulants

In order to compare the solution of the linearized model **(B16)** with the solution of the fully nonlocal and nonlinear equation **(B1)**, we apply the previous results to the particular case:

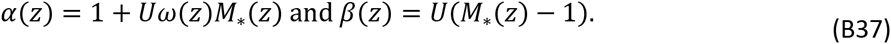

Eq. **(B16)** with these coefficients corresponds to the linearization of an arbitrary epistatic model when *m* → 0, they should thus be consistent with the results of **(B1)**, which is a particular form of epistatic model with log-linear context-dependence. Under this interpretation (see main text), *ω*(*z*) = ∂_*m*_ log *M*^*s*^(*z*, *m*) |_*m*=0_ is the slope of the change in the CGF of the DFE with *m*, as *m* approaches *m* = 0 (for backgrounds close to the optimal genotype), while *M*_*_(*z*) is still the MGF of the DFE in the optimal background.

*Case 1: z*_1_ *is finite*. In this case, the expression 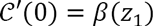 implies that the mutation load is:

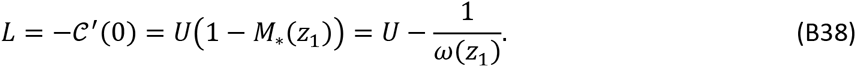

*Case 2:* 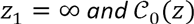 *coincides with the solution of the nonlinear model **(B1)** under the Assumption **H** or **H'***. As the DFE has no mass at 0, *M*_*_(∞) = 0 (see Eq. **(B11)**). The expression 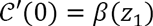 thus implies that the load is

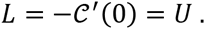

Overall, the results are consistent with the results obtained for the fully nonlocal and nonlinear equation **(B1)** (see Section I.3 above). Similarly, in both cases 1 and 2, formula **(B36)** leads to:

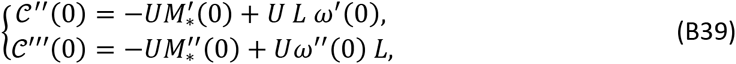

which is fully consistent with the result **(B12)** obtained while studying Eq. **(B1)**.

*Case 3: z*_1_ = ∞*, and ω* ≡ 0 *(context-independent models)*. As already observed, the fully nonlocal equation **(B1)** reduces exactly to the linear PDE **(B16)**. In this case, with *M*_*_(∞) = 0, and, as shown in Appendix C:

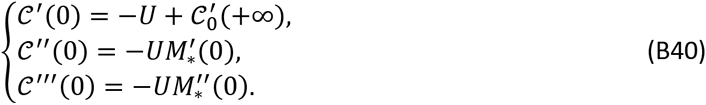

Dependence to the initial condition 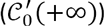 arises because the model contains no beneficial mutation here (otherwise *M*_*_(∞) = +∞), so the upper bound of the ultimate fitness distribution is the maximum of the initial one.

#### II.4 Spike of optimal genotypes at equilibrium

In model **(B1)**, a spike can only exist if 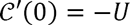 at equilibrium (Section I.4). Here, we focus on this situation and we derive an explicit formula for the weight of the spike by assuming that 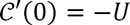 at equilibrium in the linearized equation **(B16)** corresponding to an epistatic model: 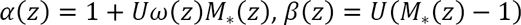.

From the analysis in Section II.3, 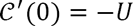 implies that *α*(*z*) has no root over 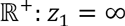. At this point, we recall that *ω* and *M*_*_ here have the same interpretation as for Eq. **(B1)**, so their properties should still apply. Therefore, we should have *ω*(*z*) ≤ *ω*′(0) *z* over 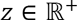, i.e., *α*(*z*) ≤ 1 + *U z M*_*_(*z*) *ω*′(0). Since *α*(*z*) has no root over 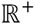, one must assume that ω′(0) ≥ 0.

Using the formula **(B35)**, we obtain that the equilibrium fitness distribution has 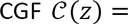 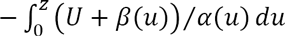. A spike may then exist and its expected weight is

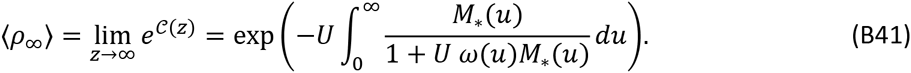

As *α*(*u*) = 1 + *U ω*(*u*)*M*_*_(*u*) has no root over 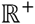 and *α*(0) = 1, up to a slight change in *U* in the pathological case *ω*(∞)*M*_*_(∞) = *-*1/*U*, we know that *α*(.) has a strictly positive lower bound over *z* ∈ 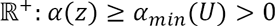, where 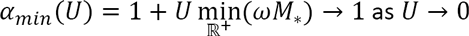. Similarly, we define an upper bound for 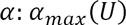, which may be finite or not, depending on *ω* and *M*_*_. Note that, when *ω*′(0) = 0, as in Fisher’s geometric model, 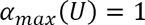. Finally, a lower and upper bound to the spike's expected weight at equilibrium are then given by:

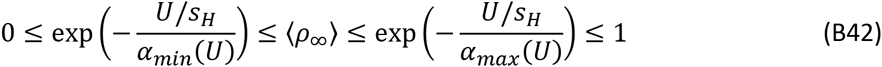

where 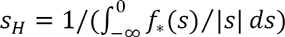 is the harmonic mean (in absolute value) of the DFE at optimum. This follows from the definition of 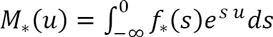, which implies that 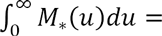 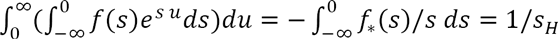.

If the integral 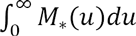 diverges (*S*_*H*_ = 0), then the spike vanishes (whatever the form of 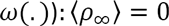. This is if 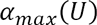 is finite, which depends on the form of *ω*(.) (as 0 < *M*_*_(*u*) ≤ 1), but should be the case with many models.

If the integral 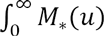 converges (*S*_*H*_ > 0), then the spike is non-vanishing and its expected weight can be approached as 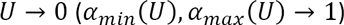 by

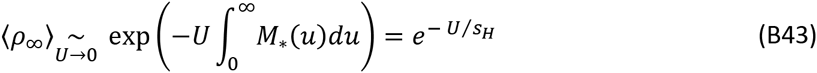

which corresponds to the predicted spike weight in a non-epistatic model (*ω*(*z*) = 0) with the same DFE at the optimum (characterized by the MGF *M*_*_(.)). If *ω*′(0) = 0, the mean of the DFE is unaffected by maladaptation, as in Fisher’s geometric model; then 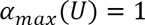 and the limit is also the upper bound: 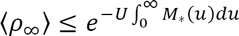.

### III. Long-term accuracy of the deterministic approximation

The PDEs which describe the dynamics of the expected CGF 〈*C*_*t*_(*z*)〉 and of the deterministic approximation 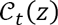 differ by a term:

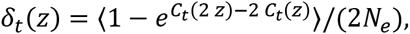

see Appendix A, part II (Eqs. **(A3–4)**). With linear background dependence, this is indeed the exact deviation between the dynamics derived from the diffusion generator (that account for drift) and from the deterministic approximation (Eq. **(A9)**). With log-linear background dependence, however, the mutational term is also approximated to obtain the closed system in Eq. **(A12)**. Yet, this second approximation is at the same order (it also assumes *V*(*C*_*t*_(*z*)) ≪ 〈*C*_*t*_(*z*)〉). Therefore, with this model too, the above error term should correctly describe the deviations between the exact non soluble system and the approximate PDE dynamics.

We derive here some properties of this error term *δ*_*t*_(*z*) and we analyze its effect on the difference between 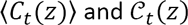. We mostly focus on the error in the expected mean fitness, 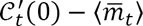, with linear background-dependence (epistatic or non-epistatic models).

#### III.1 Properties of the error term *δ*_*t*_(*z*)

By convexity of the CGFs and since *C*_*t*_(0) = 0, we have *C*_*t*_(2 *z*) - 2 *C*_*t*_(*z*) ≥ 0, which implies that *δ*_*t*_(*z*) ≤ 0. Thus, neglecting *δ*_*t*_(*z*) leads to an overestimation of 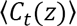, and consequently of the mean fitness 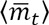. We can draw two broad qualitative conclusions on the short term error.

First, we observe that 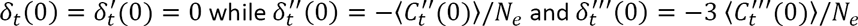. Therefore, when starting from the correct 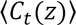 at some given time *t*, the error made by using the deterministic approximate dynamics to predict later times is small on the bulk of the distribution (the first cumulants, mean, variance, third moment). This does not preclude this error from accumulating over time, thus creating large deviations later, even on the bulk. More precisely, these errors must be compared to the other source terms due to mutation and selection, which are implemented in the deterministic approximation. *δ*_*t*_ does not directly contribute any error on the mean fitness dynamics, but for the variance and third moment, these source terms are of 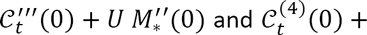 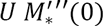 respectively (plus some potential extra terms due to linear background – dependence). For example, over some short term at least, the error made in the variance dynamics (relative to the deterministic prediction), remains limited if 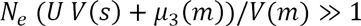.

Second, we also note that *δ*_*t*_(*z*) decreases with *z*: consider *h* ≥ 0, and compute *δ*_*t*_(*z + h*) − *δ*_*t*_(*z*) = 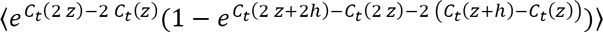. By convexity of the CGFs, *C*_*t*_(2 *z* + 2*h*) − 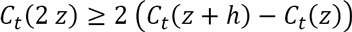. This implies that *δ*_*t*_(*z + h*) − *δ*_*t*_(*z*) ≤ 0 and the conclusion follows. Thus |*δ*_*t*_(*z)* | is bounded by |*δ*_*t*_(∞)|. If we define *p*_*max*_(*t*) the current frequency of the fittest class in a given replicate, with fitness *m*_*max*_(*t*) = max(*m*(*t*))) we notice that:

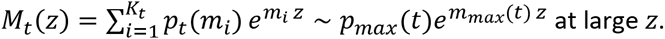

Then, coming back to the definition of *C*_*t*_:

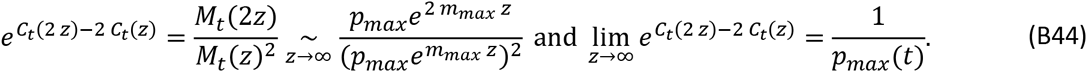

Finally, we thus get that

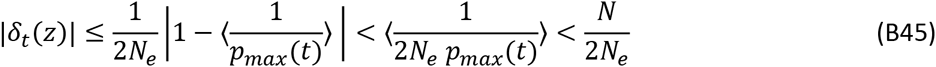

where the upper bound is obtained by noting that, at least, *p*_*max*_ ≥ 1/*N*. This upper bound is necessarily small whenever *N*_*e*_ *p*_*max*_ (*t*) ≫ 1, namely when the number of individuals in the fittest class is sufficiently large, in all replicates.

Therefore, whenever a large absolute number of individuals lies at the maximum of the current distribution, the error made at current time is also small. It will be the case with models that have an upper fitness bound (epistatic with an optimum, non-epistatic with purely deleterious mutations), once this bound is highly populated (of course the larger *N*_*e*_ the milder the criterion is in terms of *frequencies*). On the contrary, in non-epistatic models the deviation from the deterministic approximation can become substantial when the fittest edge of the distribution is small and stochastic.

#### III.2 Cumulative error with linear background-dependence

To get a more quantitative characterization of how discrepancies accumulate over time, let us define the deviation between the exact and approximate CGF at time 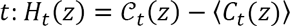. We reduce our analysis here to linear background-dependence (which includes all non epistatic and some epistatic models). We have shown in appendix A that the deterministic and ‘exact’ stochastic dynamics (under the diffusion approximation, actually) are given by:

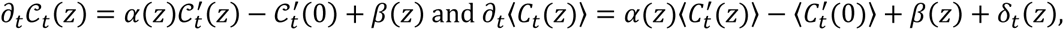

with *α*(*z*) = 1 + *U a*(*z)* and *β*(*z*) = *U* (*M*_*_(*z*) − 1), and we assume here that *α* ≤ 1. The deviation 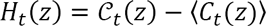 satisfies the PDE (with initial condition *H*_0_(*z*) = 0):

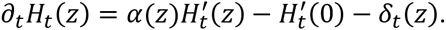

Considering *δ*_*t*_(*z*) as an external forcing term, this equation can be solved exactly (Part II.1 above), yielding 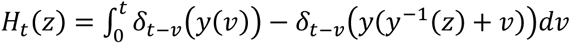, where *y* is the solution of the ODE *y*′(*z*) = *α*(*y*(*z*)), with *y*(0) = 0. As expected, this shows that *H*_*t*_(*z*) ≥ 0 since *δ*_*t*_ is a decreasing function of *z* and *y*(*v*) is increasing in *v*. The deterministic approximation overestimates the mean fitness trajectory over time. Computing *H*_*t*_(*y*(*z*)), we get:

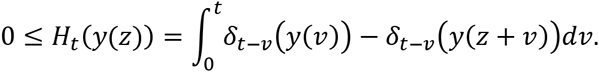

Dividing this expression by *z* and passing to the limit *z* → 0, (recall that *y*′(0) = *α*(*y*(0)) = *α*(0) = 1) we obtain

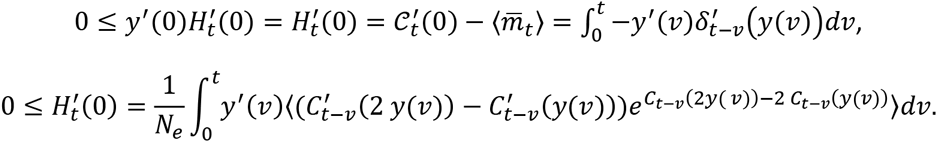

With a relevant choice of reference for relative fitness, the fitness of the fittest class is non-positive up to any given time (max(*m*)≤0). Thus, 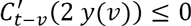 and as already observed in part III.1, 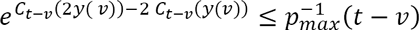. Moreover, 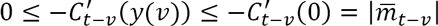 by convexity of *C*_*t*_(*z*) with respect to *z*. Overall, this shows that:

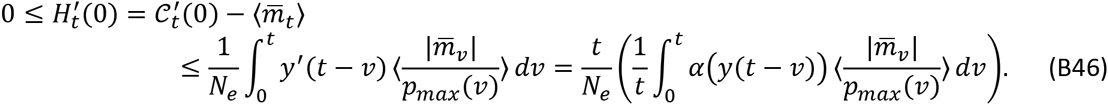

If 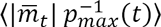 remains bounded by some positive constant *ε* after a transient period of duration *t*\*, we get for *t*>*t*\*:

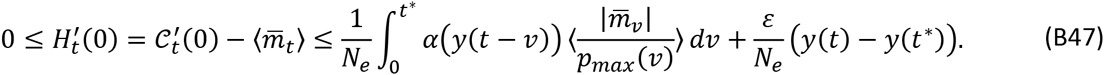

If *α*(*z*) admits a finite positive root *z*_1_, as in the generalized FGM under the WSSM approximation (Appendix E) or in Rouzine et al.’s binary model, we have the upper bound *y*(*t*) ≤ *z*_1_ for all *z*, and *α*(*y*(*t* − *v*)) → 0 as *t* → ∞, for all *v* ∈ (0, *t*\*), which means that the error term

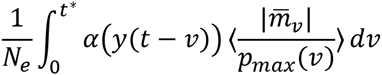

converges to 0 as *t* → ∞ and therefore transient error has no effect on the ultimate deviation. In particular, at large times,

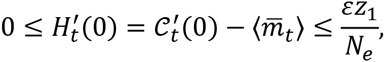

which shows that the error 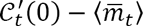 remains bounded by some constant which is independent of *t* and independent of the values taken by 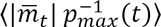 during the transient period (0, *t*\*). The error on the long term behavior has no memory of the error in the past (error does not accumulate).

For log-linear background dependence, below the threshold for *U* where *α* has a finite root, we cannot state the order of the error and must rely only on simulations to check the accuracy of the deterministic approximation.

## Appendix C: Application to non - epistatic models

The results provided in **Appendix B** are of general form: here we first apply them to general context-independent mutation kernels (no epistasis), and look for analytical solutions over time in these cases. This serves first as a check that the model is consistent with results obtained previously, and allows to derive some new insights. We study the trajectory of the fitness distribution during a bout of adaptation from given arbitrary starting conditions (standing variance in fitness), then describe the long term stationary regimes of the fitness distribution (equilibrium or stationary travelling wave).

### I. General analytic solution and properties

#### Solution over time

All non-epistatic models assume that the DFE *f*(*s*|*m*) = *f*_*_(*s*) is independent of the background in which mutations arise. This is characterized by *M*^*s*^(*z,m*) = *M*_*_(*z*) independent of *m*. We thus retrieve a special case of ‘linear-background dependence’ with *α*(*z*) = 1 and *β*(*z*) = *U* (*M*_*_(*z*) − 1), which solution is given in appendix B. The solution to the ODE in Eq. **(B17)** (*y*′ = *α*(*y*) = 1, *y*(0) = 0), is obviously *y*(*z*) = *z* and its inverse is *y*^−1^(*z*) = *z*. The general solution in **(B26)** yields the trajectory of the CGF of the fitness distribution:

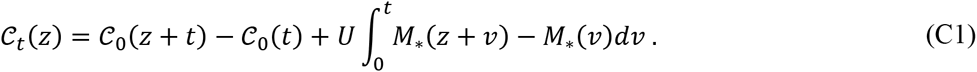

This solution can be computed for any model with analytical MGF/CGF of the initial distribution and of the DFE: be it discrete or continuous, including beneficial or deleterious mutations or both. We note the consistency between the mutational term in **(C1)** and the discrete time version previously found in eq. (13) of (Johnson 1999), and, after a slight rearrangement, that found for continuous time in eq. (10) of (Desai and Fisher 2011). This is expected, as these models also describe the Laplace transform of the fitness distribution under non-epistatic mutation. The main difference with these previous results is that Eq. **(C1)** needs not be limited to a purely deleterious mutation model, and that it allows for arbitrary initial standing variance. We now turn to some further general insight that may be gained by studying the form of the CGF in Eq. **(C1)**.

The form of 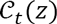 in **(C1)**, as a sum of two CGF terms, implies that the fitness distribution at any time is a sum (convolution) of two independent variables (contributions) generated by two processes. The first contribution is the result of selection acting on pre-existing standing variance, yielding a random component with CGF 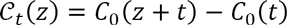. The second contribution is generated by the interplay of mutation and selection and yields a random component with CGF 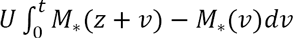. We detail these contributions below.

#### Cumulant trajectory

The cumulant of order *k* at time *t* is obtained by taking the *k*^th^ derivative 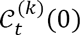(0), with respect to *z*, taken at *z* = 0, of the expression **(C1)**. This yields a general expression for the expected cumulant *c_k_(t)* of arbitrary order *k*, at time *t*:

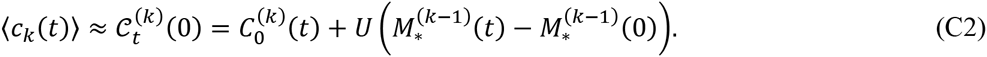

The ′ ≈ ′ is in fact ′ = ′ for the first cumulants 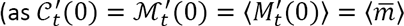. The first term 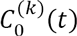 describes the dynamics of adaptation from selection on standing variance (with an arbitrary initial fitness distribution). The second term, proportional to *U*, describes the contribution of new mutations with an arbitrary non - epistatic DFE. The three first cumulants equal the three first central moments of the distribution, e.g. the expectation of the mean fitness 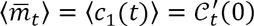 and of the fitness variance 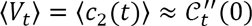 have the following trajectories:

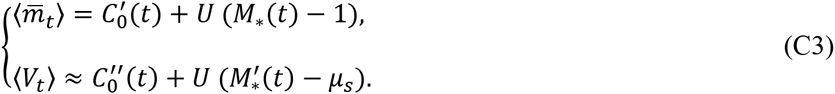

Here 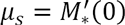 is the mean of the DFE. As expected, the mutational contributions are consistent with Johnson’s (1999) results (eqs. 14-15): allowing for beneficial mutation does not affect the relationship between mutational contribution and the MGF of the DFE.

Trajectories of fitness mean and variance are illustrated in **Fig. C1** for a constant DFE *(s = µ*_*s*_ < 0) and a negative gamma DFE 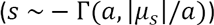. The simulated trajectories, averaged over replicates, are accurately captured by the deterministic theory **(C3)**, and replicates show limited variation around these expectations. Note that all these predictions lose accuracy over much longer timescales (of the order of *N*_*e*_ time units), as Muller’s ratchet starts to play a role in finite populations, see Section III below.

**Fig. C1.**
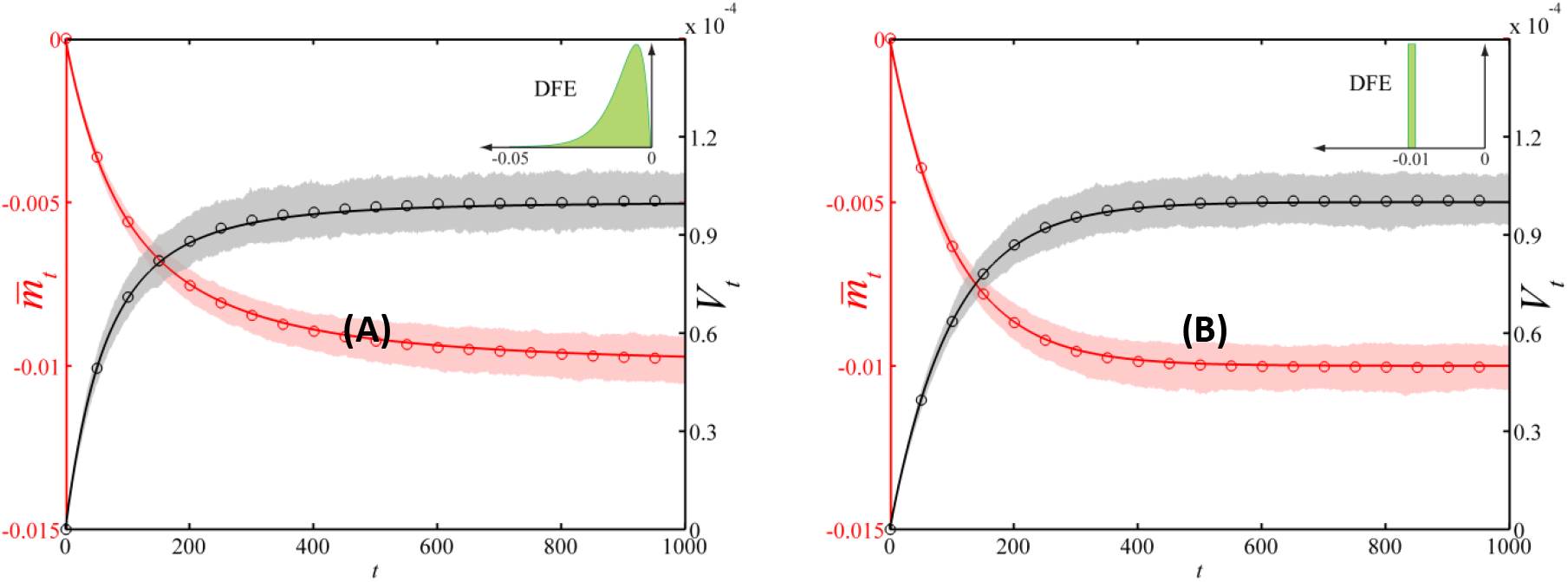
Mean fitness 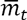 and variance *V*_*t*_ trajectories in non-epistatic models with deleterious mutations. (A) gamma DFE *s* ~ − Γ(a, |*µ_s_|/*a), with shape parameter a *= 2* and scale parameter |*µ*_*s*_|/*a =* 5 · 10^−3^; (B) constant DFE *s* = *µ*_*s*_ = −0.01. Plain lines: trajectories given by the analytical theory (Eq. **[8]** in main text and Eq. **(C3)**); circles: empirical mean fitness and variance given by individual based simulations, averaged over 10^3^ replicate simulations *(N* = *N*_*e*_ = 10^5^); shaded regions: 99% confidence intervals for the mean fitness (in red) and the variance (in gray). We assumed initially clonal populations with *m*_0_ = 0. Sup files **Movie 1A** and **Movie 1B** show the dynamics of the corresponding full fitness distributions.

#### Existence of an equilibrium

Equilibrium of the fitness distribution, here, corresponds to mutation-selection balance; it is characterized by a finite limit 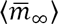, in Eq. **(C3)**, as *t* → ∞. The existence of such equilibrium depends on qualitative properties (i) of the initial fitness distribution and (ii) of the DFE. The mutational term 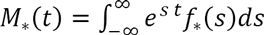 tends to a finite limit as *t* → ∞ if and only if all mutations are deleterious, namely iff *f*_*_(*s*) = 0 for all *s* > 0. Otherwise the term *U M*_*_(*t*) “explodes” over time, as expected if beneficial mutations allow adaptation to increase mean fitness indefinitely. We have also seen that the term 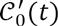 converges to the maximum of the initial fitness distribution: 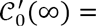 max(*m*_0_). Overall, a simple and intuitive rule applies: a mutation-selection sets in a non-epistatic model whenever (i) there are no beneficial mutations (max(*s*) ≤ 0) and (ii) the initial fitness distribution is bounded on the right (max(*m*_0_) < ∞). We now consider this equilibrium in more detail.

#### Mutation load

We consider only deleterious mutations so that a mutation-selection balance may exist. The maximum of the initial fitness distribution is 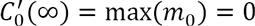, without loss of generality, so that 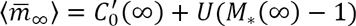 (Eq. **(C3)**). Furthermore, with only deleterious mutations, the MGF term is integrated over 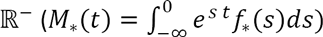, so it vanishes at equilibrium (*M*_*_(∞) = 0). Note that this assumes that there is no discrete probability mass at *s* = 0. This would boil down to neutral mutations, which are not included in the mutational process with rate *U*. Overall, the mutation load *L* is always equal to the mutation rate:

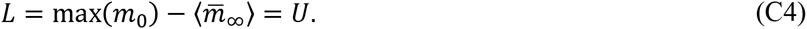

This essentially the continuous time version of Kimura and Maruyama’s (1966) classic result (also known as Haldane-Muller principle) for a discrete time, discrete DFE model: 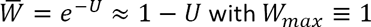. For continuous time models, Bürger and Hofbauer (1994) already obtained Eq. **(C4)** for general non-epistatic models, as a small *U* approximation. The result proves in fact exact for all *U* (under the deterministic approximation).

### II. Stochastic representation of the fitness distribution with arbitrary DFE

CGFs can be fitted directly to data (see Knight and Satchell 1997), but to use the power of a maximum likelihood framework requires knowing the distribution function. This function can be derived numerically as the inverse Laplace transform of 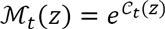, but this method is very sensitive to inaccuracies in the computation of 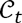. Overall, it may prove useful to derive an explicit pdf, via a ‘stochastic representation’ of the fitness distribution at any time.

We now detail the dynamics of the fitness distribution in more detail by interpreting Eq. **(C1)** in terms of its two contributions.

#### Contribution from standing variance

The term *C*_0_(*z + t*) − *C*_0_(*t*) in Eq. **(C1)** describes the contribution from standing variance. It implies that this is an ‘exponential tilting’, by a factor *t*, of the initial fitness distribution (with CGF *C*_0_(.)): the expected frequency of fitness class *m* at time *t* is 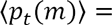 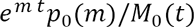. Any starting distribution that is unchanged by exponential tilting (e.g. negative gamma or Gaussian) will remain qualitatively the same over the course of adaptation from standing variance. A Gaussian initial fitness distribution *m*_0_ ~*N*(*µ*_0_,*V*_0_) yields a Gaussian wave *m ~ N*(*µ*_0_ + *V*_0_*t,V*_0_), travelling at constant speed 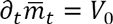. Similarly, a negative gamma initial fitness distribution 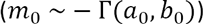 yields a negative gamma wave 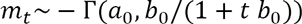 with decreasing speed 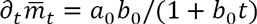.

As expected, this contribution ultimately converges to a Dirac at the fittest class of the preexisting variants, as *t* → ∞ (this maximum class may be infinite, e.g. with a Gaussian wave). Indeed, the CGF *C*_0_(*z* + *t*) − *C*_0_(*t*) converges to 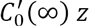, the CGF of a dirac at 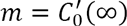, which is the maximum of the distribution with CGF 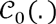.

#### Contribution from mutation

the term 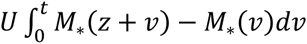 in Eq. **(C1)** describes the contribution from new mutations. Below we detail its form for constant effects, then extend it to arbitrary DFEs and derive simplified approximate forms to this contribution.

<u>Constant effects</u>: Consider first the simplest model: a constant effect of mutations *s = s*_*d*_ < 0 (so that *M*_*_(z) = *e*^*s*_*d*_*z*^). The solution in **(C1)** then yields 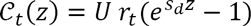 where *r*_*t*_ = 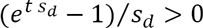. This CGF corresponds to a compound Poisson distribution with stochastic representation: *m* = *n*_*t*_*s*_*d*_, where *n*_*t*_ *~ poissin(U r_t_)*, consistent with the classical result of (Haigh 1978).

<u>Arbitrary DFE</u>: Let us now consider the generalization of this result. Assume now that the DFE is arbitrary, with only deleterious effects 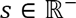 given by the PDF *f*_*_(*s*), and corresponding MGF: *M*_*_(*z*) = 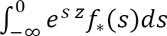. As for all our treatments, this includes the subcase of *k* classes of discrete effects 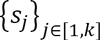 by defining 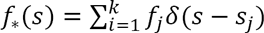 where the *f*_*j*_ are the weights of each class and *δ*(.) is the Dirac delta function. Because Equation **(C1)** is linear in *M*_*_(*z*), the solution 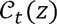 can be written as a weighted sum of constant effects terms (*δC*_*t*_(*z*,*s*) below) contributed by each (potentially infinitesimal) fitness class [*s*,*s* + *ds*]. More precisely, by swapping integrals on *v* and *s*, we can write

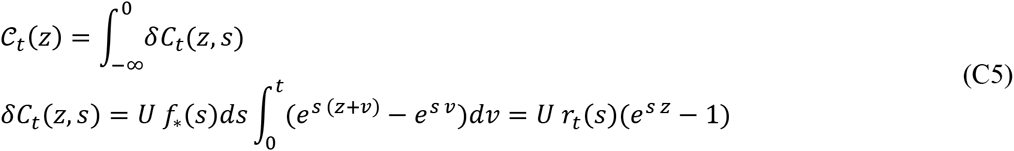

where 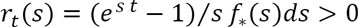. The infinitesimal contributions *δC*_*t*_(*z*, *s*) are the CGFs of compound Poisson random variables of the form *δm*_*s*_ = *n*_*t*_(*s*)*s* where *n*_*t*_(*s*) ~ *Poisson*(*U r*_*t*_(*s*)), so the sum is the CGF of a sum of independent draws from these compound Poisson variables. As such sum is also a compound Poisson distribution, this formulation yields an explicit stochastic representation for *m* as a compound Poisson distribution 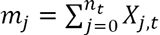, where *n*_*t*_ *~ Poisson*(*U r*_*t*_) with

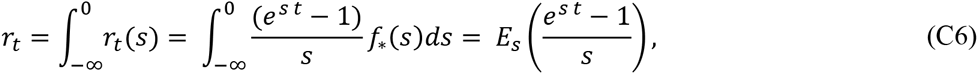

which is always positive ((*e*^*st*^ − 1)/*s* > 0 for all 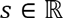). The increments *X*_*j,t*_ are distributed as a mixture of all contributions, namely their probability density function is

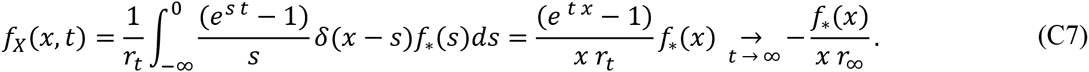

The MGF of the distribution of increments *X*_*j,t*_ for time *t* is

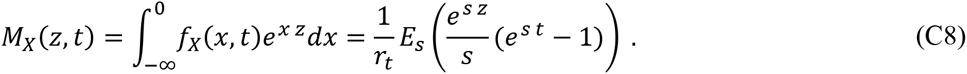

Where the expectation is taken over the DFE. Up to now we have not required mutations to be purely deleterious, we now study the case of deleterious mutations in more detail.

#### Equilibrium fitness distribution with deleterious DFE

Letting time go to infinity, with purely deleterious mutations 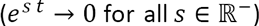 in **(C6)**, the Poisson parameter tends to *U r*_∞_ = *U/s*_*H*_, where *s*_*H*_ = 1/*E*(−1/*s*) is the harmonic mean of the DFE, in absolute value. The increments *X*_∞_ are distributed as *s* weighted by −1/*s* = 1/|*s*|. As should be, we thus retrieve the equilibrium results of T. Johnson (1999) and H.A. Orr (2000), for a set of discrete effects.

#### Small *U/s* approximation

It may be difficult to explicitly write the pdf of a compound Poisson distribution (multiple convolutions of the pdf in Eq. **(C7)**). However, whenever *U r*_*t*_ ≪ 1, the rate of the Poisson *r*_*t*_ remains small at all times, so that it suffices to consider only two fitness classes, the non-loaded class (*m* = 0) with weight 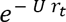 at time *t*, and the single mutation class (*m = X*_*t*_) with weight 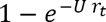 and pdf given by Eq. **(C7)**: the pdf of the fitness distribution is thus

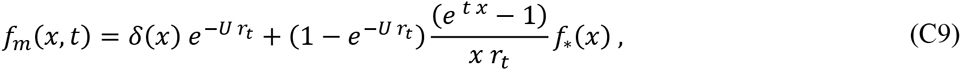

Where *δ*(*x*) is the Dirac delta. This explicit distribution can always be computed with any DFE with known pdf *f*_*_(*x*).

#### Gaussian approximation

On the other hand, the rate *U r*_*t*_ may be large in two situations: with purely deleterious mutations (0 < *r*_*t*_≤1/*s*_*H*_) provided *U*≫ *s*_*H*_, or with beneficial mutations in general conditions after sufficient time has elapsed (as *r*_*t*_ → ∞ then). In this case, the compound Poisson converges to a normal distribution (by application of the central limit theorem), with mean and variance equal to the moments derived in Eq. **(C3)**:

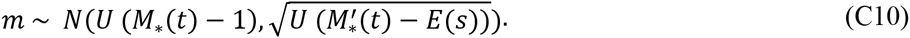

This approximation should be particularly well suited to DFEs that include a substantial portion of beneficial mutations, as the rate *r*_*t*_ then quickly becomes large. On the other hand, it will fail over long timescales as the deterministic approximation fails in this case.

#### Negative Gamma DFE

An important subcase is to consider the gamma DFE (*s* ~ − Γ(*a,b*)), which is widely used to describe deleterious mutations, both theoretically and empirically. It is also of interest as a point of comparison with the Gaussian FGM (see Results and Appendix D). The rate parameter of the Poisson is then *U r*_*t*_ with *r*_*t*_ = (1 − (1 + *b t*)^1–a^)/(*b* (*a* – 1)), which diverges at infinite time whenever *a* ≤ 1 and converges to *r*_∞_ = 1/(*b*(*a* − 1)) whenever *a* > 1.

When *a* > 1, Eq. (C8) at equilibrium yields *M_X_(z*, ∞) = (1 + *b z*)^*a*−1^, namely *X*_∞_ ~− Γ(*a* − 1,*b*) is also a gamma distribution, with smaller shape. In general, the pdf of the resulting Poisson-Gamma distribution can be computed analytically and fitted to empirical data, for any given value of *a* > 1.

The dynamics of the full fitness distribution under deleterious mutation and selection are illustrated in **Movies 1A** (negative gamma DFE) and **1B** (constant DFE), with the same parameters as in Fig. **C1**, but this time comparing theory with results from a single stochastic simulation.

### III. Characteristic timescales for purely deleterious DFEs

#### Timescale of load build-up

Ignoring standing variance, Eq. **(C1)** allows to derive the characteristic time to reach mutation selection balance. The time *t*_*q*_ to reach a high fraction *q* → 1 of the equilibrium mean fitness 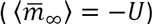 is the solution of 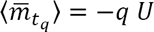, namely *M*_*_(*t*_*q*_) = 1−*q*. This timescale is thus independent of the mutation rate. Let 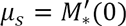 be the mean of the DFE, then by Jensen’s inequality: 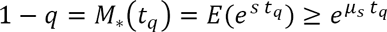 so that 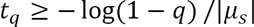. Additionally, *1–q* = 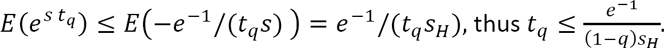. For example, the time to reach 95% of the load is 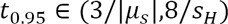.

#### Time to loss of accuracy

As we have seen (Appendix B, part III.2), the error made by the deterministic approximation on the mean fitness, 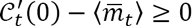, can be bounded from above by 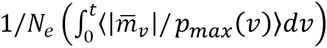. With purely deleterious mutations *r*_*t*_ = *E*((*e^s t^ −* 1)/*s*) is an increasing function of time, so that the frequency *p*_*max*_(*w*) of the fittest class remains above its equilibrium value 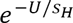 at all times *w* (under the deterministic approximation itself). Additionally, 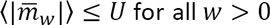. Thus, the error 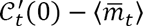 is bounded by 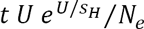. The prediction should remain accurate as long as this error is smaller than the deterministic term (*U* (*M*_*_(*t*) − 1)) which is of order *U*. This implies that the deterministic approximation should remain accurate as long as 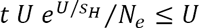, namely at least while 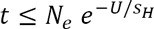 for models that have a non-vanishing *s*_*H*_.

### IV. Connection to stochastic models in the presence of beneficial effects

#### Transition to stationary adaptation

Equation **(C1)** can in principle handle any form of DFE, including ones with beneficial mutation effects, for which equilibrium is impossible. However, simulations and our heuristic treatment of genetic drift show that the model is inaccurate over some timescale, whenever non - epistatic beneficial mutations are present. This can be understood by considering the term that was neglected in the expected MGF dynamics, 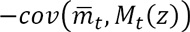 (see Eq. **(A2)** in Appendix A), in particular its contribution to mean fitness dynamics 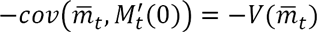 (minus the variance in mean fitness among replicate trajectories). For any non-epistatic model with beneficial mutations, mean fitness can grow indefinitely and 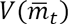 scales with it, while the expected within population variance 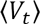 reaches a constant level at stationary regime, so that mean fitness increases at constant rate. Therefore, the between-population variance becomes large relative to the within-population variance, after sufficient time, and our deterministic approach breaks down: it tends to overestimate the fitness increase (as the neglected contribution is 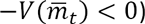.

A more intuitive explanation can be given as follows: in the deterministic PDE, beneficial mutants (even vanishingly rare), create a tail in the fitness distribution that spreads over infinite values, in a vanishingly small time interval due to the continuous time approximation. A wide portion of these mutants are in fact lost by genetic drift, imposing a speed limit to adaptation (a constant rate 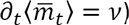), that is neglected here.

This tail starts to impact the dynamics after some time, when *de novo* beneficial mutants start to become dominant in the population, so the transient fitness dynamics are still well approximated by the deterministic PDE. The stationary process that sets later is better handled by stochastic fixation models, in the classic “clonal interference theories”. Yet, a heuristic approach suggests that the fitness dynamics derived here provide a connection between the transient non-stationary dynamics and the ultimate stationary regime of steady fitness increase.

Consider some stationary rate of adaptation *v* > 0, assumed known as a function of (*N,N_e_, U*,*f*_*_(*s*)). In stationary regime, the expected mean fitness increases at constant rate 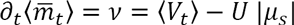 (where 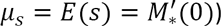. As standing variance affects the system as an independent contribution, we can ignore it in the present argument and study the transition to stationarity of the mutational contribution in **(C1)**. A consistency argument implies that at the transition to stationarity, the deterministic and stochastic rates of adaptation must be equal. From **(C3)**, the deterministic dynamics of the mutational term yields 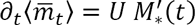 so transition must occur at some time *τ* satisfying

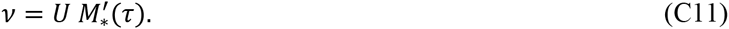

This *τ* can in principle be found (numerically or analytically) for any model with analytic *M*_*_(.) and stationary rate of adaptation *v*. After this point, all higher cumulants must remain stable for the variance to be stationary (a travelling wave solution). They are thus set to those predicted by the deterministic dynamics at time *τ*. The CGF of the fitness distribution, after adding the independent component generated by standing variance, should thus approximately be

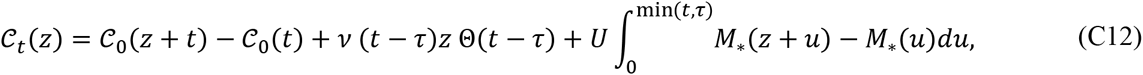

where Θ(·) is the Heaviside theta function: Θ(*x*) = 0 for all *x* < 0 and Θ(*x*) = 1 for all *x* ≥ 0.

#### Exponential DFE

Let us apply this to a classic subcase: an exponential DFE of exclusively beneficial mutations: *s* ~ Exp (1/*µ*_*s*_) with mean *µ*_*s*_ > 0, so that *M_*_(z)* = 1/(1 – *µ*_*s*_*z*) defined on *z* ∈ [0,1/*µ*_*s*_]. The time of transition to stationarity is the first positive root of **(C11)**:

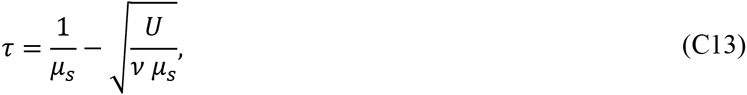

and the corresponding CGF of the fitness distribution, covering non stationary and stationary regimes is

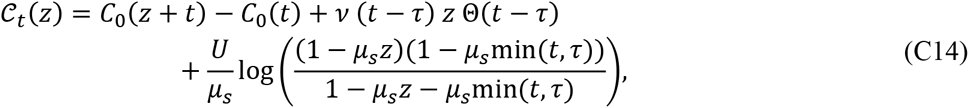

defined over the finite positive domain 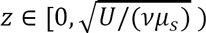. The stationary adaptation rate *v* used in **(C13)** and **(C14)** depends on the mutation rate, the shape of the DFE (Fogle *et al*. 2008) and the demographic processes that determine the stochasticity of the model. For example, consider a discrete generation model with a Wright-Fisher model of genetic drift (Poisson offspring distribution). With mild mutation rates *U*, the stationary rate of adaptation is given by Gerrish and Lenski’s (Gerrish and Lenski 1998) original “clonal interference” theory, which, applied to an exponential DFE, yields

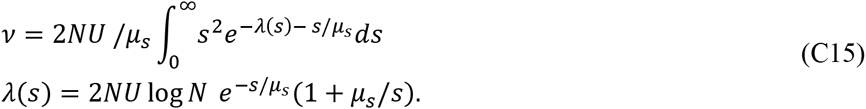

With higher mutation rates (and a continuous time process of birth and death yielding probabilities of establishment equivalent to the Wright-Fisher model), the stationary rate of adaptation is better captured by Desai and Fisher’s “multiple mutation” theory (Desai and Fisher 2007), which, applied to an exponential DFE, (eqs. 15 and 16 in Good *et al*. 2012), yields

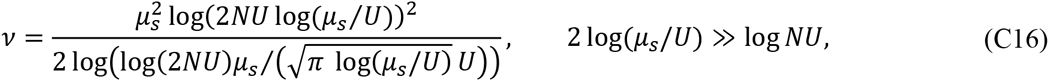

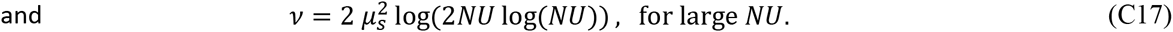

In **Fig. 2** (main text), and **Figs. C2, C3** below, we illustrate this conjecture with various DFEs and parameters. In each case, we check (i) whether the transient expected trajectories (fitness mean and variance) are correctly described up to time *τ* and (ii) whether the average *v* from existing analytic theory (dashed lines) is sufficient or whether the agreement is improved by using *v* inferred from simulation (plain lines).

**Fig. C2.**
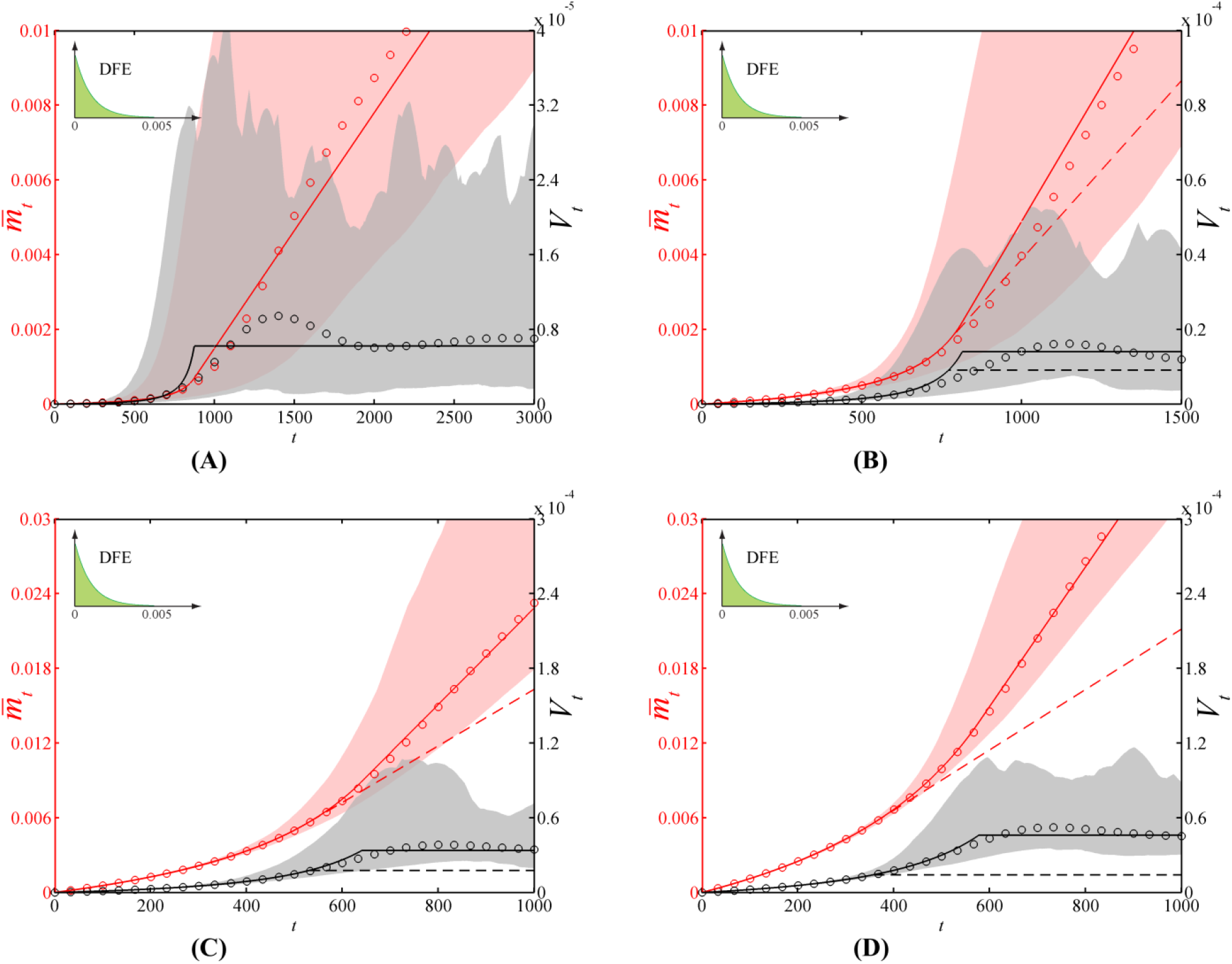
Mean fitness 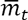 and variance *V*_*t*_ trajectories in non-epistatic models with beneficial mutations. The DFE is given by an exponential distribution *s* ~ *Exp(1/µ*_*s*_) with mean *E(s) = µ*_*s*_ = 0.001 and panels correspond to different mutation rates: (A): *U* = 10^−4^; (B): *U* = 5 10^−4^; (C): *U* = 5 10^−3^; (D): *U* = 10^−2^. The transition time *τ* = 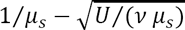 (Eqs. **(C11), (C13)**) is computed for a given stationary rate of adaptation 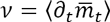 either from stationary regime theory (dashed lines) or observed in simulations (plain lines, *v* averaged over replicates from *t* = 2000 to *t* = 3000). For *t < τ* the expected trajectories 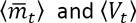 are given by our analytical theory (Eq. **(C3)**). From *t > τ*, the slope 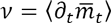 and the variance ⟨*V*_*t*_⟩ are kept constant. Dashed lines: *v* equals the slope given by the most suited theory (i.e., the theoretical slope which is closest to the empirical one: Eq. **(C16)** in panels A and B; Eq. **(C17)** in panels C and D). Circles: empirical mean fitness and variance, averaged over 10^3^ individual based simulations (*N = N*_*e*_ = 10^6^); shaded regions: 99% confidence intervals for the mean fitness (in red) and the variance (in gray). We assumed initially clonal populations with *m*_0_ = 0.

**Fig. C3.**
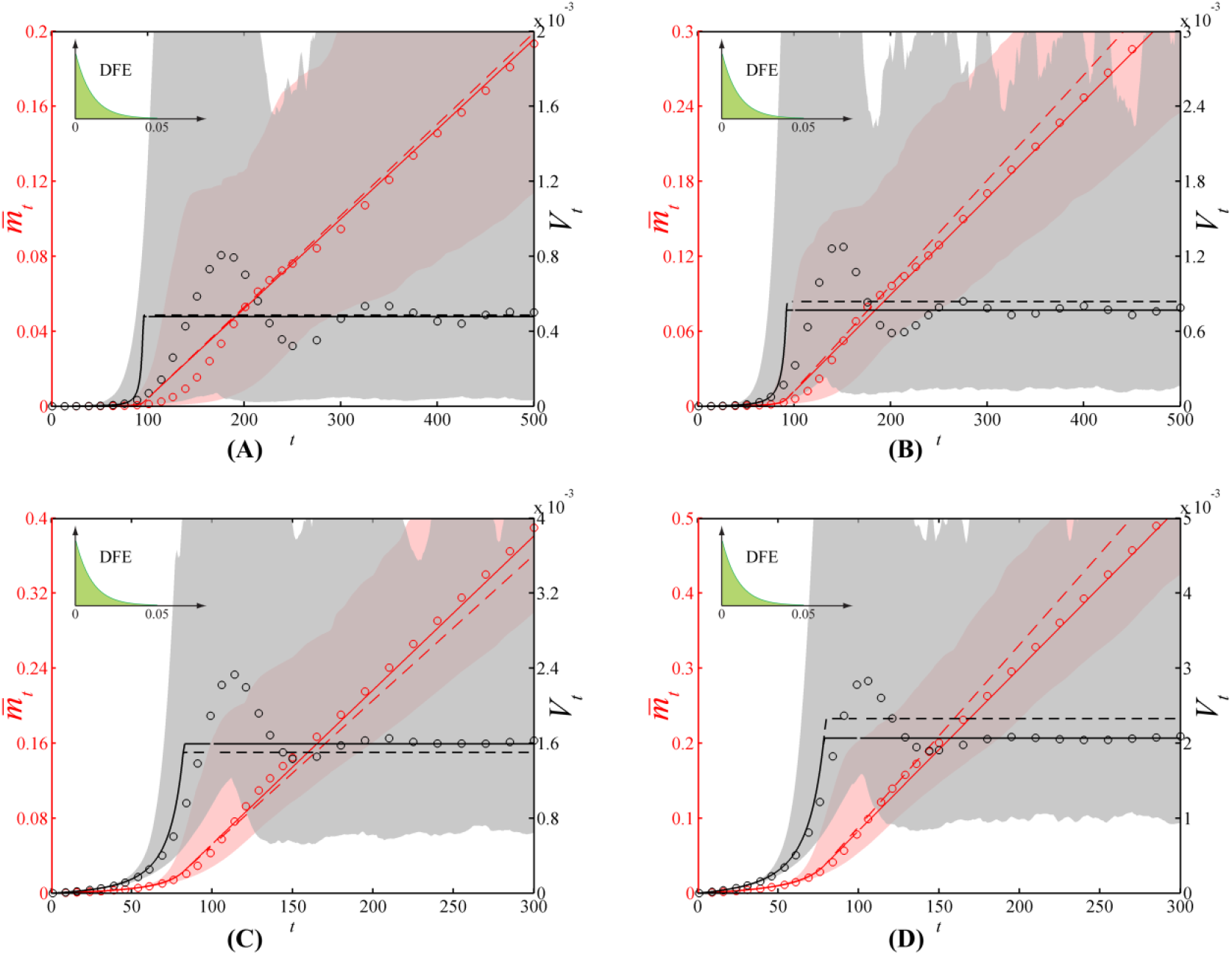
Mean fitness 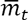 and variance *V*_*t*_ trajectories in non-epistatic models with beneficial mutations. Same as **Fig. C2**, with a higher mean effect of mutation: *E(s) = µ*_*s*_ = 0.01. The theoretical slopes *v* (dashed lines) are given by Eq. (C15) in panel A, Eq. (C16) in panels B and C and Eq. (C17) in panel D.

#### Other DFEs

As a second example, we consider a DFE with constant effects, either deleterious or beneficial: *s* = *δ* > 0 with probability 1/2 and *s =–δ* with probability 1/2. In such case, *M*_*_(*z)* = cosh(*δ z*), and given the stationary rate of adaptation *v*, the time of transition to stationarity is *τ* = 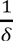 arcsinh 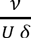. The corresponding mean fitness and variance trajectories obtained by connecting the analytical expressions of Eq. **(C3)** with the stationary adaptation regime are compared with individual-based simulations in **Fig. C4.**

Lastly, **Fig. C5** shows trajectories corresponding to a displaced gamma DFE, which is continuous and includes both deleterious and beneficial mutations. Here, *s* ~ *s*_0_ − *x*, with *s*_0_ > 0 and x ~ Γ(*a, b*) (in this case, 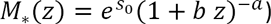. The time of transition to stationarity is obtained numerically here, for a given *v*.

**Fig. C4.**
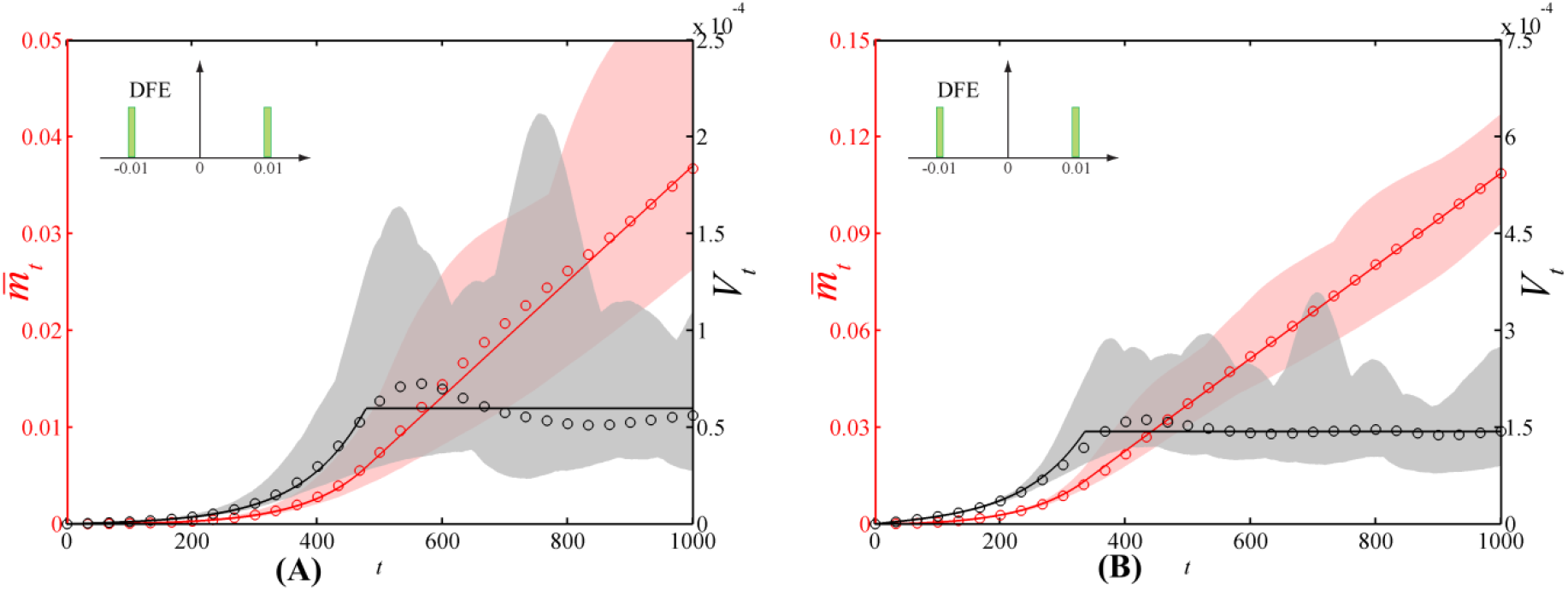
Mean fitness 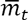 and variance 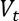 trajectories in non-epistatic models with beneficial and deleterious mutations. Same as **Figs. C2–C3**, but with 10^2^ replicate simulations and another DFE, consisting of symmetrical constant effects: *s* = *δ* > 0 with probability 1/2 and *s =−δ* with probability 1/2, and *δ* = 0.01. (A): *U* = 10^−4^; (B): *U* = 10^−3^. Transition to stationarity at *τ* = arcsinh(*v*/(*U δ*)) /*δ*, with rate *v* inferred from simulations as a measured stationary rate 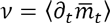 averaged over replicate simulations from *t* = 2000 to *t* = 3000 (no analytical expression available).

**Fig. C5.**
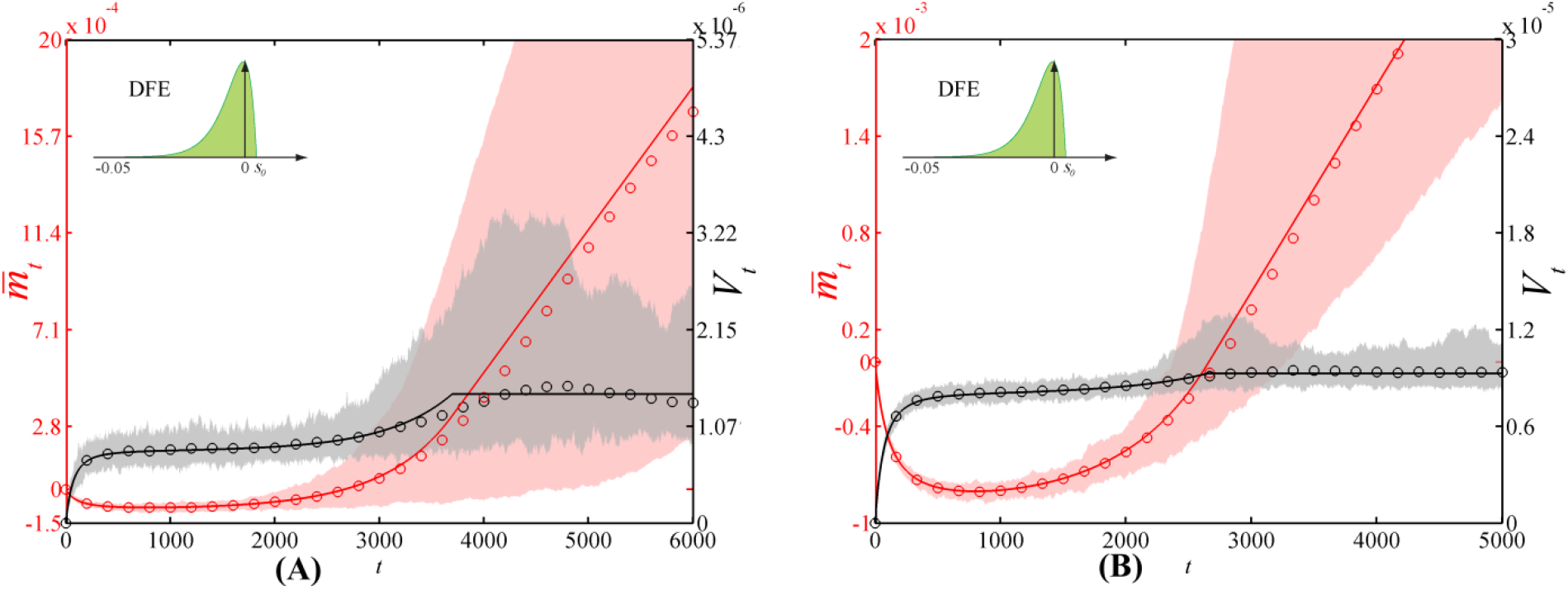
Mean fitness 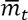 and variance *V*_*t*_ trajectories in non-epistatic models with beneficial and deleterious mutations. Same as in **Fig. C4**, with a shifted gamma DFE: *s ~s*_0_+*x*, with *s*_0_ > 0 and *x* ~ −Γ(*a,b*), with *a* = 2, *b* = 5 · 10^−3^ and *s*_0_ = a *· b*/5. (A): *U* = 10^−4^; (B): *U* = 10^−3^. The time *τ* is obtained by numerically solving Eq. **(C11)** with a stationary rate 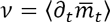 inferred from simulations from *t* = 4000 to *t* = 6000.

## Appendix D: Equilibrium and numerical computations for the ‘Gaussian Fisher’s geometrical model’

### I. Definition of the “Gaussian FGM”

Fisher’s geometrical model (FGM) is our landmark example of context-dependent DFEs. In all versions of FGM considered in this article, Darwinian fitness *W*(**g**) is a Gaussian isotropic function in *n* dimensions, and/or Malthusian fitness is a quadratic isotropic function: 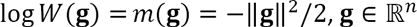. The particular version studied in this appendix arises when the mutation effects on phenotype follow an isotropic multivariate Gaussian distribution (**dg** ~ *N*(**0**, *λ* **I**_*n*_)) where **I**_*n*_ is the identity matrix in *n* dimensions and *λ* > 0 is a positive scaling constant. We denote this particular model the ‘Gaussian FGM’. In the ‘Gaussian FGM’, the MGF of the DFE in a background with arbitrary fitness *m* ≤ 0 is (Martin 2014)

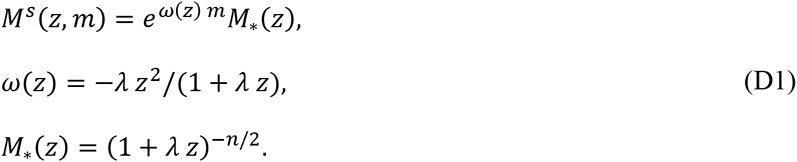

In this case the DFE from an optimal background is gamma distributed (*s*_*_~ − Γ(*n*/2,*λ*)) with MGF *M*_*_(*z*). It is noteworthy that the MGF of *s* is log-linear with background fitness *m* in the Gaussian FGM: the log-linear background-dependence assumption is satisfied and the PDE:

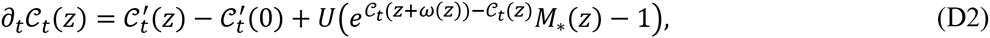

corresponding to Eq. **[7]** in the main text and **(B1)** in Appendix B is exact in the Gaussian FGM. This PDE can be solved numerically, see Paragraph "Numerical methods" at the end of this appendix, leading to accurate description of fitness mean and variance trajectories (**Figs. 3A** and **3B** in the main text). The mathematical properties of this PDE have been analyzed in Appendix B (Section I), under an assumption which is readily satisfied in the Gaussian FGM:

Assumption **H** (appendix B) is obviously verified as any background can mutate to the optimum; it can also be checked that the mathematical counterpart of **H** is verified: 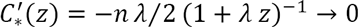 as *z* → ∞ and *ω*′(*z*) = (1 + *λz*)^−2^ − 1 → −1 as *z* → ∞.

Some exact results regarding the equilibria (e.g. memoryless property) of this PDE are derived in Appendix B (Sections I.2 and I.3). However, more general insight is gained via a simple approximation. Near equilibrium (*m* → 0), log-linear background-dependence becomes approximately linear, so the equilibrium solution of the PDE **(D2)** can be approached by the corresponding solution of the linear PDE:

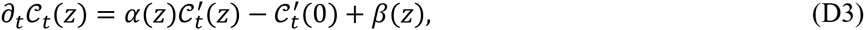

corresponding to Eq. **[4]** in the main text, with the coefficients:

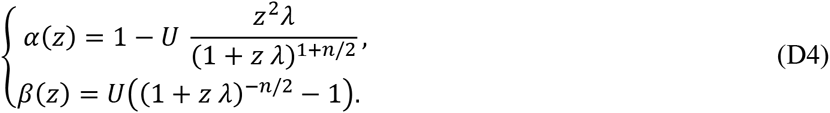

The general equilibrium properties of this linear PDE are detailed in Appendix B (Section II); we apply these results to the particular functional coefficients above to obtain the equilibrium fitness distribution in Fisher’s model.

### II. Dynamics of the fitness distribution

Away from equilibrium, a general explicit solution to **(D2)** could not be found. We thus rely either on (i) a numerical solution (detailed below) or (ii) an analytical weak selection strong mutation approximation (detailed in Appendix E).

### III. Mutation load and spike at the optimum

As seen in Appendix B, the key to derive the mutation-selection balance and mutation load is the first positive root *z*_1_ of *α*. This root, when it exists, can be computed numerically or analytically (at least when *n* = 1, 2,4 or 6). A sign analysis of *α* over 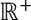 (Eq. (D4)) shows the following rules for the existence of a finite positive root of *α*, depending on the value of *n*, and on the value of the mutation rate *U* with respect to a critical threshold *U_c_:*

(i): *n* = 1: *α*(*z*) decreases monotonically from *α*(0) = 1 to *α*(∞) = –∞ and there is always a positive root 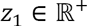. The exact load can be computed explicitly but has a complicated closed form. The results of Appendix B imply that there is no spike at the optimum.

(ii): *n* = 2: *α*(*z*) decreases monotonically from *α*(0) = 1 to *α*(∞) = 1 – *U/λ* ; whenever *U* > *U*_*c*_ = *λ*, there is a finite positive root 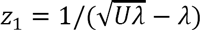 and the load is 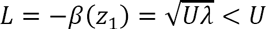, otherwise *z*_1_ = ∞ and *L* = *U*. In the latter case, a spike may exist at the optimum in principle, but its upper bound is 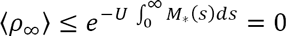 as the integral 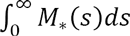 does not converge with *n* = 2 (*M*_*_(*z*) = (1+*z λ*)^−1^).

(iii): *n* ≥ 3 : *α*(*z*) reaches a minimum at *z*_*min*_ = 2/(*λ*(*n*/2 − 1)), equal to *α*(*z*_*min*_) = 1 − 4 *U/* 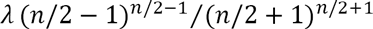. A finite positive root exists iff this minimum is below zero, namely whenever

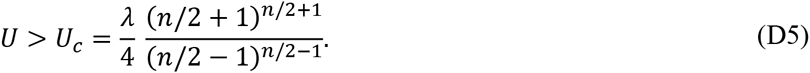

The root *z*_1_ can then be computed numerically or analytically (e.g., 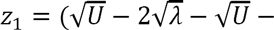 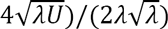 in the case *n* = 6 presented in **Figs. 3A** and **3B**). The results of Appendix B show that the corresponding mutation load is *L* = −*β*(*z*_1_) = *U*(1 −*M*_*_(*z*_1_)). Otherwise whenever *U ≤ U_c_: z*_1_ = ∞, the load is *L = U*, and a spike of optimal genotypes exists in this case, with weight close to (and below) 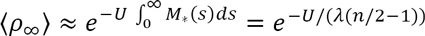.

Note that the general formulae for *n* ≥ 3 handle in fact the case *n* = 2: simply taking the limit for *n* → 2 of *U*_*c*_ in Eq. (D5) yields *U*_*c*_ = *λ* and ⟨*ρ*_∞_⟩ → 0.

### IV. Equilibrium fitness distribution

The CGF of the fitness distribution at equilibrium is obtained by setting *L = U*(1 − *M*_*_(*z*_1_)) and 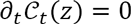 in Eq. **(D3)**: 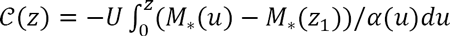, where *M*_*_ and *α* and are given by Eqs. **(D1)** and **(D4)**, respectively. This yields the general expression (for *n* ≥ 3):

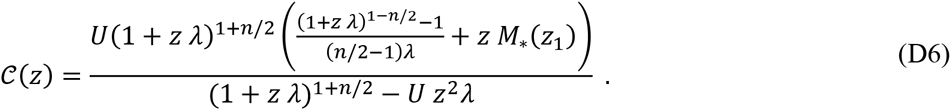

Below the phase transition (*U* <*U*_*c*_) we know that *z*_1_ = ∞ and *M*_*_(*z*_1_) = *M*_*_(∞) = 0, and the equilibrium CGF simplifies to

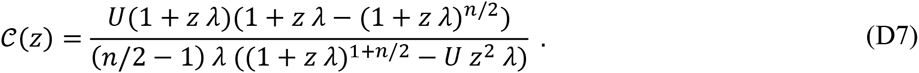

From this expression, it can be shown 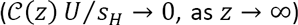 that the estimation of the expected spike weight which is derived for small *U* in Appendix B, Section II.4, is in fact *exact* in the Gaussian FGM for all *U* < *U*_*c*_:

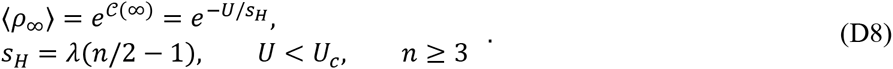

The fitness distribution among suboptimal genotypes has MGF given by 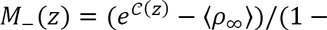 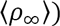 where *C*(*z*) is given by Eq. **(D7)** and *ρ*_∞_ by Eq. **(D8)**. Its expression is complex, but a leading order in small *U* (relevant here as we are below the phase transition), yields *M*_(*z*) = (1+*λ z*)−^(*n*/2−1^), namely the MGF of a negative gamma distribution: *m*_ *~ − Γ*((*n* – 2)/2, *λ*). This is exactly the result expected, as *t* → ∞, from the small *U/s* approximation in a non epistatic model (see Appendix C, Eq. (C9)) with context-independent DFE given by that at the optimum (*s* ~− *Γ*(*n*/2,*λ*)). Therefore, to leading order in *U* (approximately for any *U* < *U*_*c*_), the equilibrium fitness distribution in the FGM is blind to the presence of epistasis and behaves as the equivalent non-epistatic model with gamma DFE.

Beyond the phase transition (*U > U*_*c*_), an exact treatment is more involved as we do not have a general expression for *z*_1_. A weak selection strong mutation treatment, detailed in Appendix E, proves surprisingly accurate in this regime.

### V. Numerical methods

The numerical computation of the solution of the nonlinear PDE **(D2)** was based on a finite difference method with variable step sizes in *z* (smaller steps near *z* = 0, to get accurate values of the derivatives 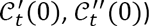 and an implicit scheme in time. The nonlinearity was dealt with using a Newton-Raphson algorithm. The values of the functions at the positions *z* + *ω*(*z*), which generally do not belong to the finite difference mesh, were computed by linear interpolation with the closest positions in the mesh.

Because of the transport term 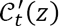, which tends to translate the solution towards the left and with speed 1, the solution was computed on a finite interval *z* ∈ (0,*z*_*max*_) where *z*_*max*_ = *T*, the final time of the computations. The approximation of the solution of **(D2)** on 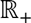 by a solution on a bounded interval was made possible thanks to the property 0 ≤ *z* + *ω*(*z*) ≤ *z*, which ensures that all the positions where 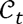 has to be evaluated belong to the interval (0,*z*_*max*_) as long as *z* belongs to this interval. Using the property **(B3)** of Appendix B it was natural to impose the Neumann boundary condition 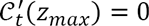. The Matlab^©^ source code of the solver is available as supplementary material, together with a Matlab^©^ graphical user interface (**Fig. D1**). Examples of numerical computations are given in **Figs. 3A**, **3B** (main text) and **Figs. D2** and **D3** (below). Notice that in the top left panels discrepancies with the theory arise for *U* = 0.0002: as expected, (see ‘Convergence to the deterministic approximation’, main text) these correspond to situations where *N*_*e*_*U* |*µ*_*s*_| < 1.

**Fig. D1.**
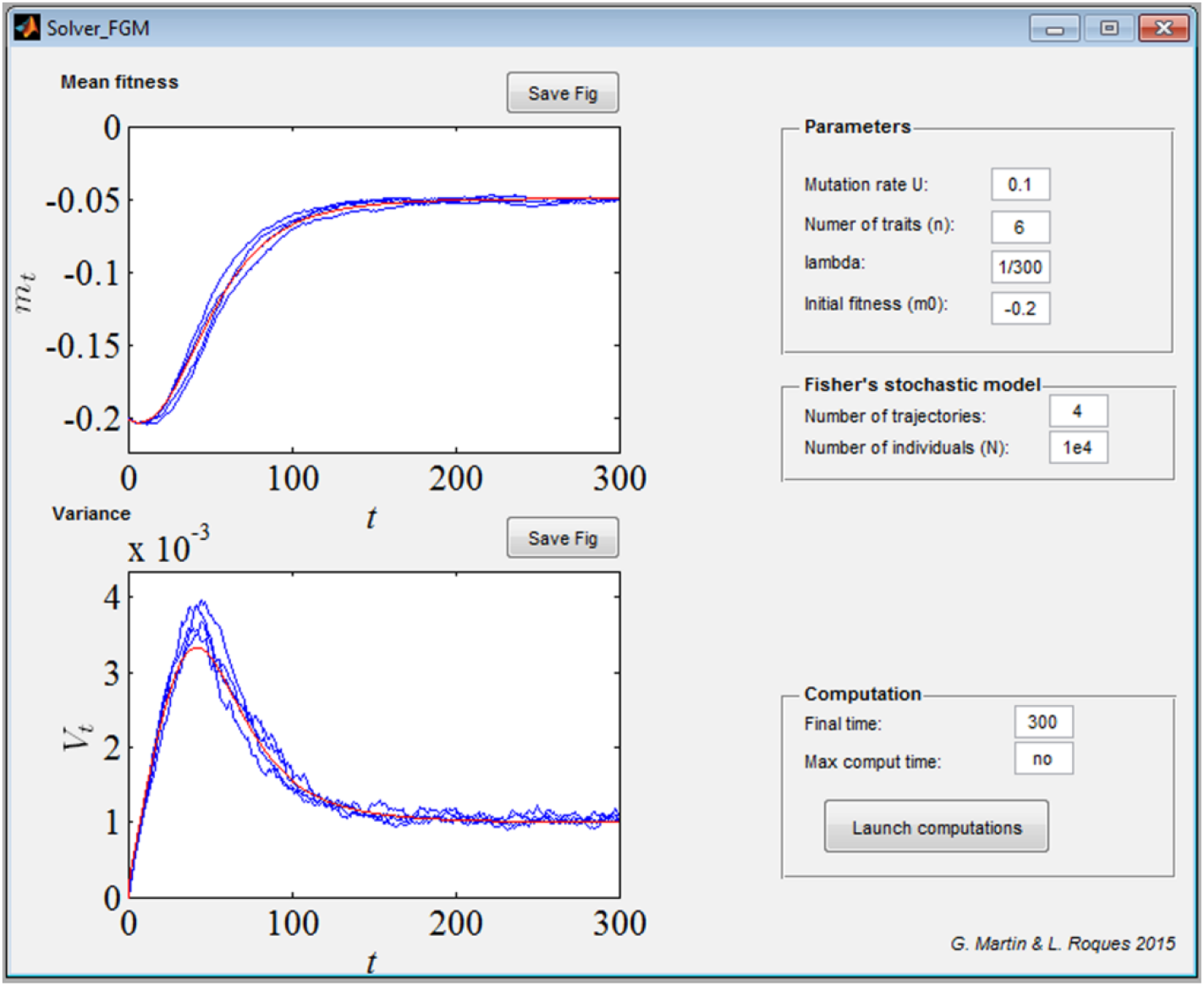
Snapshot of the graphical user interface of the Matlab^©^ solver for the numerical computation of the solution of the nonlinear PDE **(D2)** (Eq. **[7]** in the main text).

**Fig. D2.**
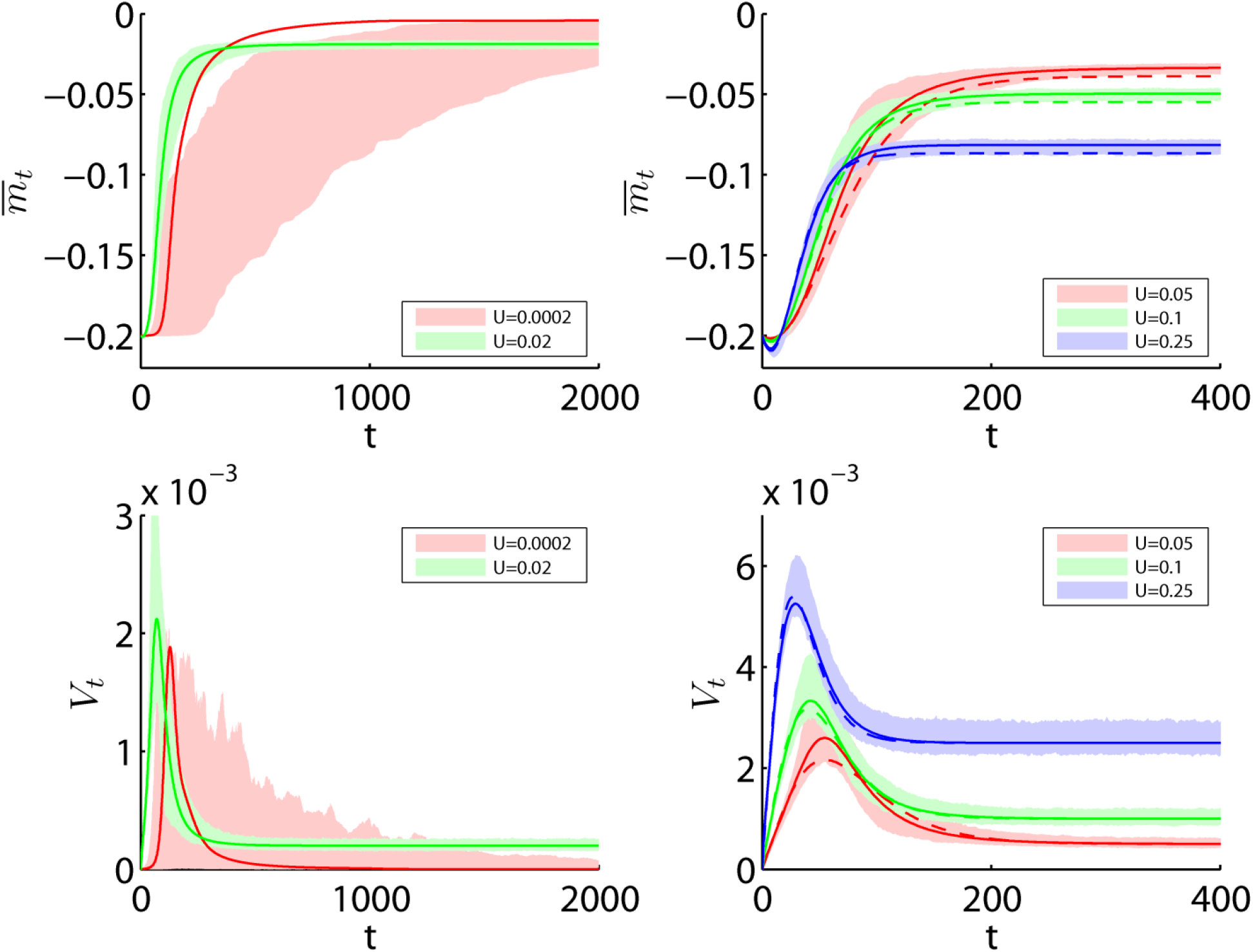
*N = N*_*e*_ = 10^4^ individuals. Mean fitness 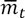 (top panels) and variance *V*_*t*_ (bottom panels) trajectories in Gaussian Fisher’s geometrical model with several values of the mutation rate. Plain lines: expected trajectories 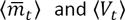 given by the numerical solution of the PDE **(D2)** (or **[7]** in the main text), with *M*_*_ and *ω* as in Eq. **(D1)**. Dashed lines (right panel): expected trajectories given by the analytical solution of the linear PDE **[4]** under the weak selection strong mutation (WSSM) approximation (see Appendix E). The shaded regions correspond 99% confidence intervals given by individual based simulations, with 10^3^ populations of *N* = *N*_*e*_ = 10^4^ individuals. The parameter values are *n* = 6 traits and *λ* = 2|*µ*_*s*_|/*n* = 1/300 (|*µ*_*s*_| = 0.01), leading to a critical mutation rate *U*_*c*_ = 16*λ* ≃ 0.05. We assumed initially clonal populations with *m*_0_ = −20|*µ*_*s*_| = −0.2.

**Fig. D3.**
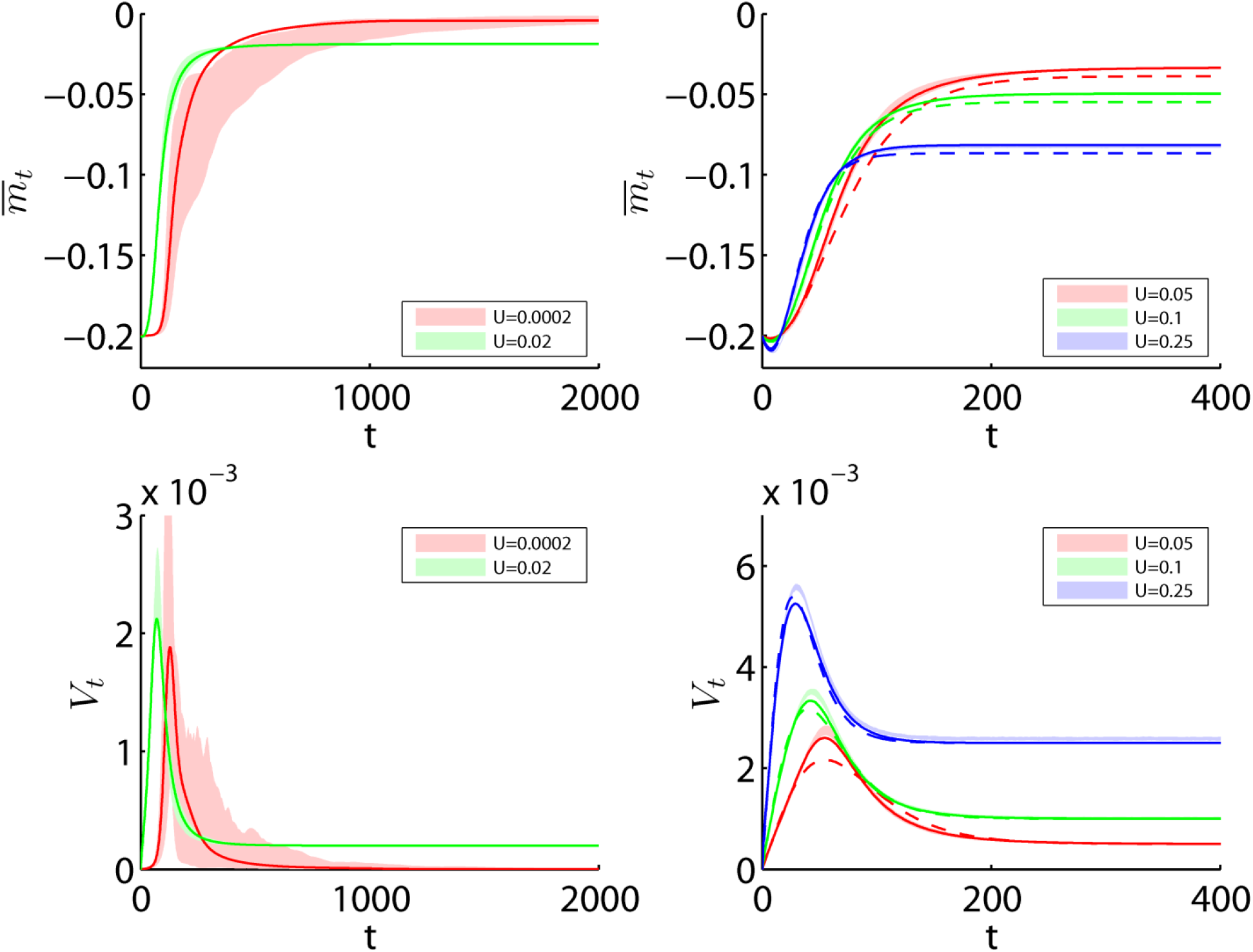
*N = N*_*e*_ = 10^6^ individuals. Mean fitness 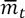 (top panels) and variance *V*_*t*_ (bottom panels) trajectories in Gaussian Fisher’s geometrical model with several values of the mutation rate. Plain lines: expected trajectories 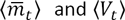 given by the numerical solution of the PDE **(D2)** (or **[7]** in the main text), with *M*_*_ and *ω* as in Eq. **(D1)**. Dashed lines (right panel): expected trajectories given by the analytical solution of the linear PDE **[4]** under the weak selection strong mutation (WSSM) approximation (see Appendix E). The shaded regions correspond 99% confidence intervals given by individual based simulations, with 10^3^ populations of *N* = *N*_*e*_ = 10^6^ individuals. The parameter values are *n* = 6 traits and *λ* = 2|*µ*_*s*_|/*n* = 1/300 (|*µ*_*s*_| = 0.01), leading to a critical mutation rate *U*_*c*_ = 16*λ* ≃ 0.05. We assumed initially clonal populations with *m*_0_ = −20|*µ*_*s*_| = −0.2.

## Appendix E: Generalized FGM

Here we derive explicit results for a particular class of background - dependent models: Fisher’s Geometrical model (FGM). We extend our results from the ‘gaussian FGM’ (the standard model of quantitative genetics) where mutation effects on traits are normally distributed and which is analyzed in appendix D. We derive approximate results (both equilibria and dynamics) for a generalization of this model that relaxes the normality assumption for mutation effects.

**“Generalized FGM”:** We consider here an extension of the Gaussian FGM (Appendix D) to more general phenotypic distributions. Darwinian (resp. Malthusian) fitness is still a Gaussian (resp. quadratic) isotropic function of 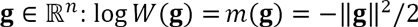. However, the distribution of mutation effects on phenotype may now pertain to a broad class of distributions, with the only constraint that it be spherically symmetric and continuous in the vicinity of **dg** = 0 (the latter to satisfy **H**, see Appendix B). Note that we may set that the distribution has no spike at **dg** = **0** without loss of generality, as *U* is the *non-neutral* mutation rate. We denote this model the “generalized FGM” as it is characterized by fairly general mutant phenotype distributions.

### I. Functions *ω*(.) and *M*_*_(.) in the generalized FGM

The key to apply our approach to a particular model is to derive the functions *M*_*_(*z*) and *ω*(*z*) for this model. We derive these functions in the generalized FGM below. The stochastic representation of the DFE, from a background with phenotype g is 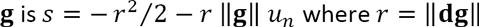 is the norm of the mutant phenotypic effect, and *u*_*n*_ is the cosine of the angle between **dg** and **g**. For any spherically symmetric distribution of **dg**, *u*_*n*_ has MGF *M*_*u*_(*z*) = _0_*F*_1_ (*n*/2, *z*^2^ /4) (see e.g. (Martin and Lenormand 2015)), where _0_*F*_1_(.) is the regular confluent hypergeometric function. Therefore, the DFE among those mutants with fixed magnitude *r*, has MGF 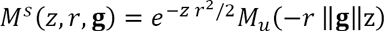. This can be rewritten in terms of the background fitness 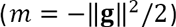 and the fitness effect that the mutants with magnitude *r* would have, in an optimal background (*s*_*_ = −*r*^2^/2):

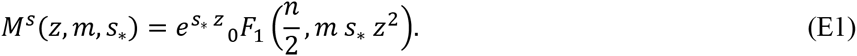

As required, at *m* = 0, the DFE for the mutant class with fitness *s*_*_ has MGF 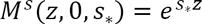 (a Dirac at *s* = *s*_*_). Taking the expectation of Eq. **(E1)** over *s*_*_, we retrieve the MGF of the DFE in background 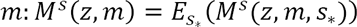. The MGF of the DFE at *m* = 0 is 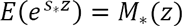, as required.

### II. Equilibrium in the Generalized FGM: application to an exponential DFE

In order to compute general results on the equilibrium fitness distribution (see main text), we must further derive the function 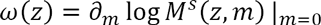. We note from **(E1)** that 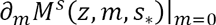 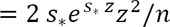 and use the exchangeability of expectation and derivation to get

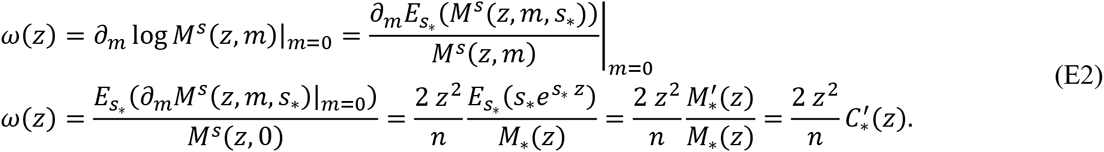

Note that, as should be, Eq. **(E2)** retrieves the correct *ω*(*z*) = *–λ z*^2^(1 + *λ z*)^−1^ for the Gaussian FGM. The two corresponding functional coefficients *α*(.) and *β*(.) are given by

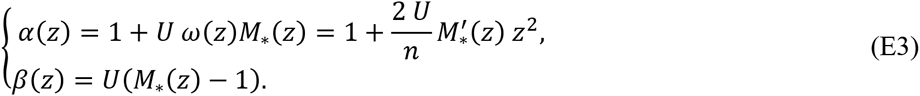

From a given model for the distribution of **dg**, the framework can be applied to compute the corresponding equilibrium of the fitness distribution. The load is given by *L = U*(1 − *M*_*_(*z*_1_)) where *z*_1_ is the smallest positive root of *α*(*z*) = 1 + 2 *U/n* M*′*_*_(*z*) *z*^2^, or *z*_1_ = ∞ if *α*(*z*) > 0 for all *z* ≥ 0.

As an example, consider that **dg** is such that the DFE is exponential at the optimum: *s*_*_~ − *Exp*(1/|*µ*_*s*\*_ |) with mean 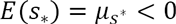, with MGF 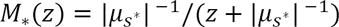. Assuming arbitrary number *n* of dimensions, this model differs from the Gaussian FGM unless *n* = 2. Define *λ* = 2|*µ*_*s*\*_|/*n*, the variance of the phenotypic distribution at each trait (for consistency with the Gaussian FGM). If *U* ≤ *U_c_ = n*^2^*λ*/4, there is no root to *α* and the load is then *L* = *U*. Beyond *U* > *U*_*c*_, *α* has a unique positive root *z*_1_ = 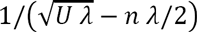 and the load is 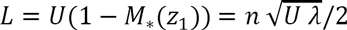. Over all possible *U* values, the load can thus be written: 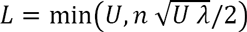.

We now turn to more general results, independent of the details of the distribution of mutant phenotypes (hence of *M*_*_), in a weak selection limit.

### III. Weak selection strong mutation (WSSM) approximation

We note that in Eq. **(E1)**, *s*_*_ enters the function *M*^*s*^(*z, m, s*_*_) in product with *z*. This implies that mutation effects in background *m* scale with their counterparts in background *m* = 0 (with the norm of the mutant phenotypic effect). Taking a leading order in *z s*_*_ yields

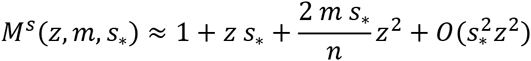

Taking expectations over the distribution of *s*_*_ yields a mutational kernel of linear background-dependence form: 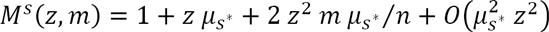, where we require that the coefficient of variation of *s*_*_ be of order 1 (or equivalently 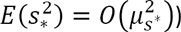. Computing the weighted sum over the within population distribution of *m* yields 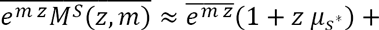 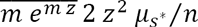. Noting that 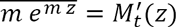 and taking the ensemble expectation yields 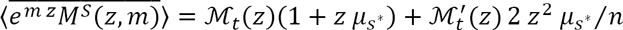, once plugged into Eq. **[2]** yields the mutational term:

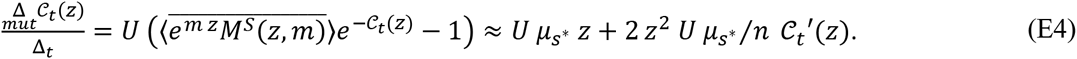

And (as any form of linear background-dependence) a linear PDE for *C*_*t*_ (see Eq. **[4]**):

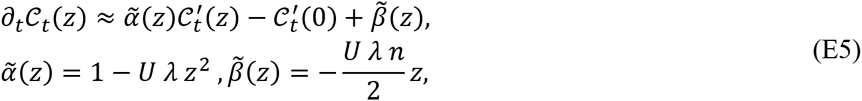

where we recall that λ = 2 |*µ*_*s*\*_|/*n* is the variance of the phenotypic distribution at each trait. Note that the above expressions of 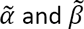 correspond to series expansions at *λ z* = 0 of the coefficients *α, β* proposed in Appendix D for the Gaussian FGM.

#### Range of validity

The approximation is valid to leading order in |*µ_s*_ | z*; therefore, it cannot be accurate over the full range of 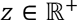, but remains accurate in some finite range *z* ∈ [0,*ϵ*/|*µ*_*s*\*_|] where *ϵ* ≪ 1. This reflects the fact that it does not capture the right tail of the DFE (fitter mutants), this tail being determined by the values of *M*^*s*^(*z, m*) at large *z*. Such limited range implies that the mutation rate must be strong enough relative to mutation fitness effects that the fitness dynamics are not driven by this tail: this is a strong mutation weak selection regime. More precisely, under the approximation, the range of *z* where the solution to Eq. **[4]** evolves is bounded by 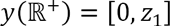 (see Eq. **[5]**). In Eq. **(E5)**, the first positive root of 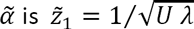: consistency thus implies that a sufficient condition for the solution of Eq. **(E5)** to capture the full dynamics over time is that 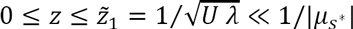. This boils down to a lower bound on the mutation rate, relative to the strength of selection: the weak selection approximation is valid when

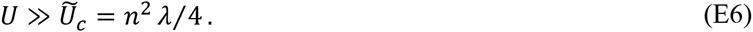

Note here that we have used the notation 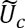 (recalling the critical mutation rate *U*_*c*_ in Appendix D) on purpose. It is also the critical mutation rate where the phase transition occurs between *L = U* and *L < U*, as computed from Eq. **(E5)**. We have (beyond the phase transition) the load 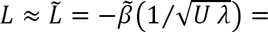 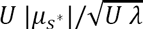 which is exactly equal to 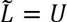 (below phase transition) at the transition point 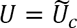.

#### Equilibrium fitness distribution

We know that the mutation load is approximately 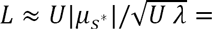 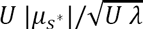, whenever the approximation applies. We also know that, outside this regime, there must be a lower bound to *U* below which *L = U* (with *L < U* beyond this bound). A natural approximation connecting all the range of *U* is

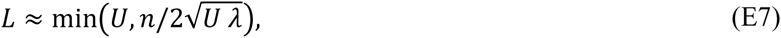

with a phase transition at 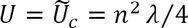. This ‘rule of thumb’ happens to yield the exact behavior for a particular form of FGM (detailed above) with an exponential distribution of *s*_*_, also corresponding to the Gaussian FGM in *n* = 2 dimensions.

The equilibrium fitness distribution has CGF 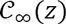 satisfying 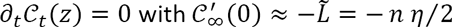, which (from Eq. **(E5)**) yields:

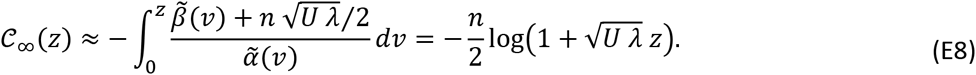

This is the CGF of a negative gamma distribution 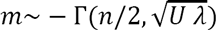.

#### Normal equilibrium trait distribution

Because the FGM links fitness and phenotype, the equilibrium fitness distribution corresponds to a given multivariate distribution of the *n* traits. As fitness is a quadratic function of breeding values for the traits 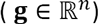, it is easily shown that the gamma distribution of fitness (Eq. **(E8)**) implies a multivariate normal (*MVN*) distribution of the breeding values **g** at equilibrium, with mean at the optimal phenotype 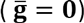 and variance 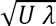 on each trait: 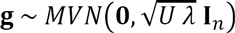, where **I**_*n*_ is the identity matrix in *n* dimensions. Therefore, our weak selection approximation exactly matches Kimura’s (1965) and Lande’s (1980) normal approximations for trait distributions at mutation - selection balance under stabilizing selection. Indeed, although obtained in strikingly different manners, these two approaches rely on the same assumption of strong mutation relative to selection (Eq. **(E6)**). It is also maybe more straightforward here (although already noted by Lande) that this equilibrium is mostly independent of the underlying distribution of mutation effects on phenotypes (generalized FGM).

Beyond its application to fitness, the present treatment thus clarifies that the well-known normal approximation for traits applies, at equilibrium, under two explicit quantitative conditions. First, the mutation rate must be well above 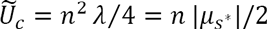. Second, the distribution of mutant phenotypic effects must yield a DFE, at the optimum, that has a coefficient of variation of order 1 or less 〈*CV*(*s*_*_) = *O*(1)〉.

#### General fitness distribution dynamics

The solution of Eq. **(E5)** over time (Eq. **[5]** and Appendix B) depends on the solution of the ODE 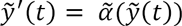 with boundary condition 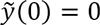. This yields 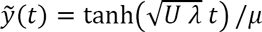 with functional inverse 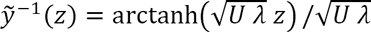. From a given initial fitness distribution characterized by some CGF *C*_0_(*z*), and plugging the particular functions 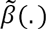, 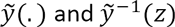 into the general solution in Eq. **[5]**, the CGF of the fitness distribution at time *t* is

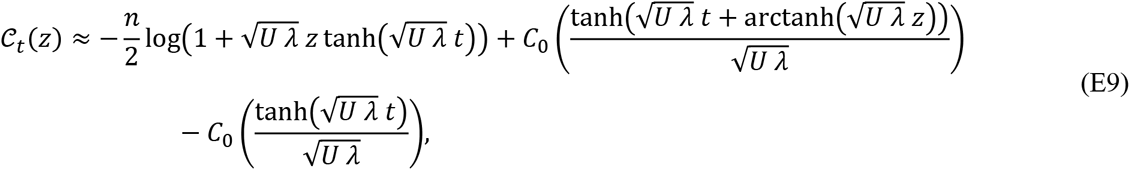

with mean fitness trajectory given by (Eq. **[8]**):

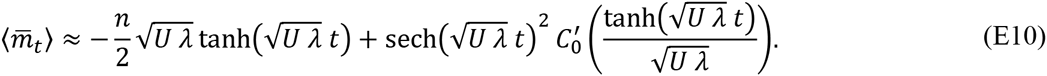

Particular forms of *C*_0_(.) can be studied to obtain explicit forms, depending the on initial conditions: we consider two standard scenarios.

#### Adaptation from a clone

If the population is initially clonal, with fitness *m*_0_ < 0, then *C*_0_(*z*) = *m*_0_*z* and the CGF of the fitness distribution becomes

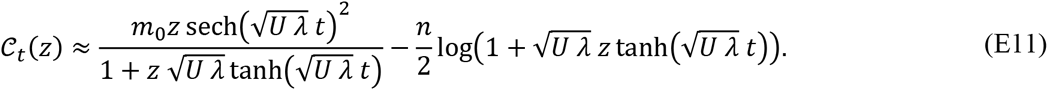

The mean fitness is given by Eq. **(E10)**, with 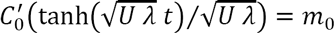. Eq. **(E11)** can be equated to the CGF of a known distribution, providing a stochastic representation for the fitness distribution over time:

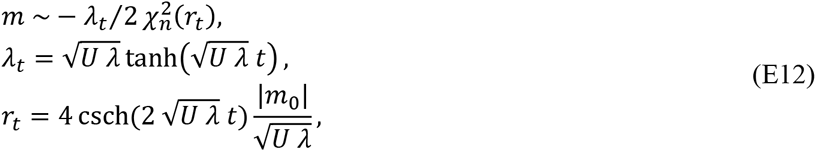

where 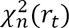 is a non-central chi-square distribution with *n* degrees of freedom and non-centrality parameter *r*_*t*_. As for equilibrium, a corresponding trait distribution can also be derived: we retrieve again a multivariate normal trait distribution, but with time-varying mean and variance. Consider a given position **g**_0_ of the initial clone in phenotype space, in any direction but with norm satisfying ‖**g**_0_‖ = 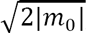; the trait distribution at time *t* is

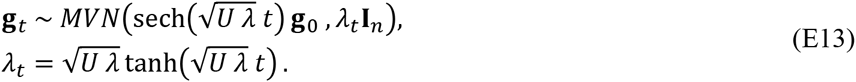

#### Characteristic time of the trajectory

Let us consider the time *t*_*q*_ it takes to fulfill a proportion *q* of the full fitness trajectory. This *t*_*q*_ is the time at which 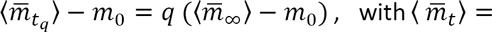 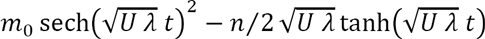. When *m*_0_ = 0 it yields the time to reach equilibrium, from an optimal clonal population. This time is 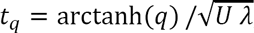 for example with *q* = 0.99, it is *t*_*q*_ ≈ 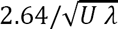. When considering adaptation from a strongly suboptimal clone 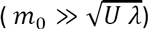, most of the trajectory is driven by the term 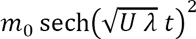, and the characteristic time is then 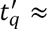 arcsech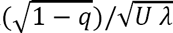. Again with *q* = 0.99 it is 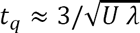. The two characteristic times are remarkably close; in general, their ratio is 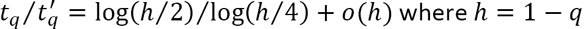, which remains very close to 1 for any small *h* (large *q*). Therefore, it takes roughly the same time 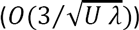 for an optimal clone to reach mutation selection balance, and for a suboptimal clone to reach the vicinity of the optimum. The characteristic time is independent of the initial distance and only proportional to 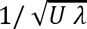.

#### Adaptation from a population formerly at equilibrium

Alternatively, the population may be initially at equilibrium, around some optimum in phenotype space **O**_0_. At time *t* = 0 the optimum shifts from g_0_ to the origin **O**_1_ = **0** of the landscape, due to a change in the environment, and the population adapts to it. As the equilibrium is characterized by a normal distribution of phenotypes in the diffusion regime, we may import the theory of the Gaussian FGM to compute the initial fitness distribution after the optimum has shifted. We have 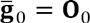 the initial position of the mean phenotype (the former optimum), corresponding to some fitness lag (in the new environment) 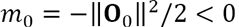, in whatever direction. The initial trait distribution is Multivariate Normal: 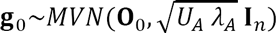 corresponding to the ancestral (‘A’) mutation rate *U*_*A*_ and phenotypic variance *λ*_*A*_ that were affecting the population before the environmental change. In the new environment, the mutation rate and phenotypic variance are *U* and *λ*, respectively, yielding a new 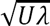. The stochastic representation of fitness, at the onset of the environmental change, is 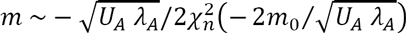, and the CGF of the fitness distribution is 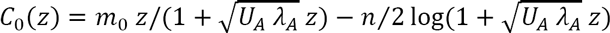. Let 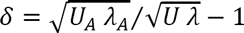 be the relative change in 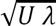 before and after the onset of stress. Plugging *C*_0_(.) into our general solution yields a form similar to Eq. **(E11)**:

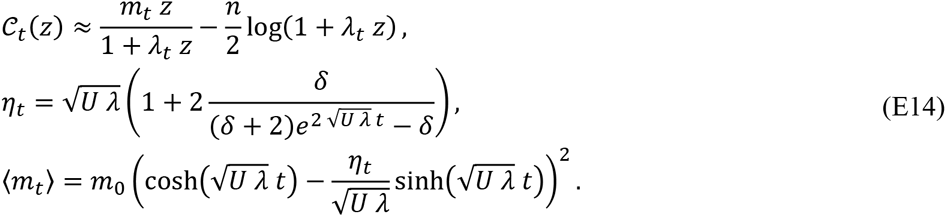

For a small effect of the environmental change on 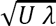 (to leading order in *δ*), we have 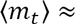 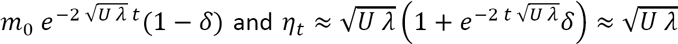 after some time. With no change in either *U* or *λ* across environments (*δ* = 0), the expected mean fitness trajectory is simply 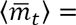 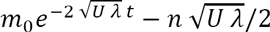. The stochastic representation of the fitness distribution is

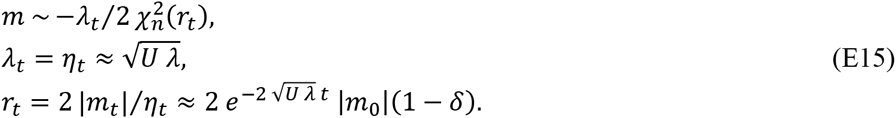

In similarity with the model starting from a clonal population, the corresponding trait distribution is again Gaussian (with approximately constant variance when *δη* ≪ 1):

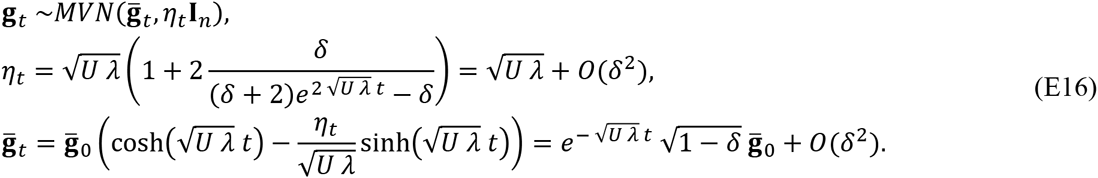

The qualitative behavior of the trait distribution in the case of a constant mutation rate and effects (*δ* = 0) has been pointed out previously (Hereford *et al*. 2004): from equilibrium, the trait distribution evolves as a Gaussian traveling wave with constant variance; the mean distance from the optimum decreases exponentially as 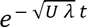. The effect of mild changes in U or *λ* between environments is approximately to modify the effective distance to the optimum, by a factor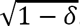.

#### Characteristic time of the trajectory

As above, let us consider the characteristic time for this trajectory. When *m*_0_ ≈ 0 the interesting situation arises when *U*_*A*_ *≠ U* or λ_*A*_ *≠ λ* (otherwise the system stays at the same equilibrium). The characteristic time then describes how long it takes to adjust to a new mutation rate or mutational variance, without moving from the optimum. Taking a leading order *δ* yields 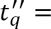 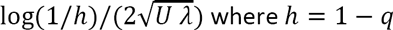 where *h* = 1 – *q*. The time to adjust is independent of the difference in mutation rates, as long as they are close (*δ* ≪ 1). When 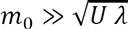, away from the optimum, and if we consider this time that *δ* = 0 for simplicity, the trajectory is driven by the term in 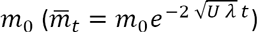 and we obtain again 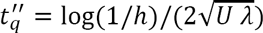. It takes roughly the same time to adjust between different mutation rates at equilibrium and to adapt to a new environment, from equilibrium.

The characteristic time for adaptation to a new environment, from standing variance or from an initially clonal population are of similar order: 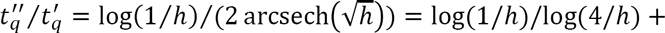 *o*(*h*) remains within [0.6,1] with *h* ∈ [0,0.1]. In this regime of strong mutation weak selection, *de novo* mutation drives the dynamics.

#### PDF of the fitness distribution over time

As it appears, both models yield the same form of stochastic representation; it is thus useful to derive its corresponding pdf. The distributions are of the form *m* ~ − 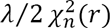 with some *λ* and *r* (which, here, depend on time), given by Eqs. **(E12)** and **(E15)**, depending on the model. The pdf of the distribution is (from that of the non-central chi-square):

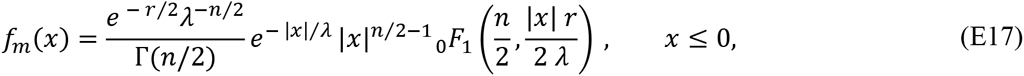

where _0_*F*_1_(.,.) is the confluent hypergeometric function and |*x| = −x* is the absolute value of *x*.

## References

Barton, N. H., 1998 The geometry of adaptation. Nature 395: 751–752.

Brunet, E., I. M. Rouzine and C. O. Wilke, 2008 The stochastic edge in adaptive evolution. Genetics 179: 603–620.

Burger, R., 1991 moments, cumulants, and polygenic dynamics. Journal of Mathematical Biology 30: 199–213.

Burger, R., 1998 Mathematical properties of mutation-selection models. Genetica 103: 279–298.

Burger, R., 2000 The Mathematical theory of selection, mutation, recombination. John Wiley & Sons, Chichester, UK

Burger, R., and J. Hofbauer, 1994 Mutation Load and Mutation-Selection-Balance in Quantitative Genetic-Traits. Journal of Mathematical Biology 32: 193–218.

Chou, H. H., H. C. Chiu, N. F. Delaney, D. Segre and C. J. Marx, 2011 Diminishing Returns Epistasis Among Beneficial Mutations Decelerates Adaptation. Science 332: 1190–1192.

Desai, M., M., 2013 Statistical questions in experimental evolution. Journal of Statistical Mechanics: Theory and Experiment 2013: P01003.

Desai, M. M., and D. S. Fisher, 2007 Beneficial mutation-selection balance and the effect of linkage on positive selection. Genetics 176: 1759–1798.

Desai, M. M., and D. S. Fisher, 2011 The Balance Between Mutators and Nonmutators in Asexual Populations. Genetics 188: 997–1014.

Desai, M. M., D. S. Fisher and A. W. Murray, 2007 The speed of evolution and maintenance of variation in asexual populations. Current Biology 17: 385–394.

Dwyer, J. P., 2012 The dynamics of adapting, unregulated populations and a modified fundamental theorem. Journal of The Royal Society Interface 10.

Eigen, M., 1971 Self-organization of matter and evolution of biological macromolecules. Naturwissenschaften 58: 465–523.

Fisher, R. A., 1930 The genetical theory of natural selection. Oxford University Press, Oxford.

Frank, S. A., 2014 Generative models versus underlying symmetries to explain biological pattern. Journal of Evolutionary Biology 27: 1172–1178.

Gerrish, P., 2001 The rhythm of microbial adaptation. Nature 413: 299–302.

Gerrish, P. J., and R. E. Lenski, 1998 The fate of competing beneficial mutations in an asexual population. Genetica 103: 127–144.

Gerrish, P. J., and P. D. Sniegowski, 2012 Real time forecasting of near-future evolution. Journal of the Royal Society Interface 9: 2268–2278.

Good, B. H., and M. M. Desai, 2013 Fluctuations in fitness distributions and the effects of weak linked selection on sequence evolution. Theoretical Population Biology 85: 86–102.

Good, B. H., and M. M. Desai, 2014 Deleterious Passengers in Adapting Populations. Genetics 198: 1183–1208.

Good, B. H., and M. M. Desai, 2015 The Impact of Macroscopic Epistasis on Long-Term Evolutionary Dynamics. Genetics 199: 177–190.

Good, B. H., I. M. Rouzine, D. J. Balick, O. Hallatschek and M. M. Desai, 2012 Distribution of fixed beneficial mutations and the rate of adaptation in asexual populations. Proceedings of the National Academy of Sciences 109: 4950–4955.

Good, B. H., A. M. Walczak, R. A. Neher and M. M. Desai, 2014 Genetic Diversity in the Interference Selection Limit. PLoS Genet 10: e1004222.

Gordo, I., and P. R. A. Campos, 2012 Evolution of clonal populations approaching a fitness peak. Biology Letters.

Hallatschek, O., 2011 The noisy edge of traveling waves. Proceedings of the National Academy of Sciences 108: 1783–1787.

Hansen, T. F., 1992 Selection in Asexual Populations - an Extension of the Fundamental Theorem. Journal of Theoretical Biology 155: 537–544.

Hietpas, R. T., C. Bank, J. D. Jensen and D. N. A. Bolon, 2013 shifting fitness landscapes in response to altered environments. Evolution 67: 3512–3522.

Johnson, T., 1999 The approach to mutation-selection balance in an infinite asexual population, and the evolution of mutation rates. Proceedings of the Royal Society of London Series B-Biological Sciences 266: 2389–2397.

Keightley, P. D., and A. Eyre-Walker, 1999 Terumi Mukai and the riddle of deleterious mutation rates. Genetics 153: 515–523.

Khan, A. I., D. M. Dinh, D. Schneider, R. E. Lenski and T. F. Cooper, 2011 Negative Epistasis Between Beneficial Mutations in an Evolving Bacterial Population. Science 332: 1193–1196.

Kimura, M., 1965 A Stochastic Model Concerning Maintenance of Genetic Variability in Quantitative Characters. Proceedings of the National Academy of Sciences of the United States of America 54: 731–736.

Kingman, J. F. C., 1978 Simple-Model for Balance between Selection and Mutation. Journal of Applied Probability 15: 1–12.

Kryazhimskiy, S., D. P. Rice, E. R. Jerison and M. M. Desai, 2014 Global epistasis makes adaptation predictable despite sequence-level stochasticity. Science 344: 1519–1522.

Kryazhimskiy, S., G. Tkacik and J. B. Plotkin, 2009 The dynamics of adaptation on correlated fitness landscapes. Proceedings of the National Academy of Sciences of the United States of America 106: 18638–18643.

Lande, R., 1979 Quantitative genetic analysis of multivariate evolution, applied to brain:body size allometry. Evolution 33: 402–416.

Lande, R., 1980 The Genetic Covariance between Characters Maintained by Pleiotropic Mutations. Genetics 94: 203–215.

Manna, F., R. Gallet, G. Martin and T. Lenormand, 2012 The high-throughput yeast deletion fitness data and the theories of dominance. Journal of Evolutionary Biology 25: 892–903.

Manna, F., G. Martin and T. Lenormand, 2011 Fitness Landscapes: An Alternative Theory for the Dominance of Mutation. Genetics 189: 923–937.

Martin, G., 2014 Fisher’s geometrical model emerges as a property of large integrated phenotypic networks. Genetics 197: 237–255.

Martin, G., S. F. Elena and T. Lenormand, 2007 Distributions of epistasis in microbes fit predictions from a fitness landscape model. Nature Genetics 39: 555–560.

Martin, G., and S. Gandon, 2010 Lethal mutagenesis and evolutionary epidemiology. Philosophical Transactions of the Royal Society B-Biological Sciences 365: 1953–1963.

Martin, G., and T. Lenormand, 2006 A general multivariate extension of Fisher’s geometrical model and the distribution of mutation fitness effects across species. Evolution 60: 893–907.

Mccandlish, D. M., C. L. Epstein and J. B. Plotkin, 2014 THE INEVITABILITY OF UNCONDITIONALLY DELETERIOUS SUBSTITUTIONS DURING ADAPTATION. Evolution 68: 1351–1364.

Miralles, R., A. Moya and S. F. Elena, 2000 Diminishing returns of population size in the rate of RNA virus adaptation. Journal of Virology 74: 3566–3571.

Muller, H. J., 1932 Some genetic aspects of sex. Am. Nat. 66: 118–138.

Neher, R. A., and O. Hallatschek, 2013 Genealogies of rapidly adapting populations. Proceedings of the National Academy of Sciences 110: 437–442.

Orr, H. A., 2000 Adaptation and the cost of complexity. Evolution 54: 13–20.

Orr, H. A., 2005 The genetic theory of adaptation: A brief history. Nature Reviews Genetics 6: 119–127.

Otto, S., and N. Barton, 2001 Selection for recombination in small populations. Evolution 55: 1921–1931.

Perfeito, L., A. Sousa, T. Bataillon and I. Gordo, 2014 Rates of fitness decline and rebound suggest pervasive epistasis. Evolution 68: 150–162.

Poon, A., and S. P. Otto, 2000 Compensating for our load of mutations: Freezing the meltdown of small populations. Evolution 54: 1467–1479.

Rattray, M., and J. L. Shapiro, 2001 Cumulant dynamics of a population under multiplicative selection, mutation, and drift. Theoretical Population Biology 60: 17–31.

Rouzine, I. M., J. Wakeley and J. M. Coffin, 2003 The solitary wave of asexual evolution. Proceedings of the National Academy of Science of the USA 100: 587–592.

Roze, D., and A. Blanckaert, 2014 epistasis, pleiotropy, and the mutation load in sexual and asexual populations. Evolution 68: 137–149.

Sniegowski, P. D., and P. J. Gerrish, 2010 Beneficial mutations and the dynamics of adaptation in asexual populations. Philos Trans R Soc Lond B Biol Sci 365: 1255–1263.

Sousa, A., S. Magalhaes and I. Gordo, 2011 Cost of Antibiotic Resistance and the Geometry of Adaptation. Molecular Biology and Evolution 29: 1417–1428.

Tenaillon, O., 2014 The Utility of Fisher’s Geometric Model in Evolutionary Genetics. Annual Review of Ecology, Evolution, and Systematics 45: 179–201.

Tenaillon, O., O. K. Silander, J. Uzan and L. Chao, 2007 Quantifying Organismal Complexity using a Population Genetic Approach. Plos One 2: e217. doi:210.1371/journal.pone.000021.

Trindade, S., L. Perfeito and I. Gordo, 2010 Rate and effects of spontaneous mutations that affect fitness in mutator Escherichia coli. Philosophical Transactions of the Royal Society B-Biological Sciences 365: 1177–1186.

Trindade, S., A. Sousa and I. Gordo, 2012 antibiotic resistance and stress in the light of fisher’s model. Evolution 66: 3815–3824.

Tsimring, L. S., H. Levine and D. A. Kessler, 1996 RNA virus evolution via a fitness-space model. Physical Review Letters 76: 4440–4443.

Turelli, M., 1984 Heritable Genetic-Variation Via Mutation Selection Balance - Lerch Zeta Meets the Abdominal Bristle. Theoretical Population Biology 25: 138–193.

Waxman, D., and J. R. Peck, 1998 Pleiotropy and the preservation of perfection. Science 279: 1210–1213.

Waxman, D., and J. R. Peck, 2006 The frequency of the perfect genotype in a population subject to pleiotropic mutation. Theoretical Population Biology 69: 409–418.

Wilke, C. O., 2005 Quasispecies theory in the context of population genetics. BMC Evol Biol 5: 44.

## References

Alfaro, M., and R. Carles, 2014 Explicit solutions for replicator-mutator equations: extinction vs. acceleration. SIAM Journal on Applied Mathematics 74: 1919–1934.

Ewens, W. J., 2004 Mathematical Population Genetics. I. Theoretical Introduction, 2nd Edition. Springer.

Fisher, R. A., 1930 The genetical theory of natural selection. Oxford University Press, Oxford.

Øksendal, B. K., 2003 Stochastic differential equations: an introduction with applications. Springer, Berlin;.

## References

Fogle, C. A., J. L. Nagle and M. M. Desai, 2008 Clonal Interference, Multiple Mutations and Adaptation in Large Asexual Populations, pp. 2163–2173.

Haigh, J., 1978 The accumulation of deleterious genes in a population—Muller’s Ratchet. Theoretical Population Biology 14: 251–267.

Kimura, M., and T. Maruyama, 1966 Mutational Load with Epistatic Gene Interactions in Fitness. Genetics 54: 1337–1351.

Knight, J. L., and S. E. Satchell, 1997 The cumulant generating function estimation method - Implementation and asymptotic efficiency. Econometric Theory 13: 170–184.

Orr, H. A., 2000 The rate of adaptation in asexuals. Genetics 155: 961–968.

## References

Hereford, J., T. F. Hansen and D. Houle, 2004 Comparing strengths of directional selection: How strong is strong? Evolution 58: 2133–2143.

Martin, G., and T. Lenormand, 2015 The fitness effect of mutations across environments: Fisher’s geometrical model with multiple optima. Evolution 69: 1433–1447.

